# Striatal interneuron microcircuits gate reinforcement to stabilize adaptive choice

**DOI:** 10.64898/2026.09.21.753300

**Authors:** Evan A. Iliakis, Jonibek M. Muhsinov, Alexandra N. Ramirez, Francesco Rinaldi, Jamie Galanaugh, Saurabh Pandey, Iván Linares-García, Aaron R. Tachau, Eric Z. Song, Luigim Vargas, Kyuhyun Choi, Joel T. Woolley, Sarah M. Ferrigno, Edgar Díaz-Hernández, Elizabeth N. Holly, David J. Margolis, Eugenio Piasini, Marc V. Fuccillo

**Author notes:** These authors contributed equally (Jonibek M. Muhsinov, Alexandra N. Ramirez).

## Abstract

The dorsomedial striatum guides learning and adaptive decision-making through excitatory synaptic control of its spiny projection neuron outputs. However, the contributions of local inhibitory microcircuitry remain poorly understood. Here, we identify an interneuron circuit in the dorsomedial striatum that links outcome processing to adaptive action selection. During probabilistic push-pull reversal learning, interneurons represented immediate outcomes: somatostatin interneurons were recruited on unrewarded trials and unexpected rewarded trials, whereas tyrosine hydroxylase interneurons were suppressed on unrewarded trials and recruited on rewarded trials. In vivo recruitment of tyrosine hydroxylase interneurons suppressed somatostatin interneuron activity and increased activity in both direct- and indirect-pathway striatal projection neurons, revealing a polysynaptic disinhibitory microcircuit. Transient inhibition of somatostatin interneurons in this pull-tuned region produced a sustained increase in aberrant pull choices and occupancy of a suboptimal pull-preferring behavioral state, whereas inhibition of tyrosine hydroxylase interneurons produced a sustained impairment of pull reinforcement. Longer-term policy changes following somatostatin interneuron inhibition coincided with postsynaptic potentiation of excitatory synapses onto striatal projection neurons, suggesting a potential substrate for the persistence of altered behavioral policies. Together, these findings identify a disinhibitory striatal circuit motif gating reinforcement which transforms individual trial outcomes into temporally broader policy.

**Highlights:**

- Local striatal interneurons transform outcome signals into action-specific reinforcement.
- Tyrosine hydroxylase interneurons disinhibit striatal projection neurons through somatostatin interneurons.
- Transient interneuron inhibition drives persistent, opponent shifts in action policy.
- Somatostatin interneuron inhibition couples altered policy to excitatory synaptic potentiation.

## Introduction

Optimal decision-making requires neural circuits to integrate information about actions and their outcomes and use this information to guide future behavior. The dorsomedial striatum (DMS) is a critical hub for this process, integrating brain-wide cortical, thalamic, and neuromodulatory inputs carrying information about actions, outcomes, and behavioral state to guide choice based on value.^1,2^ Within the DMS, spiny projection neurons (SPNs) constitute the principal output population and are thought to integrate excitatory inputs via plasticity at their excitatory synapses, thereby mediating action selection and learning.^2–5^ However, SPNs are embedded within a sparse but powerful local inhibitory network whose contribution to these computations remains less well understood.^6,7^ Across cortex, hippocampus, and amygdala, specialized interneuron populations regulate complementary aspects of neural processing via control of spike timing, dendritic integration, synaptic plasticity, and population dynamics.^8–12^ How striatal interneurons regulate SPN activity and dendritic plasticity to generate adaptive behavior is unclear.

The molecular identities, intrinsic physiology, and connectivity of striatal interneurons have been resolved with increasing precision by recent anatomical, electrophysiological, and transcriptomic studies,^6,7,13–18^ but this cellular detail has outpaced our understanding of their contributions to striatal-associated behaviors. Soma-targeting parvalbumin-expressing (PV) interneurons exert powerful control over SPN spike timing and coordinated ensemble activity and have been implicated in multiple forms of learned behavior.^19–27^ In contrast, dendritic-targeting somatostatin-expressing (SST) interneurons regulate early motor learning and dopamine release.^28–32^ A third, striatum-specific population of nondopaminergic tyrosine hydroxylase-expressing (TH) interneurons remains less well characterized,^33,34^ although emerging work potentially implicates these cells in goal-directed behavior, sensorimotor gating, and delay-based choice.^35–37^

Despite progress, we lack a unified account of how distinct striatal interneuron populations contribute to adaptive choice. It remains unclear whether they carry redundant or complementary information about actions and outcomes, how their local connectivity transforms these signals into SPN activity, and whether their influence is restricted to ongoing computation or instead produces lasting changes in behavioral policy. SST interneurons are particularly well positioned to regulate learning through their connectivity to SPN dendrites, while the connectivity of TH interneurons raises the possibility that they recruit a striatal disinhibitory motif analogous to interneuron-selective circuits elsewhere in the brain. Resolving these questions requires examining interneuron encoding, circuit organization, causal function, and downstream plasticity within a common behavioral framework.

Here, we examined SST, TH, and PV interneurons in the DMS during probabilistic reversal learning, in which mice integrated recent outcomes to guide action selection. These interneuron populations carried complementary representations of immediate outcomes: SST interneurons were recruited on unrewarded trials and unexpected rewarded trials, whereas TH interneurons were suppressed on unrewarded trials and recruited on rewarded trials. Activating TH interneurons inhibited SST interneurons while disinhibiting both direct- and indirect-pathway SPNs, revealing a polysynaptic disinhibitory microcircuit. Interneurons and SPNs also exhibited a shared preference for pull actions, identifying a pull-selective functional bias within the recorded DMS territory. Brief outcome period inhibition of SST and TH interneurons produced opposing, action-specific changes in reinforcement that persisted across sessions. SST inhibition promoted aberrant pull choice and increased occupancy of a suboptimal pull-preferring behavioral state, whereas TH inhibition selectively impaired reinforcement of pull actions. Longer-term policy changes following SST interneuron inhibition coincided with postsynaptic potentiation of excitatory synapses onto SPNs, suggesting a potential substrate for the persistence of altered behavioral policies. Together, these findings support a model in which SST interneurons constrain reinforcement through dendritic inhibition, whereas TH interneurons transiently relieve this constraint through local disinhibition. By coupling outcome signals to SPN activity and plasticity, this microcircuit stabilizes adaptive choice across experience.

## Results

### A value-based choice task to study dorsomedial striatal interneuron function

The dorsomedial striatum is a hub that integrates cortical, thalamic, and midbrain dopamine inputs to guide value-based choice.^1,38,39^ While contributions of striatal direct and indirect pathways,^3–5^ dopamine,^40–42^ and acetylcholine^43–46^ have been extensively studied,^1,2^ there is little known about the contributions of local GABAergic interneurons, an important omission given the essential roles these circuit elements play in other brain regions.^8–12^

To elucidate differential contributions of dorsomedial striatal interneuron subtypes to value-based choice, we used a probabilistic reversal-learning task that we have previously shown to provide robust value-dependent reporting of choice and motor execution.^47^ Head-fixed mice chose between pushing and pulling a joystick (Fig 1A,C) to obtain sucrose reward. The reward probabilities associated with the two actions (80% versus 20%) reversed in an uncued, blockwise fashion after at least 17 rewarded choices of the higher-probability action (Fig 1B). The probabilistic task structure required mice to integrate outcomes across multiple trials to guide subsequent choices.

**Figure 1.**
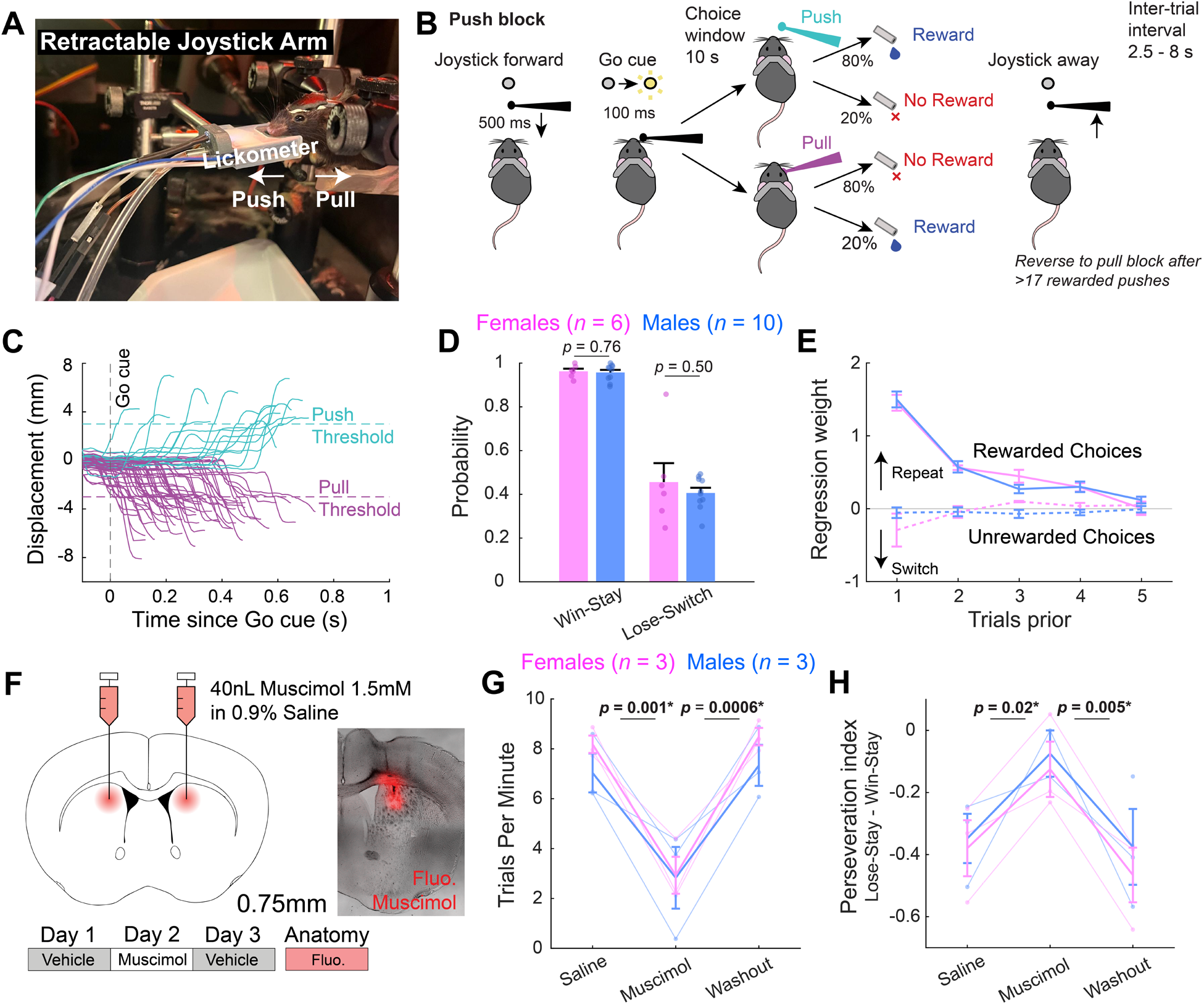
The dorsomedial striatum is required for flexible value-based choice in a probabilistic joystick task. (**A**) Head-fixed behavioral setup in which mice push or pull a joystick to obtain an 8 μL 10% sucrose reward. (**B**) At trial onset, joystick advances toward the mouse (this takes 500 ms), and a Go cue light initiates a 10 s choice window. In a Push block, a push yields reward 80% of the time, whereas a pull yields reward 20% of the time. Reward contingencies reverse after 17 or more rewarded pushes (geometrically distributed, *p* = 0.4). After choice, joystick retracts, starting 2.5 – 8 s inter-trial interval (exponential distribution). (**C**) Representative traces showing push and pull choices crossing the 3 mm displacement threshold. (**D**) Mice (N = 16, 6 females, 10 males) repeated rewarded choices (Win-Stay) and switched following unrewarded choices (Lose-Switch). There were no statistically significant sex differences (linear mixed-effects model, Sex × Metric interaction: *F*(1,28) = 0.398, *p* = 0.533; main effect of Sex: *F*(1,28) = 1.014, *p* = 0.323). (**E**) Logistic regression showed that rewarded choices up to four trials prior positively predicted repetition of the same action on the current trial, whereas unrewarded choices exerted a more modest, single-trial effect. Aside from unrewarded trials three trials prior (*p* = 0.014), there were no statistically significant sex differences (*p*s > 0.05). (**F**) Bilateral injections of the GABA-A receptor agonist muscimol (40 nL/hemisphere, 1.5 mM) into the dorsomedial striatum were performed on Day 2 to inhibit this region, flanked by saline injections on Days 1 and 3 as controls. Post hoc injection of fluorescent muscimol was used to assess targeting. Muscimol reversibly reduced trials per minute (**G**) and increased the perseveration index (**H**, P(lose-stay) – P(win-stay)).

As commonly observed in such tasks,^3,4,48^ mice (N = 16, 6 females, 10 males) preferentially repeated rewarded actions (Win-Stay) and switched at chance levels following unrewarded actions (Lose-Switch; Fig 1D). Logistic regression analysis^49–51^ further showed that rewarded actions up to four trials in the past positively predicted repetition of the same action, whereas the influence of unrewarded actions was weaker and largely limited to the immediately preceding trial (Fig 1E). This finding was consistent with integration of evidence across trials. We detected no major sex differences in either behavioral strategy.

To confirm whether dorsomedial striatal activity supports performance of this task, we reversibly inhibited the region bilaterally using the GABA-A receptor agonist muscimol (Fig 1F; targeting in Fig S1a). Muscimol reversibly reduced task engagement (trials per minute; Fig 1G), and impaired choice flexibility, measured using a perseveration index [*p*(Lose-Stay) – *p*(Win-Stay); Fig 1H]. Muscimol also reduced lose-switch behavior, increased choice latency and omissions, and decreased the number of completed blocks (Fig S1d,h-j). We detected no sex differences in these effects, and no other behavioral measures were significantly altered (Fig S1b,c,e-g). Muscimol produced no detectable changes in most measured movement kinematics (Fig S2), suggesting that its effects on task engagement and choice flexibility are unlikely to arise from a broad impairment in movement execution. Together, these findings establish that our task recruits dorsomedial striatal processes supporting flexible value-based choice and provides a common behavioral framework in which to compare contributions of distinct striatal interneuron subtypes.

### Distinct striatal interneuron populations represent immediate outcomes and recent outcome history

In other brain regions, interneuron subtypes make distinct contributions to behavior by carrying specialized task-related signals and engaging divergent local circuit targets. We therefore asked whether dorsomedial striatal parvalbumin (PV), somatostatin (SST), and tyrosine hydroxylase (TH) interneurons differentially represented actions and outcomes during probabilistic reversal learning. To address this question, we recorded DMS PV, SST, and TH interneuron population activity during our probabilistic reversal task using fiber photometry of jGCaMP8m^53^ (Fig 2A,B; targeting in Fig S3c-e).

**Figure 2.**
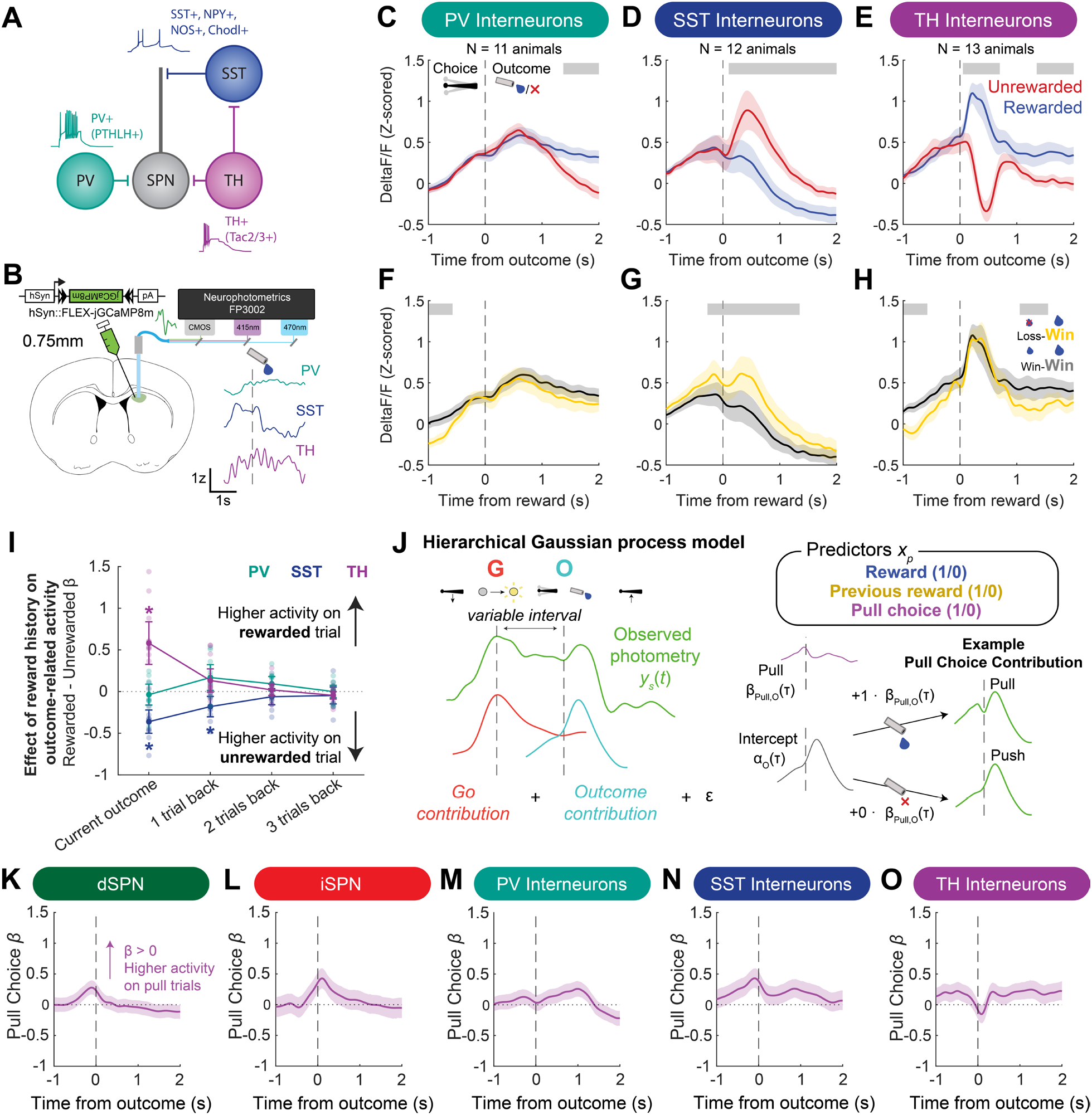
Dorsomedial striatal interneurons differentially reflect outcomes, recent history, and actions. (**A**) Schematic of the dorsomedial striatal microcircuit. Spiny projection neurons (SPNs) are embedded within a local GABAergic microcircuit comprising parvalbumin (PV) interneurons, somatostatin (SST) interneurons, and tyrosine hydroxylase (TH) interneurons. PV interneurons express parathyroid hormone-like hormone (PTHLH), SST interneurons express neuropeptide Y (NPY), nitric oxide synthase (NOS), and chondrolectin (Chodl), and TH interneurons express tachykinin-2/3 (Tac2/3).^13,16^ Representative responses to depolarizing current injection are shown for PV, SST, and TH interneurons, adapted from Kawaguchi^52^ and Ibáñez-Sandoval et al.^33^. (**B**) jGCaMP8m was expressed in dorsomedial striatal SST, PV, or TH interneurons, and population activity was recorded during task performance using fiber photometry. (**C**-**E**) Outcome-aligned population activity in (**C**) PV (N = 11 animals; 5 females, 6 males), (**D**) SST (N = 12 animals; 6 females, 6 males), and (**E**) TH (N = 13 animals; 5 females, 8 males) interneurons on rewarded and unrewarded trials. (**F**-**H**) Outcome-aligned activity on rewarded trials grouped by the outcome of the immediately preceding trial. Loss-win and win-win denote unrewarded ◊ rewarded and rewarded ◊ rewarded outcome sequences, respectively. Grey bars indicate time periods with significant differences identified by cluster-based permutation testing. (I) Associations between current and preceding trial outcomes and outcome-period interneuron activity. For each animal, mean activity during the first second following outcome was modeled as activity ∼ current_outcome + outcome_t-1 + outcome_t-2 + outcome_t-3 + (1 | animalID). Positive coefficients indicate greater activity following rewarded than unrewarded trials; negative coefficients indicate greater activity following unrewarded trials. SST interneuron activity was associated with both current and immediately preceding outcome, whereas TH interneuron activity was associated primarily with current outcome. (**J**) Hierarchical Gaussian process model (HGPM) used to characterize time-varying predictors of photometry activity, following Rinaldi et al. (in preparation). Trial-wise photometry signals were modeled as the sum of Go cue- and Outcome-aligned contributions plus residual error. Predictors included current-trial reward (1, rewarded; 0, unrewarded), previous-trial reward (1, rewarded; 0, unrewarded), and choice (1, pull; 0, push). The example illustrates how the Outcome-aligned intercept and Pull coefficient combine to predict activity on pull and push trials when the remaining predictors equal zero. Positive Pull coefficients indicate greater predicted activity on pull than push trials, conditional on the other predictors. (**K**-**O**) Outcome-aligned population-level Pull coefficient functions for (**K**) direct-pathway spiny projection neurons (dSPNs), (**L**) indirect-pathway spiny projection neurons (iSPNs), (**M**) PV interneurons, (**N**) SST interneurons, (**O**) TH interneurons. Solid lines show posterior estimates, and shaded regions indicate 95% credible intervals.

The three interneuron subtypes exhibited complementary outcome-related dynamics. Dendritic-targeting SST interneurons were preferentially recruited on unrewarded trials (Fig 2D). In contrast, SPN- and SST-targeting TH interneurons showed an opposing bidirectional response, with transient suppression on unrewarded trials and phasic recruitment on rewarded trials (Fig 2E). Interestingly, soma-targeting PV interneurons represented rewarded trials with a slower decrease in activity beginning approximately 1 s following reward (Fig 2C), showing that outcome was represented in the microcircuit across different timescales. This reward-related increase in average population activity of PV and TH interneurons was maintained into the start of the subsequent trial (Fig S4d-f).

Previous work has shown that somatostatin (SST) interneurons are phasically recruited at reward early in the learning of an instrumental lever pressing task, and that this recruitment decreases as animals learn the task and reward is less unexpected.^29^ This finding led us to predict that SST interneurons could be recruited in the setting of unexpected rewards in our probabilistic reversal task. Indeed, SST interneurons showed greater phasic recruitment during rewards following an unrewarded trial (local uncertainty) than during rewards following a rewarded trial (Fig. 2G). In contrast, neither PV nor TH interneurons showed phasic reward responses that depended detectably on the preceding outcome (Fig 2F,H). An expanded analysis of outcome history on current trial signals showed that SST activity during one second following outcome was significantly associated with both current- and previous-trial outcomes, whereas TH activity was associated only with the current-trial outcome (Fig. 2I). Within this post-outcome window, neither current-nor previous-trial outcomes significantly explained PV activity.

To determine whether SST outcome signals were merely dependent on the immediately preceding outcome or instead reflected a temporally broader value estimate, we asked whether reinforcement-learning-derived predictors from a Q-learning model (Fig S5) improved performance of a linear mixed effects model of photometry activity. In SST interneurons, adding either signed or unsigned reward prediction error (RPE) modestly improved held-out prediction relative to a reduced model containing current reward and choice. However, neither model-derived predictor improved held-out prediction once previous-trial reward was included (Fig S7a-c). Thus, while THINs faithfully represent current trial outcomes, SST outcome recruitment is dependent on very recent outcome history.

### Striatal population activity preferentially represents pull actions within a spatially localized DMS territory

We next sought to analyze interneuron choice representations. Since choices immediately precede outcomes in our decision-making paradigm, outcome-aligned windows can also provide a sense of choice-related activity. However, the advance of the joystick at trial start reliably recruits all of our recorded cell types (Fig S4d-f; Fig S8f,j), making it difficult to disentangle to what extent the pre-outcome signal reflects Go-related recruitment versus choice-related activity.

To disentangle overlapping Go- and outcome-aligned signals and isolate their modulation by choice, we used a Bayesian hierarchical Gaussian process model (HGPM) following Rinaldi et al. (manuscript in preparation). Each trial was modeled as the sum of smooth Go- and outcome-aligned response functions positioned according to the events’ actual timing and modulated by trial-level behavioral predictors (Fig S6). This event-resolved design separated choice-modulated activity from overlapping Go- and outcome-related signals, while its hierarchical structure retained trial-level variability and accounted for repeated observations nested within animals.

The HGPM revealed preferential representation of pull choices in many striatal cell types, including direct-pathway SPNs (dSPNs), indirect-pathway SPNs (iSPNs), and SST interneurons (Fig 2K-O). Pull coefficients peaked slightly before choice in dSPNs, and slightly afterward in iSPNs (Fig 2K,L), suggesting differences in the temporal organization of action-related signals across the two projection-neuron pathways (e.g., Sippy et al.^54^). Notably, these choice representations were not robustly identifiable in the raw neural signals across SPN subtypes (Fig S8g,h), underscoring the utility of our novel modeling approach.

We next utilized the natural variability in our fiber placement to further map action tuning (Fig S9). In dSPNs and TH interneurons, more ventrally targeted fibers were associated with greater push tuning, whereas more dorsally targeted fibers were associated with greater pull tuning (Fig S9c,d). SST interneurons showed a similar overall pattern, although fibers in the dorsal- and medial-most portions of the dorsomedial striatum instead exhibited push-related signals (Fig S9b). Together, these findings suggest that population activity within the sampled DMS region exhibited stronger encoding of pull actions.

Overall, our photometry recordings suggest complementary task encoding across DMS interneuron populations. SST and TH interneurons carried particularly prominent and opposing immediate-outcome signals, while outcome-history effects emerged in distinct temporal windows but were largely restricted to the immediately preceding trial rather than reflecting an integrated action-value signal. Action-related activity was additionally biased toward pull choices within our sampled DMS territory.

### TH interneurons suppress SST activity and disinhibit SPNs

Next, we sought to address whether differential striatal interneuron function could also reflect distinct connectivity within the striatal microcircuit. While it has already been established that PV and SST interneurons respectively exert soma- and dendritic-targeted inhibition of SPNs,^32^ the net effect of TH interneuron recruitment is unclear.

Previous work has established direct inhibitory connections on both SPNs^33^ and SST interneurons.^36^ These findings suggest a disinhibitory role of TH interneurons on SPNs via SST interneurons, a hypothesis further supported by our photometry experiments showing that SST interneuron recruitment on unrewarded trials coincided with phasic decreases in TH interneuron activity (Fig 2H-I).

To investigate TH-SST interactions, we asked whether TH interneuron recruitment was sufficient to suppress SST outcome responses. During task performance, we optogenetically excited TH interneurons at outcome using the excitatory opsin ChrimsonR, while recording jGCaMP8m in SST interneurons using fiber photometry (Fig 3A; see Fig S10a,c for targeting). Whereas TH interneuron excitation only modestly attenuated SST activity on rewarded trials (Fig 3B), it produced a pronounced suppression of the SST response to unrewarded outcomes (Fig 3C).

**Figure 3.**
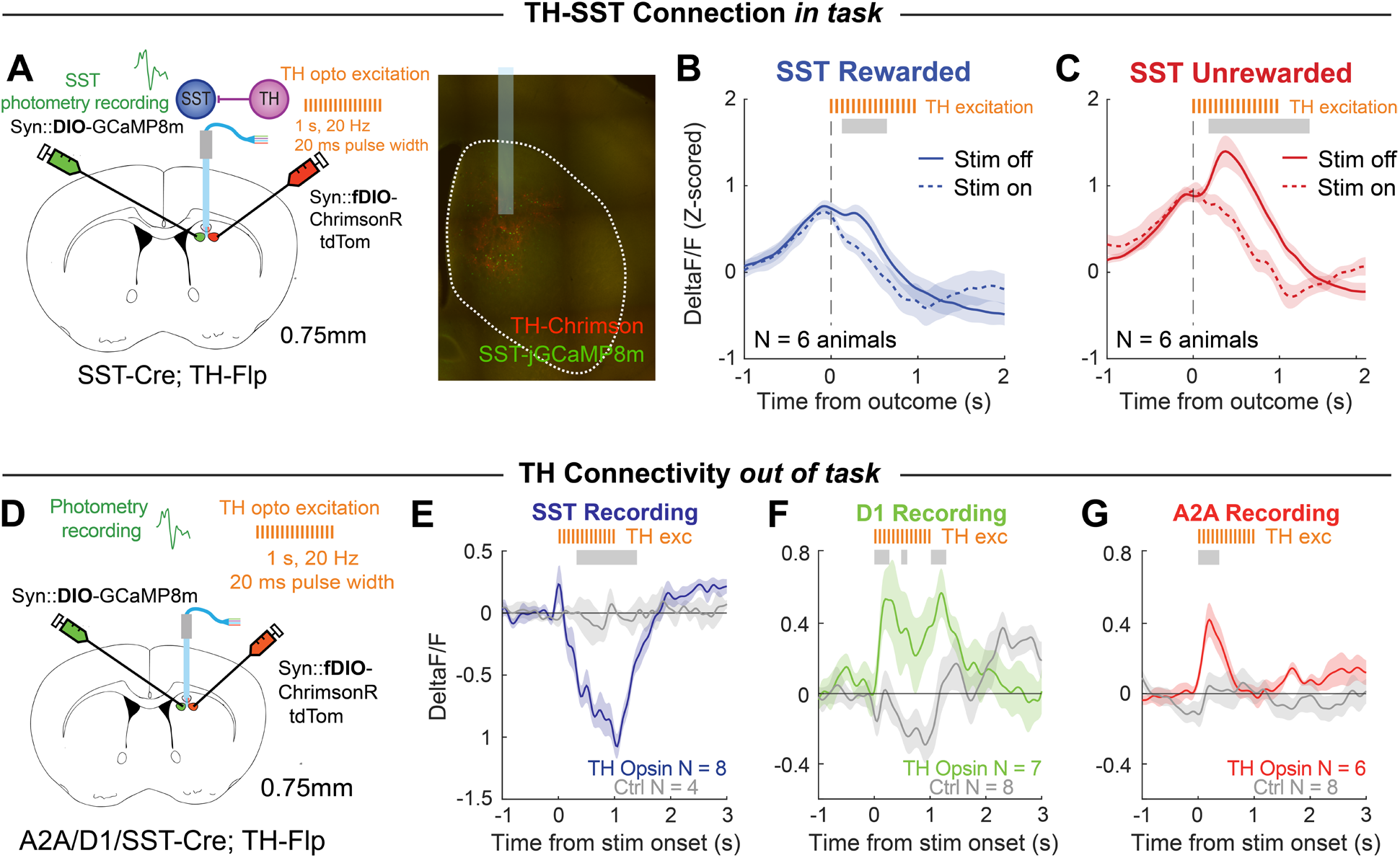
TH interneurons suppress SST encoding of unrewarded outcomes and disinhibit striatal projection neurons. (**A**) Schematic illustrating the recombinase-based strategy used to express the excitatory opsin ChrimsonR in TH interneurons and the calcium sensor GCaMP8m in SST interneurons, enabling simultaneous optogenetic stimulation of TH interneurons and fiber photometry recording of SST interneurons. (**B**) Optogenetic recruitment of TH interneurons during rewarded trials in the probabilistic reversal task modestly attenuates SST reward responses (animal-level mean ± SEM). (**C**) TH interneuron recruitment during unrewarded trials strongly suppresses SST responses to unrewarded outcomes. Stimulation-on and stimulation-off responses were compared using two-sided paired cluster-based permutation tests across time. Grey bars mark the descriptive temporal extent of clusters significant at the cluster level (*p*_cluster_ < 0.05, with family-wise error correction). (**D**) Schematic illustrating the recombinase-based strategy used to express ChrimsonR in TH interneurons and GCaMP8m in dSPNs, iSPNs, or SST interneurons. (**J**) TH interneuron stimulation suppresses SST interneuron activity in TH-opsin animals (*N* = 8 animals) relative to no-opsin controls (*N* = 4 animals). (**E**-**G**) TH interneuron stimulation increases dSPN activity (TH opsin, *N* = 7 animals; control, *N* = 8 animals) and iSPN activity (TH opsin, *N* = 6 animals; control, *N* = 8 animals; animal-level mean ± SEM). Opsin and no-opsin groups were compared using two-sided unpaired cluster-based permutation tests across time. Grey bars mark the descriptive temporal extent of clusters significant at the cluster level (*p*_cluster_ < 0.05, with family-wise error correction).

Given their inhibition of SST interneurons, we next asked if TH interneuron recruitment might lead to net disinhibition of SPNs. To identify the local connections supporting a disinhibitory interaction, we mapped the inhibitory outputs of SST and TH interneurons ex vivo. We optogenetically stimulated either interneuron population and recorded optogenetically evoked inhibitory postsynaptic currents (oIPSCs) from neighboring, pathway-identified SPNs, while separately comparing TH ◊ SST and TH ◊ SPN connectivity (Fig S11). We found that SST interneuron stimulation evoked robust oIPSCs in sequentially recorded neighboring dSPNs and iSPNs, with no significant difference in amplitude between SPN subtypes^32^ (Fig S11a,b). In contrast, we discovered that TH interneuron excitation evoked less consistently detected oIPSCs in sequentially patched dSPNs (6/8 cells) and iSPNs (4/8 cells), again with no pathway difference (Fig S11c).

Given the lower apparent incidence of inhibitory connectivity between TH interneurons and SPNs, we wondered if TH interneurons might primarily connect locally to SST interneurons. To test the relative synaptic strength of TH-SPN and TH-SST interneuron connections, we optogenetically excited TH interneurons while recording from sequentially-patched SST interneurons (labeled green), and unlabeled putative SPNs. TH-evoked oIPSC amplitudes did not differ significantly between neighboring SPNs (11/14 cells) and SST interneurons (8/14 cells; Fig S11d,e).

To assess in vivo effects of TH interneuron activity on pathway-identified SPNs and SST interneurons, we optogenetically excited TH interneurons while recording jGCaMP8m from dSPNs, iSPNs, or SST interneurons using fiber photometry (Fig 3D; see Fig S10a,b,d-g for targeting). Relative to no-opsin light-only controls, TH interneuron excitation significantly reduced SST interneuron activity (Fig 3E), whereas it increased dSPN (Fig 3F) and iSPN (Fig 3G) activity. Together, these findings are consistent with a model in which TH interneurons can exert a net disinhibitory effect on SPNs, potentially mediated through inhibition of SST interneurons.

### Repeated outcome-period inhibition of SST and TH interneurons produces delayed, action-specific changes in choice

Previous optogenetic experiments in the dorsal striatum have identified both immediate trial-by-trial behavioral effects^3,55,56^ and slower changes that accumulate across repeated perturbations.^57–59^ Our photometry recordings showed that SST and TH interneuron signals were primarily related to the current and immediately preceding outcomes, suggesting that these populations could influence the subsequent choice on a trial-by-trial basis. However, SST interneurons also target SPN dendrites and are therefore well positioned to regulate dendritic excitability and synaptic plasticity, which could control behavior across longer timescales. Through their inhibitory connections onto SST interneurons, TH interneurons could indirectly regulate the same dendritic compartment. Therefore outcome-related activity in these populations could also contribute to slower, cumulative changes in behavior.

To simultaneously examine immediate and cumulative effects, we developed an optogenetic protocol in which SST or TH interneurons were inhibited on a randomly selected 33% of trials across six sessions (Fig 4A; see Fig S12c-d for targeting). We limited optogenetic inhibition to 1 second following outcome given the opposing, dynamic, phasic modulation of SST and TH interneurons at this epoch (Fig 4B, see Fig 2). Animals completed two pre-inhibition probabilistic sessions (Pre), six optogenetic sessions divided into Opto I and Opto II, and four post-inhibition sessions divided into Post I and Post II. Comparing choices following light-on and light-off outcomes allowed us to test the immediate effect of individual perturbation, whereas comparisons across experimental phases allowed us to identify behavioral changes that accumulated over repeated optogenetic sessions and persisted after inhibition ended. To ensure optogenetic inhibition was effective, we validated the ability of our inhibitory opsin enhanced halorhodopsin 3.0 (eNpHR3.0) to suppress SST and TH interneuron activity both in vitro and in vivo (Fig S12a,b).

**Figure 4.**
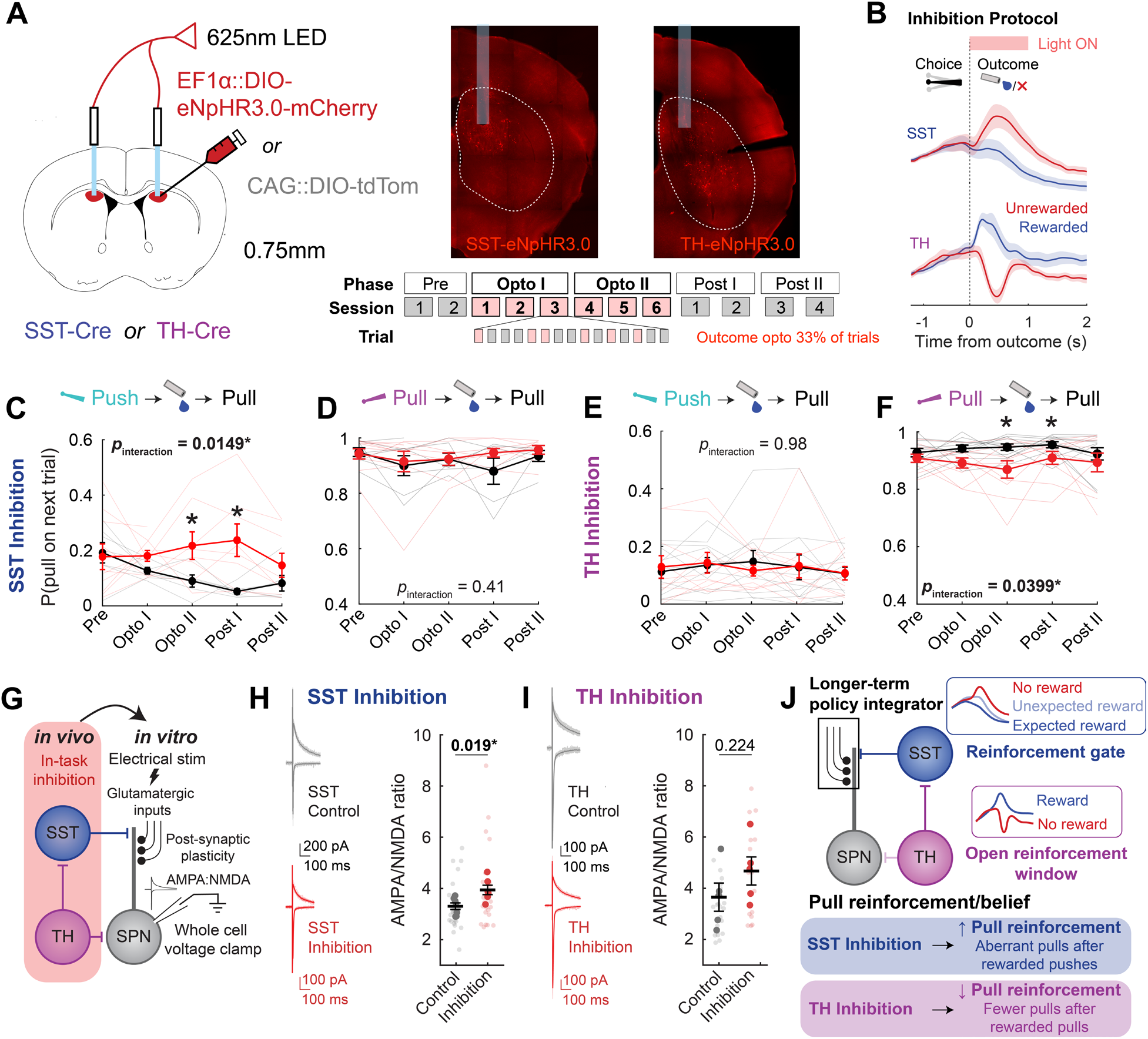
Repeated outcome-period inhibition of dorsomedial striatal SST and TH interneurons produces sustained, action-specific changes in pull behavior, with associated potentiation of excitatory synapses onto SPNs by SST inhibition. **(A)** Viral strategy for Cre-dependent expression of the inhibitory opsin eNpHR3.0 or a control fluorophore in dorsomedial striatal SST or TH interneurons. Animals (SST cohort: control *N* = 7 animals, inhibition *N* = 10 animals; TH cohort: control *N* = 11 animals, inhibition *N* = 10 animals) completed two pre-inhibition probabilistic sessions (Pre), six optogenetic sessions divided between Opto I and Opto II, and four post-inhibition sessions divided between Post I and Post II. (**B**) On 33% of trials during optogenetic sessions, light was delivered for 1 s beginning at outcome, coinciding with the phasic outcome-related activity of SST and TH interneurons observed by fiber photometry (Fig 2). (**C**-**F**) Probability of choosing pull on the subsequent trial following (**C**,**E**) rewarded push or (**D**,**F**) rewarded pull across experimental phases in (**C**-**D**) SST and (**E**-**F**) TH cohorts. Thin lines show individual-animal trajectories; thick lines and points show group means ± SEM. Control and inhibition groups were compared using binomial generalized linear mixed-effects models with phase, condition, and their interaction as fixed effects and animal and session as random intercepts. Significant phase × condition interactions were observed following rewarded pushes in SST cohorts (*p* = 0.0149), and following rewarded pulls in TH cohorts (*p* = 0.0399), but not in the corresponding rewarded-pull SST (*p* = 0.41) or rewarded-push TH (*p* = 0.98) comparisons. Relative to Pre, condition-dependent changes were detected during Opto II (*p* = 0.016) and Post I (*p* = 0.0015) following rewarded pushes in SST cohorts, and during Opto II (*p* = 0.013) and Post I (*p* = 0.043) following rewarded pulls in TH cohorts. (**G**) Strategy for measuring excitatory synaptic strength onto SPNs following the in vivo inhibition protocol. Acute slices were prepared after repeated SST- or TH-interneuron inhibition, and AMPA/NMDA ratios were measured in SPNs using electrical stimulation of glutamatergic inputs and whole-cell voltage-clamp. (**H**) SPN AMPA/NMDA ratios were significantly higher following SST interneuron inhibition (*N* = 6) relative to control animals (*N* = 7; Welch’s *t* test, *p* = 0.019). (**I**) SPN AMPA/NMDA ratios were not significantly altered following TH interneuron inhibition (*N* = 5) relative to controls (*N* = 5; Welch’s t test, *p* = 0.224). (**J**) Proposed model for complementary roles of SST and TH interneurons in reinforcement. We hypothesize that the recorded dorsomedial striatal territory preferentially reinforces pull actions, or represents belief that pull is the higher-value action, with longer-term policy information stored in excitatory synaptic weights onto SPNs. SST interneurons are proposed to act as a reinforcement gate: their recruitment on unrewarded and unexpected rewarded trials constrains excitatory synaptic plasticity and limits reinforcement of pull actions. TH interneurons may transiently open this reinforcement window on rewarded trials by suppressing SST interneurons. Accordingly, SST inhibition would inappropriately open the gate, increasing pull choices following rewarded pushes and potentiating excitatory synapses onto SPNs. Conversely, TH inhibition would reduce reward-associated opening of the gate, weakening reinforcement of rewarded pull actions.

Surprisingly, neither SST nor TH interneuron inhibition produced any detectable acute effect on choice immediately following an illuminated trial, measured using either win-stay or lose-switch metrics (Fig S13a-h). Instead, action-specific differences between inhibition and control groups emerged progressively across sessions. SST inhibition within this pull-preferring region (Fig 2K-O) increased the probability of aberrant pulls following rewarded pushes. This effect emerged during Opto I, grew during Opto II, and was largest during Post I, after optogenetic inhibition had ended (Fig 4C,D). Conversely, TH inhibition reduced the probability of repeating a rewarded pull, an effect that also persisted into Post I (Fig 4E,F). Choice following unrewarded outcomes showed no similarly consistent phase-dependent change (Fig S13i-l). Thus, repeated SST inhibition progressively biased choice toward pull even when push had just been rewarded, whereas repeated TH inhibition weakened the tendency to repeat rewarded pull actions.

These complementary effects support a model in which SST interneurons constrain reinforcement through dendritic inhibition, whereas TH-mediated inhibition of SST interneurons transiently relieves this constraint (Fig. 4J). Within a pull-preferring striatal territory, repeated SST inhibition could therefore produce aberrant strengthening of pull-related actions, whereas TH inhibition could instead prevent the normal reinforcement of pull actions (Fig 4J).

Because the behavioral effect of SST inhibition persisted after optogenetic sessions (Post I), we asked whether repeated inhibition was accompanied by altered excitatory synaptic strength onto SPNs. SST interneurons preferentially target SPN dendrites,^32^ positioning them to regulate dendritic excitability and excitatory synaptic plasticity. We therefore measured AMPA/NMDA (A/N) ratios, a commonly used index of excitatory synaptic plasticity on the post-synapse, in SPNs from acute slices prepared after completion of our in vivo inhibition protocol (Fig 4G). SPNs from SST inhibition animals exhibited significantly higher A/N ratios than those from control animals receiving identical optical illumination (Fig 4H), consistent with potentiation of excitatory inputs onto SPNs. TH inhibition produced a nonsignificant increase in A/N ratio (Fig. 4I). Thus, the persistent behavioral effects of repeated SST inhibition coincided with consistently enhanced excitatory synaptic strength onto SPNs, providing a potential synaptic substrate for the longer-term changes in behavioral policy.

### SST interneuron inhibition increases occupancy of a pull-biased latent decision state

Because the persistent behavioral effects of SST inhibition coincided with increased strength of excitatory synapses onto SPNs, we asked whether the excess pull choices reflected a longer-lasting alteration in decision strategy. Win-stay and lose-switch probabilities describe how the immediately preceding outcome affects choice, but they neither uniquely nor comprehensively identify the complete decision strategy generating value-based behavior. Animals can solve probabilistic tasks through combinations of incremental action-value learning, intrinsic action biases, choice-history dependence, and different exploration-exploitation strategies.

To represent these potential mechanisms within a single choice policy, we developed an RL-GLM in which reinforcement-learning parameters and choice-predictor weights were optimized jointly (see Methods). A central predictor was the difference between the estimated values of the push and pull actions (ΔQ), calculated using a Q-learning model with forgetting. The learning-rate parameter α determined how strongly each outcome updated the corresponding action value. Because interneuron inhibition produced asymmetric effects on push and pull behavior, we separated positive values of ΔQ, which favored pushing, from negative values, which favored pulling. Their respective weights, β+ and β−, quantified the transformation of push- and pull-favoring value evidence on choice, while the remaining predictors captured action bias and recent choice or outcome history.

Although this model represented multiple influences on choice, it continued to assume that a single parameterization governed behavior throughout the session. We therefore embedded the RL-GLM within an HMM, allowing each latent state to have its own learning rate and choice-predictor weights while estimating transitions among states across trials^56,60,61^ (Fig. 5A). Model comparison supported three recurring latent behavioral states: balanced (State 1), push-preferring (State 2), and pull-preferring (State 3). The balanced State 1 accounted for approximately 60% of trials and exhibited the highest choice accuracy (Fig. 5D,I). It showed no overall action bias, a nearly symmetric relationship between ΔQ and choice, and high action-specific win-stay probabilities (Fig. 5E–H). Correspondingly, its fitted policy had a near-zero intercept, similar sensitivity to push- and pull-favoring value evidence, and a strong effect of previous rewarded outcome on choice (Fig. 5B). Its relatively low learning rate (*α* = 0.36; Fig. 5C) was consistent with integration across a broader outcome history.

**Figure 5.**
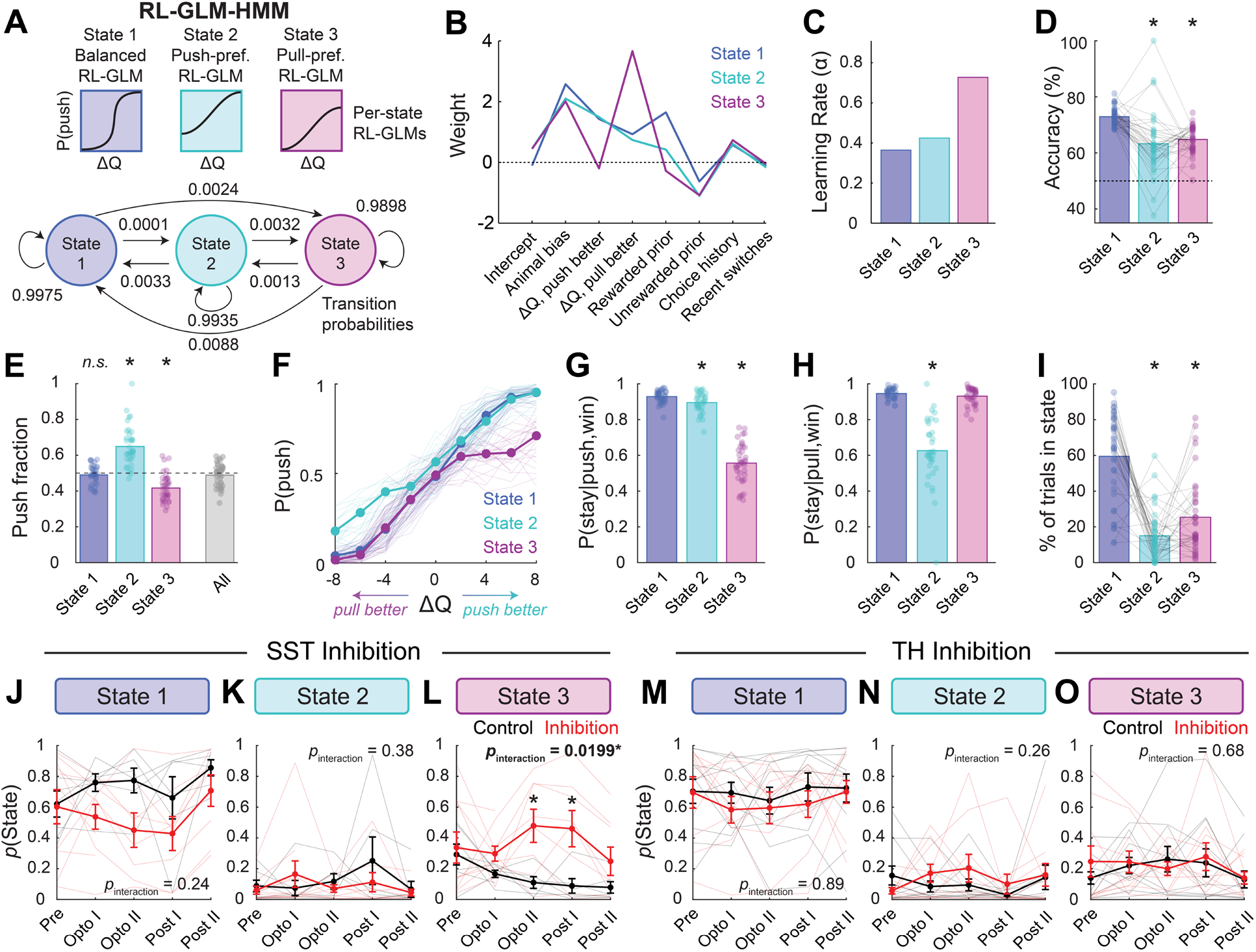
A three-state reinforcement learning–generalized linear model–hidden Markov model (RL-GLM-HMM) identifies sustained changes in latent decision-state occupancy following SST interneuron inhibition. **(A)** Schematic of the RL-GLM-HMM. Each latent state is associated with a unique set of RL-GLM predictor weights and a state-specific learning rate. Transition probabilities from the final three-state model are shown beside the corresponding arrows. **(B)** State-specific predictor weights and **(C)** learning rates from the three-state RL-GLM-HMM fit to all sessions from Pre through Post II across the SST and TH inhibition and control cohorts. **(D)** State-specific accuracy, defined as probability of choosing the action with the higher current reward probability. **(E)** Fraction of push choices within each state and across all trials (All). **(F)** Probability of choosing push as a function of the current-trial push-pull action-value difference (ΔQ) within each state. Transparent lines show individual-animal data. **(G)** Push and **(H)** pull win-stay probabilities within each state. (I) Percentage of trials assigned to each state. In **D, G-I,** asterisks indicate significant differences from State 1; in **E,** asterisks indicate significant differences from All (Wilcoxon signed-rank tests, \**p* < 0.05). (**J-O**) State occupancy probabilities across Pre, Opto I, Opto II, Post I, and Post II for (**J-L**) SST and (**M-O**) TH interneuron cohorts. Trajectories in the inhibition and control cohorts were compared using linear mixed-effects models (stateProbability ∼ phase * condition + (1 | animalID)). A marginal ANOVA test identified a significant phase × condition interaction for State 3 in the SST cohort (*p* = 0.0199). Phase-specific interaction terms were significant at Opto II (*p* = 0.0067) and Post I (*p* = 0.0063).

In contrast, States 2 and 3 exhibited opposing action biases and suboptimal value-based performance (Fig 5D,E). In the push-preferring State 2, mice remained more likely to push even when ΔQ was negative – i.e., the estimated value of pulling was greater (Fig 5F). They were also less likely to repeat rewarded pulls (Fig 5G,H). This state retained sensitivity to push-favoring value evidence but showed reduced sensitivity to pull-favoring evidence, as reflected by a preserved β^+^ but reduced β^−^ relative to the balanced state (Fig 5B).

Conversely, mice in the pull-preferring State 3 remained less likely to push even when ΔQ was positive – i.e., the estimated value of pushing was greater (Fig 5F). They were also less likely to repeat rewarded pushes (Fig 5G,H). State 3 was characterized by a β^+^ near zero indicating markedly reduced sensitivity to value evidence favoring push, together with preserved sensitivity to pull-favoring evidence (Fig 5B). This state also showed a weaker influence of previous outcome and the highest learning rate (α = 0.73; Fig 5B,C) of all states. Both states were less accurate than the ‘optimal’ balanced state and accounted for a smaller proportion of trials (Fig 5D,I). Further details about the features of these three states are outlined in Fig S15.

We next asked whether outcome-associated interneuron inhibition altered occupancy of these latent states across experimental phases. SST interneuron inhibition significantly increased occupancy of the ‘suboptimal’ pull-preferring State 3 relative to controls, and produced a corresponding numerical reduction in balanced State 1 occupancy (Fig 5J-L). In the TH cohort, inhibition was associated with a qualitative, non-significant increase in occupancy of the push-preferring State 2 (Fig 5L-O).

To confirm the principal behavioral signature of the latent-state model without assigning trials to discrete states, we examined push probability continuously as a function of ΔQ across experimental phases (Fig S14a,b). Control and inhibition showed similar choice functions at baseline. During SST interneuron inhibition, however, animals became more likely to pull when ΔQ was positive (i.e., when push was the higher-valued action). This change persisted into Post I and subsequently normalized by Post II (Fig S14a,b), paralleling the increased occupancy of the pull-preferring state. We quantified these changes by separately estimating choice sensitivity to positive and negative values of ΔQ (Fig S14c). SST interneuron inhibition selectively reduced β^+^, indicating diminished sensitivity to push-favoring value evidence and consequently increased pull choice when push had the higher estimated value, while producing no change in β^−^ (Fig S14d). This state-assignment-independent result recapitulated the defining signature of the pull-preferring State 3, in which the β^+^ was similarly near zero. TH interneuron inhibition produced a qualitatively opposite pattern: animals became less likely to pull when ΔQ was negative, consistent with reduced sensitivity to value evidence favoring pull (Fig S14e). Notably, our optogenetic inhibition did not significantly impact biases to one or the other action in the face of neutral or ambiguous value evidence.

Together, these analyses indicate that SST interneuron inhibition did not produce a global action bias or uniform increase in choice stochasticity. Instead, it selectively reduced sensitivity to push-favoring value evidence across sessions, increasing pull choice when push had the higher estimated value and increasing occupancy of the pull-preferring latent behavioral state. This policy shift emerged over repeated inhibition sessions and persisted into the post-stimulation period, recapitulating the delayed time course observed in the conditional-choice analysis. TH inhibition produced directionally complementary but weaker effects on sensitivity to pull-favoring evidence and occupancy of the push-preferring state across sessions.

## Discussion

The dorsomedial striatum guides learning and adaptive decision-making through divergent activity patterns of its spiny projection neurons (SPNs),^1,2^ but how local inhibitory microcircuits regulate this process remains poorly understood. Here, we identify a TH-SST interneuron circuit that couples transient outcome signals to persistent changes in action policy. SST interneurons were recruited primarily by unrewarded outcomes, whereas rewarded outcomes recruited TH interneurons, whose activation suppressed SST activity and produced net disinhibition of both direct- and indirect-pathway SPNs. Within this pull-preferring territory, brief outcome-period inhibition of SST interneurons progressively promoted aberrant pull choices even when push had the higher estimated value, and shifted behavior toward a pull-preferring latent state. Conversely, TH inhibition reduced repetition of rewarded pull actions, consistent with impaired pull reinforcement. The persistent behavioral effects of SST inhibition coincided with increased AMPA/NMDA ratios at excitatory synapses onto SPNs, suggesting a potential mechanism for longer-timescale behavioral effects. Together, these findings support a model in which SST interneurons constrain reinforcement through dendritic inhibition, whereas TH interneurons transiently relieve this constraint through local disinhibition, coupling outcome processing to SPN plasticity and adaptive choice.

### Complementary interneuron representations across outcome valence and timescale

Despite being embedded in the same local network, interneurons can represent distinct features of ongoing behavior.^62,63^ Our examination of three striatal interneuron populations within a common behavioral framework revealed subtype-specific activity patterns across both outcome valence and time. These differences could arise from distinct afferent input patterns, intrinsic physiological properties, or neuromodulatory sensitivity,^6,7,64–66^ although distinguishing among these possibilities will require further study.

These divergent activity patterns suggest a functional division of labor for value-based processing. Recruitment of dendritic-targeting SST interneurons following unrewarded and unexpected rewarded outcomes could constrain dendritic excitability and synaptic plasticity, preventing isolated probabilistic outcomes from disproportionately destabilizing established choice policies. Such regulation would be particularly important in stochastic environments, in which individual outcomes provide unreliable evidence that action– outcome contingencies have changed.^67^ Analogous roles for SST interneurons in regulating dendritic excitability and plasticity have been described throughout the central nervous system.^10,62,68,69^ Importantly, although SST population activity represented only the current and immediately preceding outcomes, repeated disruption of these brief signals produced persistent changes in choice and excitatory synaptic strength. Thus, the limited temporal horizon of an interneuron signal need not constrain the timescale of its behavioral consequences.

In contrast to the unidirectional modulation of SST interneurons, TH interneurons faithfully represented outcomes, being recruited and suppressed by the presence and absence of reward, respectively. Through their inhibition of SST interneurons, TH interneurons could therefore regulate how outcome-valence signals are incorporated into reinforcement across repeated trials. PV interneurons exhibited comparatively little immediate outcome modulation but developed slower reward-related signals that extended into the subsequent trial. Although their causal contribution was not examined here, these delayed PV population signals could alter perisomatic inhibition to regulate SPN gain, coordinated activity, or local oscillations over longer timescales.^8,21,70–72^

### Disinhibitory roles of striatal TH interneurons

Given THIN connectivity to both SPNs^33^ and SST interneurons,^36^ the net circuit effect on SPN output was unclear. Consistent with prior work, our slice recordings revealed comparable TH-evoked inhibition of SPNs and SST interneurons. In vivo, however, TH interneuron recruitment suppressed SST activity while increasing activity in both direct- and indirect-pathway SPNs. Thus, despite forming direct inhibitory connections onto SPNs, TH interneurons exerted a net disinhibitory influence on SPN population activity within the intact circuit.

Several circuit properties could reconcile these findings. The intrinsic physiology of SST interneurons may make their firing particularly sensitive to inhibitory input.^52,73^ Moreover, individual SST interneurons possess extensive axonal arbors that contact SPNs across a broad territory.^32,52,74^ Inhibiting a relatively small number of SST interneurons could therefore release inhibition across a larger SPN population, outweighing the direct inhibition of SPNs by TH interneurons. Interactions with cholinergic interneurons, dopamine axons, and other GABAergic populations could further amplify or reshape this circuit-level response.^7,30,32^

Disinhibitory circuits elsewhere in the brain act as context- and state-dependent gates, transiently releasing principal neurons from inhibition to amplify behaviorally relevant inputs and permit synaptic plasticity.^62,75–77^ Reward-associated TH recruitment may perform an analogous function in the striatum by suppressing SST-mediated dendritic inhibition and increasing SPN excitability. This motif resembles the functional role attributed to VIP interneurons in other regions^62,78^ (although see Dellal et al.^79^). Because CCK/VIP interneurons are exceptionally sparse in mouse dorsal striatum, TH interneurons (and potentially their primate TAC3 homologues) may provide a functionally analogous source of local disinhibition.^13–15,17,18^

Our experiments do not establish that SST interneurons mediate either the net SPN response or the behavioral effects of TH recruitment. Indeed, TH inhibition did not produce a fully reciprocal behavioral phenotype to SST inhibition, potentially reflecting additional TH targets, or distinct consequences of removing endogenous TH-mediated disinhibition versus directly removing SST-mediated inhibition. A prediction of the proposed model is that TH excitation should recapitulate the behavioral effects of SST inhibition.

### Brief interneuron perturbations induce persistent changes in choice policy

Brief striatal perturbations can influence behavior on multiple timescales. When delivered during action selection or execution, temporally precise SPN perturbations can immediately bias lateralized choice, influence the subsequent stay–switch decision, suppress movement, or truncate an ongoing action sequence.^3,55,80–82^ However, brief perturbations can also exert cumulative effects when repeatedly paired with specific actions or behavioral contexts. In these settings, SPN stimulation can reinforce or punish future actions and produce learned changes in movement parameters or context-dependent behavior that persist beyond individual stimulation epochs.^1,57,58,83^ Our findings extend this latter principle to sparse striatal interneuron populations. Although interneuron inhibition was delivered during brief, trial-locked epochs, its behavioral effects emerged across repeated stimulation sessions, generalized to interleaved light-off trials, and remained after stimulation ceased. Interneuron activity may therefore gate the engagement of striatal reinforcement and punishment mechanisms, regulating whether transient action-related activity produces lasting changes in choice policy.

The persistence of these effects also dissociates the timescale of interneuron activity from that of its behavioral consequences. SST and TH activity was aligned to individual actions and outcomes and reflected only recent trial history, whereas perturbing these signals produced changes lasting across many trials and sessions. Such a temporal mismatch is characteristic of an instructive or permissive signal: a brief event can gate plasticity whose consequences are subsequently stored within the circuit.^77,84,85^

One candidate substrate for these persistent effects is plasticity at excitatory inputs onto SPNs. Consistent with this possibility, SPNs exhibited greater AMPAR synaptic currents following our SST-interneuron inhibition protocol. Normal physiological SST interneuron activity may therefore constrain activity-dependent strengthening of corticostriatal or thalamostriatal inputs engaged around outcomes. This function may not be unique to SST interneurons: chronic ablation of PV interneurons similarly potentiates excitatory synapses onto SPNs.^27^ Future work should determine whether repeated PV-interneuron inhibition produces comparable synaptic potentiation, and whether it alters choice policy in the same manner as SST-interneuron inhibition.

Plasticity at excitatory inputs onto SPNs is nevertheless unlikely to be the sole substrate of the persistent behavioral effects. Longer-lasting changes in local dopaminergic, cholinergic, or peptidergic signaling provide additional candidate mechanisms. Notably, SST interneurons also release nitric oxide (NO), neuropeptide Y (NPY), and somatostatin in addition to GABA,^6,86^ raising the possibility that non-GABAergic signaling contributes to the extended timescale of the behavioral effects. Future work should distinguish the contributions of fast synaptic inhibition, plasticity at excitatory inputs, and local neuromodulation to the observed effects of interneurons on behavioral strategy.

### Latent behavioral states reveal selective disruption of value-to-action mapping

Standard implementations of reinforcement-learning models generally assume that a single set of parameters governs behavior throughout a session, whereas GLM-HMM approaches allow animals to transition among persistent latent policies that weight task information differently.^60,61^ By integrating a reinforcement learning model within a GLM-HMM, we could distinguish changes in how values were updated from changes in how learned values were translated into actions.

Our latent-state analysis added two conclusions beyond conditional-choice measures (i.e., win-stay, lose-switch). Notably, SST inhibition did not simply increase overall pull bias or choice stochasticity. Instead, it progressively increased occupancy of a pull-preferring state in which push-favoring value evidence exerted less influence over choice, explaining why animals made aberrant pull choices even when push had higher estimated value. TH inhibition produced directionally complementary but weaker changes in pull-value sensitivity and push-preferring-state occupancy. Thus, interneuron perturbation was expressed principally through the value-to-action transformation rather than through a global impairment in value-guided choice. Although the inferred behavioral states cannot be equated directly with discrete neural states, they may reflect persistent changes in striatal network configuration or, as suggested by our synaptic measurements, changes in excitatory synaptic strength onto SPNs.

### Mechanisms underlying action-specific behavioral effects

The action specificity of our optogenetic effects is not readily explained by outcome signaling alone. SST inhibition progressively promoted pull choices even after rewarded pushes and increased occupancy of a pull-preferring latent state, whereas TH inhibition reduced repetition of rewarded pulls. One explanation is that both manipulations acted upon an anatomically biased action representation. Population activity within the sampled DMS territory preferentially represented pull actions, and placement-dependent variation in dSPN and TH activity further suggested a spatial gradient in push–pull tuning. These observations are consistent with previous evidence that striatal action representations exhibit local spatial organization.^5,87,88^

Action-related excitatory signals entering the striatum may provide an eligibility signal that identifies which synapses and neuronal ensembles should be modified when an outcome occurs.^82,89,90^ Within a pull-preferring territory, repeated SST inhibition could release dendritic inhibition and permit inappropriate strengthening of the locally dominant pull representation, increasing its competition with push-related representations even when push had the higher estimated value. Conversely, TH inhibition could prevent reward-associated suppression of SST interneurons, maintaining dendritic inhibition over pull-related SPNs and weakening reinforcement of rewarded pull actions. Under this model, the opposing behavioral effects arise not from opposite action preferences intrinsic to SST and TH interneurons, but from their sign-opposed control over plasticity within the same spatially biased action representation.

This account remains provisional because our population recordings do not identify the individual SPNs modified by interneuron perturbation, and our synaptic measurements do not establish that potentiation occurred selectively in pull-related ensembles. It nevertheless makes a testable prediction: the direction of the behavioral effect should vary with the action preference of the targeted striatal territory. Perturbing the same interneuron populations in a push-preferring region should therefore produce different, and potentially reversed action-specific effects.

### Limitations and conclusions

One potential limitation in our study is the asymmetry of the required motor responses. Although we found no consistent population-level asymmetry in the early acquisition or skilled execution of push and pull actions, mice transiently preferred pulling when the decision threshold increased early in training, suggesting that pushing might be more motorically demanding. Future experiments could minimize differences in the mechanical requirements of the two responses by using an isometric joystick, which measures directional force without displacement (e.g., Rodrigues-Vaz et al.^91^). A second limitation is that fiber photometry measures population-level signals and therefore cannot resolve functional heterogeneity among individual interneurons. The sparsity of these populations makes single-cell approaches challenging, but opto-tagged electrophysiology (e.g., Duhne et al.^21^) and miniscope imaging could provide essential complementary cellular resolution. Finally, particularly in SPNs, photometry signals may reflect dendritic calcium driven by excitatory input rather than firing rate per se, given recent evidence that SPN photometry signals arise predominantly from non-somatic compartments^92^ (although see Lipton et al.^93^). Nevertheless, increased dendritic excitation in the absence of increased firing would remain consistent with the proposed release of SPNs from dendritic inhibition through TH-interneuron-mediated suppression of SST interneurons.

In sum, our findings provide in vivo functional evidence that striatal TH interneurons suppress SST activity and exert a net disinhibitory influence on SPNs. Within a pull-tuned DMS region, brief outcome-period perturbation of this circuit accumulated into persistent, action-specific changes in choice policy, with SST inhibition accompanied by increased excitatory synaptic strength onto SPNs. These findings expose a central unresolved tension: how are interneuron signals representing only immediate outcome history converted into durable, action-selective changes in synaptic strength and behavior? By linking outcome coding, local disinhibition, and longer-term policy stabilization, our work identifies inhibitory microcircuits as active regulators of how the striatum transforms transient behavioral feedback into adaptive choice.

## STAR Methods

### Key Resources Table

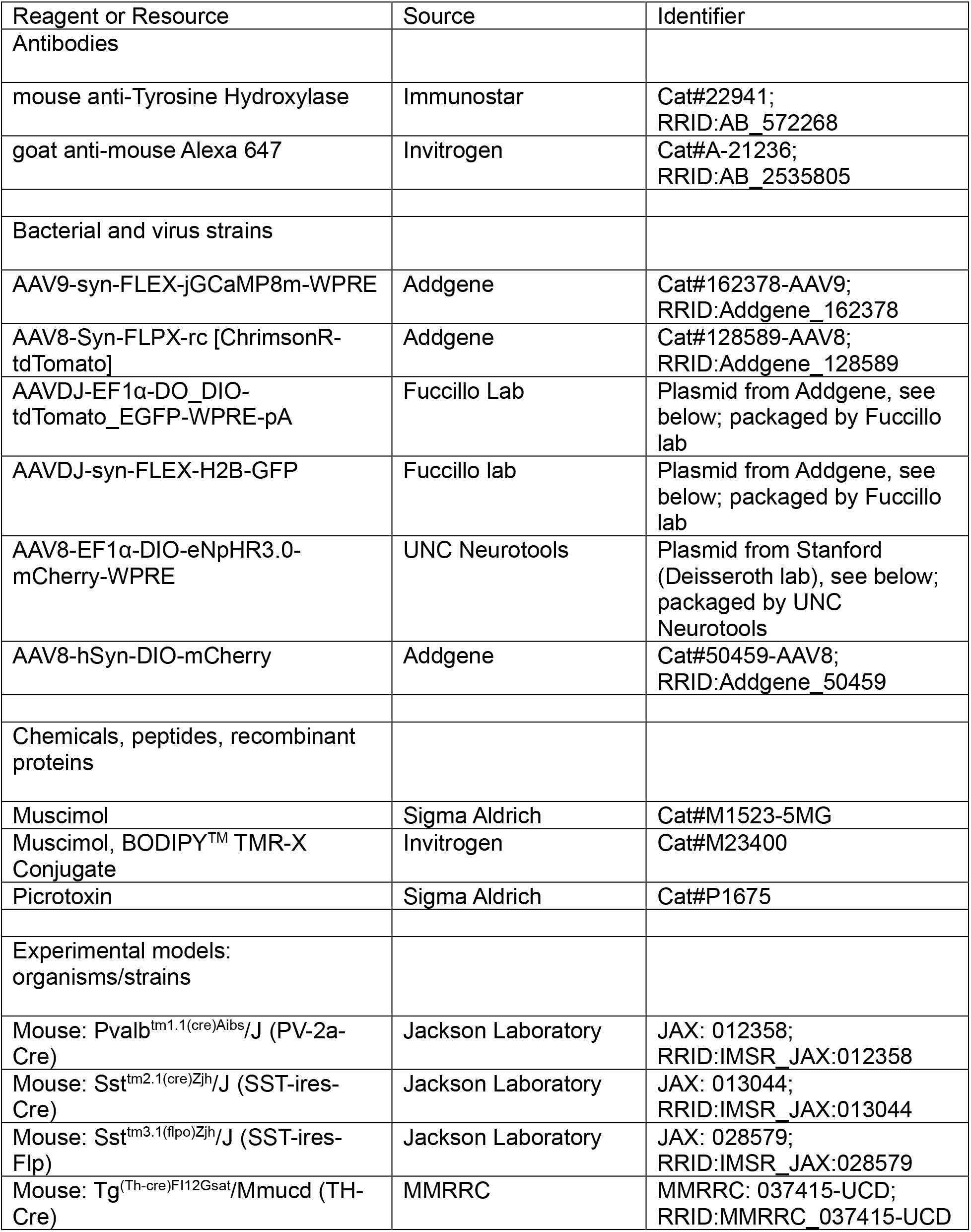
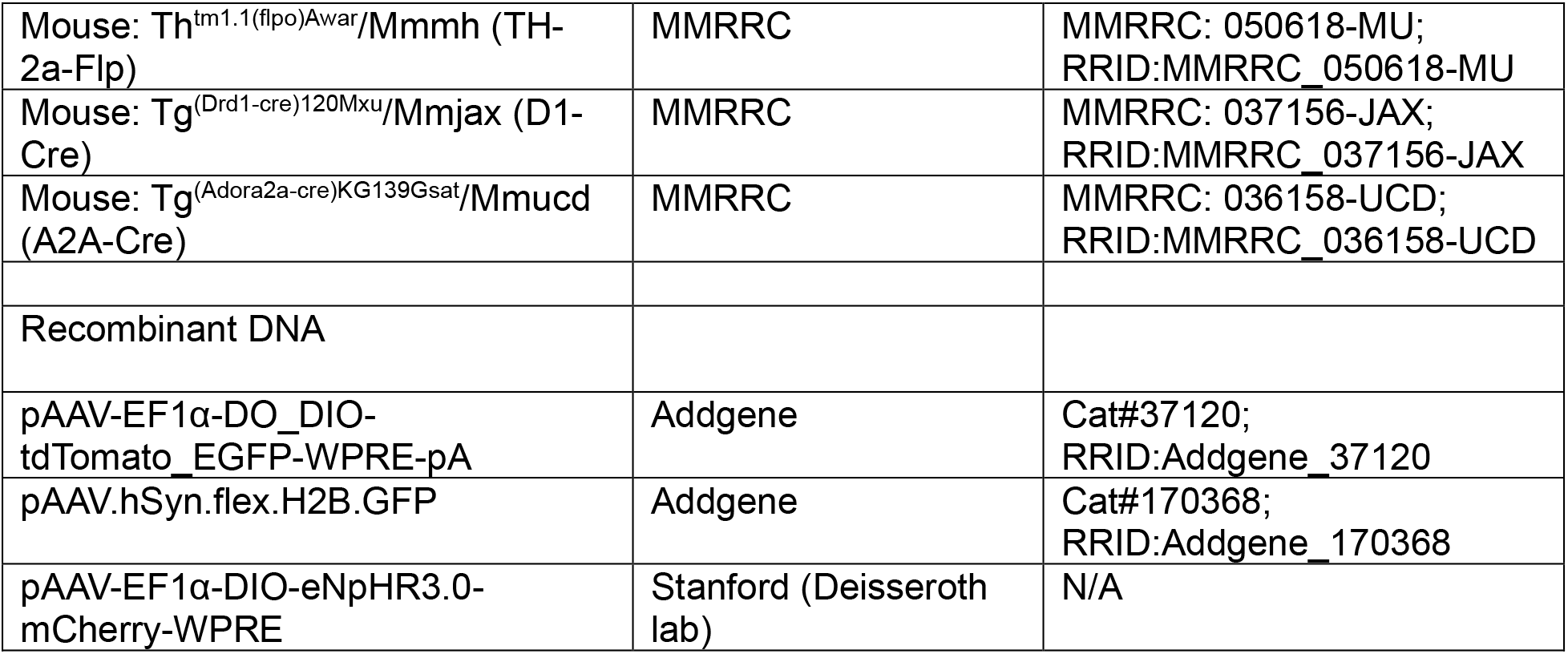

### Animals

All procedures and experiments were conducted in accordance with the Guide for the Care and Use of Laboratory Animals and approved by the University of Pennsylvania Institutional Animal Care and Use Committee (protocol: 805643; confirmation number: aagehdb). All animals used in this study were mice (mean age at training start = 4.4 months, range 1.7 to 8.2 months). All sexes were used. We used wild-type mice (C57BL/6J; the Jackson Laboratory, strain: 000664), and heterozygous (or hemizygous for BAC transgenic TH-Cre, Drd1a-Cre, and Adora2a-Cre lines) mice bred from (1) PV-2a-Cre (Pvalb(tm1.1(cre)Aibs)/J; Jax 012358), (2) SST-ires-Cre (Sst(tm2.1(cre)Zjh)/J; Jax 013044), (3) TH-Cre (Tg(Th-cre)FI12Gsat/Mmucd; MMRRC 037415-UCD), (4) Drd1a-Cre (Tg(Drd1-Cre)120Mxu/Mmjax; MMRRC 037156-JAX), and (5) Adora2a-Cre (Tg(Adora2a-cre)KG139Gsat/Mmucd; MMRRC 036158-UCD).

Homozygous SST-ires-Flp breeders (Sst(tm3.1(flpo)Zjh)/J; Jax 028579) were crossed with either Drd1a-Cre or Adora2a-Cre mice to obtain D1-Cre; SST-Flp and A2a-Cre; SST-Flp double-transgenic mice. Homozygous TH-2a-Flp breeders (Th(tm1.1(flpo)Awar)/Mmmh; MMRRC 050618-MU) were crossed with either SST-ires-Cre, Drd1a-Cre, or Adora2a-Cre mice to obtain SST-Cre; TH-Flp, D1-Cre; TH-Flp, and A2a-Cre; TH-Flp double-transgenic mice.

Control animals for all experiments consisted of age- and sex-matched mixed littermates (details are listed per experiment in the sections that follow). All animals were housed within a 12-hour light/12-hour dark cycle (7am-7pm light; 7pm-7am dark) with food provided ad libitum and a restricted water schedule. Animals were typically group housed (two to five per cage), with the exception of muscimol-treated cohorts (to avoid damage to cranial implants) and individuals displaying aggressive behavior towards cage mates. Sample sizes are detailed in the figure legends.

### Surgical procedures

All surgical procedures were performed on a stereotaxic frame (Kopf Instrument, Model 1900). Briefly, mice were anesthetized using vaporized isoflurane (4% induction; 1 to 2% maintenance; both with oxygen at 1.5 liter/min). Body temperature was continuously monitored using a rectal probe, and mice were maintained on a heating pad set to 30°C during surgery (Harvard apparatus, #50722F; 55-7030). Anesthetic depth was monitored at 5-minute intervals by confirming absence of toe-pinch response. After the administration of ophthalmic ointment (Paralube Vet Ointment) and analgesics (Meloxicam-SR polymer, 4 mg/kg, subcutaneous) and topical anesthesia (bupivacaine, 6 mg/kg, subcutaneous at the incision site), the scalp was prepared for surgery by removing overlying fur with depilatory cream, followed by washes with betadine. An anterior-posterior incision was made using surgical scissors to expose the skull. Residual overlying tissue (periosteum and connective tissue) was removed with brief application of 3% hydrogen peroxide solution, which was washed away with sterile saline. The skull was subsequently crosshatched with scalpel marks for texturization, and acid etched using 30-second application of Metabond Enamel Etchant Gel (Parkell SKU: S395), followed by thorough saline rinse.

For injections and fiber implantations, bilateral craniotomies were made by drilling small (0.5 mm) holes above the target coordinates for the dorsomedial striatum (DMS; anterior-posterior (AP): +0.75 mm from bregma; medial-lateral (ML): ±1.4mm from bregma; dorsal-ventral (DV): −2.85mm from skull surface overlying bregma for injections, −2.55mm for fiber optic cannula placement), dorsolateral striatum (DLS; AP: +0.5, ML: −2.8, DV: −3.8 virus, −3.5 fiber, all unilateral), and posterior dorsomedial striatum (pDMS; AP: −0.2, ML: - 2.0, DV −2.65 fiber and virus, all unilateral). DV coordinates were referenced to the skull surface at bregma.

Injections were performed using the Nanoject II/III system and a pulled glass needle backfilled with mineral oil. Nanoject II was mainly used for earlier photometry experiments, and Nanoject III was preferred for subsequent experiments due to finer control of flow rate and ability to minimize cortical backflow. The glass pipette was left in place for 15 minutes after completion of the injection. Removal rate was not controlled with the Nanoject II, and at a slower rate of 10 μm/s for surgeries with the Nanoject III to reduce cortical backflow.

For photometry experiments in the DMS, the Nanoject II was used to inject 506 nL of undiluted AAV9-syn-FLEX-jGCaMP8m-WPRE (titer: 1.9-2.4 × 10^13^ vg/mL) across 22 sequential 23 nL pulses (flow rate 23 nL/s). For photometry experiments in the DLS and pDMS, the Nanoject III was used to inject 300 nL of diluted AAV9-syn-FLEX-jGCaMP8m-WPRE (titer: 5.0 × 10^12^ vg/mL) at a flow rate of 1 nL/s. All viral dilutions were in sterile saline (0.9%).

For simultaneous fiber photometry and optogenetics, the Nanoject III was used to inject 300 nL of a mixture containing AAV8-AAV-Syn-FLPX-rc [ChrimsonR-tdTomato] (final titer: 5.0 × 10^12^ vg/mL) and AAV9-syn-FLEX-jGCaMP8m-WPRE (final titer: 5.0 × 10^12^ vg/mL) at a flow rate of 1 nL/s. Control animals only received AAV9-syn-FLEX-jGCaMP8m-WPRE (final titer: 5.0 × 10^12^ vg/mL), without ChrimsonR.

For slice electrophysiology experiments in Figure 4, the Nanoject III was used to inject 345 nL of a mixture containing AAV8-AAV-Syn-FLPX-rc [ChrimsonR-tdTomato] (final titer: 4.8 × 10^12^ vg/mL) and either AAVDJ-EF1α-DO_DIO-tdTomato_EGFP-WPRE-pA (titer not independently quantified) for A2A-Cre animals or AAVDJ-syn-FLEX-H2B-GFP (titer not independently quantified) for SST-Cre animals. For interneuron-SPN connectivity experiments, this resulted in expression of ChrimsonR in either SST or TH interneurons, GFP expression in A2A+ D2 SPNs, and tdTomato expression in putative D1 SPNs. For comparisons of synaptic strength of TH interneurons on SST interneurons vs. SPNs, ChrimsonR was expressed in TH interneurons, H2B-GFP in SST interneurons, while putative SPNs remained unlabeled. Injection parameters were 10 nl/s flow rate, injection volume 345 nl.

For optogenetics experiments, the Nanoject III was used to inject 300 nL of AAV8-EF1α-DIO-eNpHR3.0-mCherry-WPRE (final titer: 6.1 × 10^12^ vg/mL) or AAV8-hSyn-DIO-mCherry (final titer: 6.0 × 10^12^ vg/mL) at a flow rate of 1 nL/s.

For all fiber photometry experiments without optogenetics, we used fiber optic cannulae (RWD; R-FOC-BL400C-50NA) with black 1.25-mm ceramic ferrules with 0.5 numerical aperture (NA), 400-μm fiber optic cores at a length of 4 mm. For all optogenetics experiments and fiber photometry experiments with optogenetics, we used fiber optic cannulae (RWD; R-FOC-BL200C-50NA) with black 1.25-mm ceramic ferrules with 0.5 numerical aperture (NA), 200-μm fiber optic cores at a length of 4 mm. Fibers were slowly lowered into the open craniotomies and secured in place with dental cement (Parkell C&B Metabond Quick Adhesive Cement System).

For all animals trained in the task, regardless of whether they received intracranial injections, custom-made stainless steel headplates (Perelman Research Instrumentation Shop; eMachineShop) were secured to the skull using Metabond. The remainder of the headcap was constructed with black-tinted Ortho-Jet dental acrylic (Lang Dental 1320CLR, 3302BLK). For animals destined to receive muscimol injections following training, the area overlying bregma and the injection targets was not covered with cement, but rather sealed with KwikSil Low Toxicity Silicone Adhesive (World Precision

Instruments) for ease of access. Animals were allowed to recover for at least one week before starting water restriction.

### Behavioral apparatus and training

To study contributions of striatal interneurons to value-based choice, mice were trained in a joystick-based probabilistic reversal task as outlined in Linares-García, Iliakis et al.^47^. Briefly, behavioral rigs were custom-built within double-walled sound attenuating chambers containing head-fixation clamps (ThorLabs), a 3D-printed tube to hold the mouse, an optical lickometer (Sanworks 1020), trial start light-emitting diode (‘Go cue’ LED), a house light, a red LED that served as an optogenetics mask, a sucrose solution dispensing system (built with solenoid valve: Lee Company LHDA1231115H), and a sound system. A Hall-effect joystick (5VDC two-axis analog APEM thumbstick; TS6T2S02A, Digi-Key: 679-3658-ND + 3D-printed 8 cm stick) was mounted to a reel holder, which interfaced with a servo motor (SG-5010, Adafruit: 155) that allowed for retraction of the joystick away from the animal between trials. The joystick came in from the right side of the animal, and mice manipulated the joystick with their right forelimb through a gap in the head fixation tube. A horizontal bar beneath the joystick prevented downward displacement of the joystick. Behavioral programs were operated using custom-built circuits with Arduino Mega 2560 and Arduino Uno microcontrollers and custom software. Behavioral data were acquired via serial communication and saved as TXT files using the CoolTerm terminal application.

The reward-delivery system was calibrated using an 8-μL reference volume corresponding to an 80-ms solenoid opening. Valve-opening durations producing the other reward volumes used during training (2.5, 10, and 12.5 μL) were empirically determined relative to this reference. During subsequent calibration checks, these predetermined opening durations were used provided that an 80-ms opening continued to dispense 8 μL.

Analog joystick signals were digitized using the Arduino Mega’s 10-bit analog-to-digital converter. The anteroposterior joystick signal was converted from digital counts to displacement in millimeters using box-specific linear calibration functions hardcoded into the behavioral software. During periods in which the joystick was accessible, raw joystick readings and associated task variables were transmitted over serial communication at a nominal interval of 4 ms (250 Hz). Because serial transmission and program execution introduced variable overhead, the effective acquisition rate was approximately 150 Hz and was not strictly uniform.

For online choice detection, the software maintained a circular buffer containing the 20 most recent internally acquired anteroposterior position samples and calculated their rolling mean. Because this buffer was updated on successive iterations of the behavioral control loop, it constituted a 20-sample rather than fixed-duration moving average. A push or pull choice was registered when the rolling-average position first exceeded the current baseline by +3 mm or −3 mm, respectively for the expert phases, and +1.75 mm or −1.75 mm respectively for early training phases (outlined in detail below). An initial baseline was calculated from 50 samples collected at 50-ms intervals. Between trials, the baseline was recalculated from 15 samples collected at 15-ms intervals 1.25 s after completion of joystick retraction.

Before training, mice were restricted to ≥85% of their pre-restriction baseline weight over the course of 2-3 days. Weights and health were monitored daily during water restriction. If mice did not meet their daily water allotment in the behavior box, they received supplemental water through a syringe immediately after training. Mice were trained for 5-7 days a week, and were given supplemental water to meet their daily water allotment on days off. On Day 1 following the start of water restriction, animals were given supplemental water (0.5-1 mL) in a cage. On Day 2, they were habituated to handling and obtained their water while on the experimenter’s hand. On Day 3, they were habituated to the head fixation tube outside the behavior box, where they received their water through a syringe while inside the head fixation tube. On Day 4 following the start of water restriction, mice were head-fixed in the behavioral apparatus for the first time, and were given non-contingent 2.5 μL 10% w/v sucrose solution rewards every 3-5 seconds for 15-30 minutes as tolerated, with sessions shortened if mice displayed signs of distress. This protocol was repeated for 30 minutes on Day 5. Operant training began on Day 6. Behavioral sessions were limited to 30 minutes.

Operant training proceeded across multiple phases. In Phase 1 (move joystick), mice needed to move the joystick >1.75 mm forward (Push) or backward (Pull) to obtain 10 μL sucrose solution. Throughout, reward was delivered immediately after threshold crossing, which terminated the choice period. The joystick was then immediately retracted for an inter-trial interval of 2.5-8 s (exponential distribution, sampled as 2.5 + Exp(λ = 0.691 s^−1^), capped at 5.5 s to yield a total range of 2.5-8 s). At the start of the next trial, the joystick was extended and Go cue light turned on immediately. There was no penalty for premature choices. Mice were advanced to the next phase when they obtained 50 or more rewards in a single session.

Following this phase, mice were advanced to a debiasing phase (Debiasing I) based on the protocol of Parker et al.^48^. This phase maintained the 1.75 mm decision threshold. However, it required the animal to consistently sample both Push and Pull actions. After five successive choices of the same action, the mice were “locked out” of that action: reward could then only be obtained by registering the opposite action, after which both actions were once again rewarded. After they obtained 50 or more rewards in a single session, mice were advanced to a more difficult debiasing phase (Debiasing II) with a displacement threshold of 3 mm and a lower reward volume of 8 μL. This phase also imposed a 100 ms wait period at trial start after which a Go cue light signaled to the animal it could register a choice. Mice received 15 s time-out penalty for premature choices, during which white noise was played for 5 s, and the house light was turned off. Mice with persistent biases (side bias greater than 75%) were moved to either a Force Push or Force Pull phase in which only one action was rewarded at a displacement threshold of 1.75 mm and a reward volume of 12.5 μL: this encouraged animals to repeatedly sample their dispreferred side and thus counteracted biases, after which they were moved back to the Debiasing II phase. Mice were advanced to the next phase when they obtained over 50 rewards for two consecutive days.

Mice were then advanced to a deterministic reversal phase in which only one of the two directions was rewarded in a given block of trials. Each block had a minimum duration of 17 rewarded trials plus a geometrically-distributed random variable (*p* = 0.4), after which the rewarded side was reversed in an uncued manner. Following a 100 ms wait period, the Go cue light on the lickometer signaled the start of a 10-second choice window. Completed trials produced one of two outcomes: (1) reward delivery, followed by a 2.5-8 s ITI, or reward omission accompanied by extinction of the house light throughout the subsequent 2.5-8 s ITI. Failure to choose within 10 s produced an omission, whereas threshold crossing before the Go cue produced a premature-choice outcome. Both resulted in a 15-s timeout, during which the house light was extinguished and white noise was played for 5 s. Mice were advanced to the final phase when they obtained over 100 rewards with greater than 77% accuracy over the course of two consecutive days.

The probabilistic reversal phase was the main phase used for data collection. It maintained the trial and block architecture of the deterministic reversal phase, but with a ratio of reward probabilities of 80% for the high reward probability vs. 20% for the low reward probability side. The high probability side was flipped in a block-wise fashion, as in the deterministic phase. Expert performance was defined as selection of the higher-probability action on ≥65% of completed choice trials within a session.

Across the 213 mice that began operant training, all mice first obtained ≥50 rewards in a Phase 1 session after a median of 3 training days (mean = 3.76; SD = 2.87; range, 1–17 days). A total of 196 mice first obtained ≥50 rewards in a Debiasing II session after a median of 16 training days (mean = 18.16; SD = 10.55; range, 2–57 days). A total of 164 mice met the deterministic-reversal advancement criterion—defined as >100 rewards with >77% accuracy in each of two consecutive sessions—after a median of 41 training days (mean = 45.01; SD = 18.03; range, 15–111 days). Finally, 140 mice first achieved expert performance in the probabilistic reversal phase after a median of 46 training days (mean = 49.69; SD = 19.10; range, 16–114 days). All durations were calculated from the first day of operant training. Animals that did not reach subsequent milestones were discontinued for reasons including failure to acquire the task, headplate loss, or inadequate photometry signal.

### Histology and anatomical verification

Following completion of experiments, mice were deeply anesthetized with intraperitoneal injection of pentobarbital sodium (150 μL; Sagent, NDC #25021-676-20) and transcardially perfused with 15 mL of 10% phosphate-buffered formalin (Fisher Scientific, SF100-4) containing a total of 50 U heparin (McKesson Medical-Surgical, 1255341), without a preceding saline or PBS flush. Brains were removed and postfixed in 10% phosphate-buffered formalin at 4°C overnight for up to 24 h. Striatal tissue used for anatomical verification was sectioned coronally at 100 μm in phosphate-buffered saline (PBS) using a vibratome (Vibratome Model 1000 Plus or Campden Instruments Model 5100mz). Midbrain tissue used for tyrosine hydroxylase immunohistochemistry was sectioned at 50 μm as described below.

Sections containing the targeted striatal region were mounted and coverslipped using Fluoromount-G (SouthernBiotech, 0100-01) supplemented with DAPI (0.6 μM; Fisher Scientific, D1306). Images were acquired using a Leica DM6 epifluorescence microscope with a 10× objective and assembled as tiled images. Native reporter fluorescence was used to assess viral expression, fluorescent infusate spread, and fiber-tip location, as applicable to each experiment. Animals were excluded if detectable viral expression was absent from the intended target or if the implanted fiber tip was located outside the targeted striatal region. For experiments requiring bilateral targeting, both hemispheres were required to satisfy the applicable anatomical inclusion criteria. Target locations for included animals were registered to the Allen Common Coordinate Framework v3 as described under Mapping anatomic targeting to Allen CCF v3^94^ using SHARCQ.^95^

### Muscimol inactivation

To test the causal contribution of the dorsomedial striatum (DMS) to performance of the joystick-based decision-making task, we transiently inactivated the DMS through bilateral intracranial microinfusion of the GABA_A receptor agonist muscimol. After achieving >77% accuracy in the deterministic reversal phase on two consecutive days, mice advanced to the probabilistic reversal phase and began the infusion protocol. On each behavioral infusion day, mice were anesthetized as described above, secured by their headplates in a modified stereotaxic frame (Kopf Instruments, Model 1900), and administered standard-release meloxicam (5 mg/kg, intraperitoneal), distinct from the sustained-release formulation used during the original headplate surgery.

As described under Surgical procedures, mice assigned to this experiment received headplates that left bregma and the skull overlying the DMS accessible beneath a removable layer of silicone elastomer (Kwik-Sil Low Toxicity Silicone Adhesive, World Precision Instruments). This preparation allowed craniotomies to be deferred until mice had acquired the deterministic reversal task. On the initial infusion day, the silicone elastomer was removed and bilateral craniotomies were made above the DMS (AP: +0.75 mm; ML: ±1.4 mm relative to bregma) using a micromotor drill (Stoelting, 51449).

Vehicle infusions consisted of 40 nL of sterile saline (0.9% NaCl) per hemisphere, delivered into the DMS (DV: −2.85 mm) at 1 nL/s using a Nanoject III. Muscimol infusions consisted of 40 nL of 1.5 mM muscimol in sterile saline per hemisphere, corresponding to 6.85 ng per hemisphere, and were delivered using the same parameters. Following each infusion, the glass capillary was left in place for 7 min and then withdrawn at 15 µm/s. The complete bilateral infusion procedure, including the post-infusion dwell periods and capillary withdrawal, lasted approximately 30 min. The craniotomies were resealed with Kwik-Sil after each procedure. Following completion of the second hemispheric infusion, anesthesia was discontinued and mice were allowed to recover for 30 min before behavioral testing.

The infusion protocol consisted of an initial saline-control session, a muscimol session, and a post-muscimol saline-control session. During the initial saline-control session, mice performed the probabilistic reversal task for the first time after receiving bilateral vehicle infusions. Mice that selected the higher-probability action on ≥65% of completed choice trials proceeded to the muscimol session on the following day and the post-muscimol saline-control session one day later. Mice that did not meet this criterion returned to the deterministic reversal phase until they again achieved >77% accuracy on two consecutive days, after which they repeated the initial saline-control session. No performance criterion was imposed on the muscimol or post-muscimol saline-control sessions.

After completing the final behavioral session, mice were anesthetized as described above and received bilateral infusions of fluorescent muscimol (40 nL per hemisphere; 0.5 mM Muscimol, BODIPY™ TMR-X Conjugate; Thermo Fisher Scientific, M23400) using the same stereotaxic coordinates and infusion parameters. One hour following the completion of surgery, animals were subsequently transcardially perfused as described under Histology and Anatomical Verification. Brains were coronally sectioned at 100 µm using a vibratome and imaged with an epifluorescence microscope to assess the location and spread of the infusate. Anatomical targeting was considered successful when fluorescent muscimol labeling was centered bilaterally within the DMS. Only animals satisfying this criterion were included in the primary analysis. Twelve mice began training; two were excluded because of incomplete behavioral datasets, and four of the remaining ten were excluded because of unsuccessful bilateral DMS targeting, yielding a final analyzed sample of six mice.

### Quantification of joystick kinematic parameters

Joystick kinematics were quantified in the anteroposterior dimension using an approach adapted from Linares-García, Iliakis et al.^47^. Analyses were restricted to completed rewarded and unrewarded trials. For each trial, the rolling-average anteroposterior position signal described under Behavioral apparatus and training was extracted from trial onset, defined by completion of joystick extension, through outcome completion. Trajectories were expressed relative to the trial-specific joystick baseline and temporally aligned to choice registration and outcome onset. Stored position and distance values were divided by 1,000 to convert them to millimeters for downstream analysis.

Signed velocity was calculated as the change in joystick position between successive samples divided by the corresponding elapsed time, thereby accounting for the nonuniform sampling interval. Positive and negative velocities represented push- and pull-directed movement, respectively. The resulting velocity trace was smoothed using MATLAB’s *smoothdata* function with default settings.

Movement bouts were identified from the magnitude and direction of the smoothed velocity signal. A candidate bout was initiated when absolute velocity reached ≥7.5 mm/s. The candidate movement was accepted as a bout after accumulating ≥50 ms at an absolute velocity of ≥2.5 mm/s. An ongoing bout ended upon reaching the end of the extracted trajectory, after accumulating ≥50 ms at an absolute velocity of ≤2.5 mm/s, or following a reversal in velocity sign accompanied by an absolute change in velocity >1.8 mm/s. To ensure detection of the movement producing the registered choice, movement occurring within ±50 ms of choice registration was incorporated into a bout when its absolute velocity was ≥2.5 mm/s, even if the standard initiation criterion had not yet been satisfied.

For each detected bout, direction was assigned from the sign of its mean velocity. Bouts with positive mean velocity were classified as push-directed, whereas bouts with negative mean velocity were classified as pull-directed. A bout was classified as the “decisive” bout when its beginning and end occurred on opposite sides of the choice-registration time; all other bouts were classified as “deliberative.” Bout-based trial-level measures were calculated for trials containing exactly one decisive bout.

Peak displacement was defined as the largest direction-concordant signed displacement during the decisive bout: the maximum displacement for push-directed bouts and the minimum displacement for pull-directed bouts. Mean velocity was the mean signed velocity during the decisive bout. Peak velocity was its largest direction-concordant signed value: the maximum velocity for push-directed bouts and the minimum velocity for pull-directed bouts. Bout number was the total number of detected bouts within the extracted trial trajectory. Directional consistency was calculated as the proportion of bouts whose direction matched that of the decisive bout:

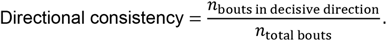

Thus, a directional-consistency value of 1 indicated that every detected bout occurred in the eventual choice direction, whereas lower values indicated the presence of movement bouts in the opposite direction. Path length was defined as the summed absolute sample-to-sample change in joystick position across all detected bouts within the extracted trial trajectory.

### Mapping anatomic targeting to Allen CCF v3 using SHARCQ

Following behavior training, animals were transcardially perfused as described under Histology and Anatomical Verification. Brains were coronally sectioned at 100 µm using a vibratome and imaged with an epifluorescence microscope to assess infusate spread, viral expression, and fiber-tip location, as applicable. For included animals, injection sites in muscimol experiments, and fiber-tip locations in experiments involving fiber-optic cannulae were registered to the Allen Common Coordinate Framework v3^94^ using SHARCQ^95^ (documentation available on Root Lab GitHub under: https://github.com/wildrootlab/SHARCQ). Target locations were manually marked in FIJI and exported as image-space coordinates. Each histological section was matched to the corresponding Allen atlas plane using white-matter tracts, ventricular morphology, and other anatomical landmarks, with atlas tilt adjusted when necessary. At least 10 corresponding landmarks were placed on the histological and atlas images, and SHARCQ used these points to geometrically register the section and transform the marked target location into Allen CCF space. Anteroposterior, dorsoventral, and mediolateral coordinates were then extracted from the registered point.

### Logistic regression model

To assess the impact of trial history across multiple trials on current trial choice, we used a logistic regression model adapted from Lau & Glimcher^49^, Parker et al.^50^, and Alabi et al.^51^. It is defined in Equation 1 below.

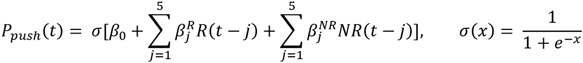

Here, *P*_push_(*t*) is the probability of a push on trial *t*. For each lag *j* = 1, …, 5, *R*(*t* – *j*) was coded as +1 for a rewarded push, −1 for a rewarded pull, and 0 otherwise; *NR*(*t* – *j*) was coded as +1 for an unrewarded push, −1 for an unrewarded pull, and 0 otherwise. The corresponding coefficients, 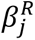 and 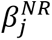, therefore quantify the influence of rewarded and unrewarded choices occurring *j* trials previously on the log-odds of the current choice. Positive coefficients indicate a tendency to repeat the corresponding action, whereas negative coefficients indicate a tendency to select the alternative action. The binary response variable was coded as 1 for push and 0 for pull. Analyses included trials with a valid current choice and five preceding valid-choice trials from the same session. Models were fit separately for each animal in MATLAB using logistic regression with ridge regularization (λ = 0.01) implemented using *fitclinear* without predictor standardization.

### Q-learning model

We adapted a simple Q-learning reinforcement learning model with two parameters to fit the behavioral data produced in our probabilistic reversal task. Mouse choice and outcome history were the primary inputs of the model. The model consisted of an interplay of two steps.

The first step (the ‘Critic’) updated values of both push and pull actions on each trial based on the choice and trial outcome. Values of the choice were estimated in terms of μL of reward and could range from 0 to 8 μL. Values were initiated at 4 μL at the start of each session and updated as follows. For the chosen action:

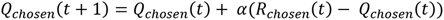

where *Q*_chosen_(*t*+1) refers to the value of the chosen action on the subsequent trial, *Q*_chosen_(*t*) refers to the value of the chosen action on the current trial, *R*_chosen_(*t*) refers to the reward obtained (8 μL vs. 0 μL) from the chosen action.

The difference between *R*_chosen_(*t*) and *Q*_chosen_(*t*) is also referred to as the reward prediction error (RPE). The learning rate α, which ranges from 0 to 1, scales the impact of the reward prediction error on the value updating process. A maximum learning rate of 1 implies complete overwriting of the value information by reward on every trial, whereas a minimum learning rate of 0 implies that reward evidence has no bearing whatsoever on action value estimates.

The value of the non-chosen action decayed on every trial by a factor of 1 – α (one minus the forgetting parameter, set equal to the learning rate in our model):

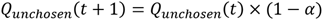

The second step in our Q-learning model (‘Policy’), was modeled with a Softmax rule, which with two variables simplifies to the standard logistic (sigmoid) function, given by:

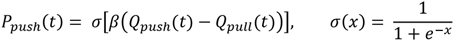

where *P*_push_(*t*) is the push probability on the current trial (and 1 – *P*_push_(*t*) is the pull probability on the current trial), *Q*_push_(*t*) is the value of push on the current trial, and *Q*_pull_(*t*) is the value of pull on the current trial. The difference between these two values gives the value difference ΔQ. The inverse temperature parameter β controls the sensitivity of animals’ choices to ΔQ. High values for β indicate that animals’ choices are almost completely predictable from ΔQ, whereas low values for β imply a greater degree of stochasticity, especially at intermediate values of ΔQ.

This model was selected among several other reinforcement learning models, which incorporated combinations of bias, forgetting, and choice kernel terms (Murphy et al., 2024). The bias term (ε_A_, bias toward action A) quantifies the extent to which animals consistently prefer action A over action B.^49,96^ It is only incorporated in the Softmax decision function and does not impact updating functions:

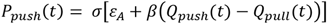

Another possible addition to the Q-learning model is a forgetting term,^97,98^ as we have done in our chosen model. The forgetting term scales the extent to which animals’ inferred value estimates for an action decay when that action is not chosen on a given trial. We evaluated two different versions of a forgetting term: one in which the forgetting term was fit as a separate parameter φ, and another in which the forgetting term was set equal to α, where α is the learning rate. Another option is to omit the forgetting step altogether, resulting in a standard Q-learning model.

Another version we tested included a choice stickiness parameter κ.^99–101^ This term describes the tendency of animals to repeat actions they have executed previously, regardless of their outcome. The choice stickiness parameter was incorporated in the modeling process through inclusion of a choice kernel for each available action. These describe the tendency for an animal to register a given choice on a trial-by-trial basis, and are updated across trials with the following Kernel Updating Equation, one for push (below) and another for pull (identical, omitted for brevity):

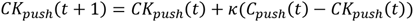

where *CK*_push_(*t*+1) gives the value of the choice kernel on the next trial, *CK*_push_(*t*) gives the value of the choice kernel on the current trial, and C_push_ tracks whether a push was executed on the current trial (1) or not (0). The impact of the current choice on the choice kernel is scaled by the kernel updating rate κ, where lower updating rates imply integrative action tendencies shaped by action history across many trials, whereas higher updating rates imply action tendencies that are shaped exclusively by previous trial choices.

Choice kernels are incorporated in the decision function as a choice kernel difference (Δ*CK* = *CK*_push_ - *CK*_pull_) scaled by a kernel-specific inverse temperature parameter β_K_. The resulting choice function is:

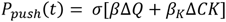

where the contribution of choice kernels to choice was scaled separately by β_K_.

Candidate models were compared using leave-one-session-out cross-validation performed separately for each animal. For each animal and candidate model, parameters were estimated from all but one behavioral session using 25 randomly initialized optimization runs; the parameter set producing the lowest training negative log-likelihood was retained and then held fixed while scoring choices in the omitted session. This procedure was repeated until every session had served once as the test session. Held-out log-likelihoods were summed across sessions within each animal and divided by the total number of held-out trials to obtain an animal-level held-out log-likelihood per trial. Performance of each model was expressed relative to the standard Q-learning model for the same animal and compared across animals using two-sided Wilcoxon signed-rank tests.

### Tyrosine hydroxylase immunohistochemistry

To assess whether striatal delivery of Cre-dependent viral constructs resulted in retrograde transduction of midbrain dopaminergic neurons in TH-Cre mice, we examined native jGCaMP8m fluorescence in tyrosine hydroxylase (TH)-immunolabeled midbrain sections using a protocol adapted from Holly et al.^30^.

Mice were transcardially perfused with 15 mL of 10% phosphate-buffered formalin (Fisher Scientific, SF100-4) containing a total of 50 U heparin (McKesson Medical-Surgical, 1255341), without a preceding saline or PBS flush. Brains were removed and postfixed in 10% phosphate-buffered formalin at 4°C overnight for up to 24 h. For immunohistochemistry, tissue was sectioned coronally at 50 μm in phosphate-buffered saline (PBS) using a vibratome (Vibratome Model 1000 Plus or Campden Instruments Model 5100mz).

Free-floating sections were permeabilized and blocked for 1 h in PBS containing 0.5% Triton X-100, 10% fetal bovine serum, 1% bovine serum albumin, and 0.02% sodium azide. Sections were then incubated overnight at 4°C with mouse anti-TH primary antibody (1:1,000; Immunostar, 22941) diluted in the same solution. After three 5-min washes in PBS, sections were incubated with goat anti-mouse Alexa Fluor 647 secondary antibody (1:200; Invitrogen, A-21236) for 90–120 min, followed by three additional 5-min washes in PBS. Sections were mounted and coverslipped using Fluoromount-G (SouthernBiotech, 0100-01) supplemented with DAPI (0.6 μM; Fisher Scientific, D1306).

Images were acquired using a Leica DM6 epifluorescence microscope with a 10× objective and assembled as tiled images. Native jGCaMP8m fluorescence and TH immunoreactivity were evaluated by visual inspection for the presence of TH-positive, jGCaMP8m-positive neuronal somata within midbrain dopaminergic nuclei.

### Fiber photometry recordings

Batched GCaMP fluorescence was recorded using the Neurophotometrics fiber photometry system (FP3002, MBF Bioscience), while animals performed the behavioral task. Briefly, we used time-dependent modulation in which 415- and 470-nm LEDs were alternated at a combined frequency of 80 Hz (40 Hz per wavelength) to capture isosbestic and calcium-dependent GCaMP fluorescence respectively. Emission was collected via a 400-μm core, 0.57 NA fiber optic patch cord (Doric lenses; FCM – 4 × MF1.25, 400 μm – 0.57 NA; P161376-1) and detected by an internal CMOS camera with submillisecond synchronization. Excitation power at the patch cord tip was calibrated to 80-120 μW per wavelength using an external power meter (Thorlabs: PM100D). Recordings were acquired and initially processed using the FP3002 node on Bonsai, with behavioral events time-locked to the fluorescence signal via 5 V TTL input signals delivered from an Arduino Uno.

### Photometry analysis

Fluorescence signals were preprocessed using a custom MATLAB pipeline adapted from previously published code.^29^ Raw fluorescence measurements were separated into 415-nm isosbestic-control and 470-nm GCaMP channels, and the first 400 samples of each trace were discarded to remove initial signal instability. Each trace was low-pass filtered at 5 Hz using a fourth-order Butterworth filter applied in the forward and reverse directions to avoid phase distortion with *filtfilt* in MATLAB. To correct for photobleaching, a double exponential curve was fit separately to each channel, and fractional fluorescence residuals were calculated by subtracting the fitted curve from the observed fluorescence and dividing by the fitted curve. A robust linear regression was then used to predict the debleached 470-nm trace from the debleached 415-nm trace. The fitted 415-nm-related component was subtracted from the debleached 470-nm trace, and the resulting corrected fractional fluorescence signal was multiplied by 100 to yield isosbestic-corrected %ΔF/F. Finally, this signal was z-scored across the entire retained session by subtracting its session-wide mean and dividing by its session-wide standard deviation to yield z-scored, isosbestic-corrected ΔF/F.

Session-z-scored, isosbestic-corrected ΔF/F signals were aligned to behavioral events using behavior-clock timestamps. For each trial, fluorescence was linearly interpolated onto a common time axis extending from 1 s before to 2 s after the event in 10-ms increments. No additional peri-event baseline subtraction was applied. Trial-aligned traces containing fewer than five finite samples were excluded. For each experimental condition, all eligible trial traces from all included sessions were pooled within animal and averaged pointwise, yielding one peri-event time course per animal. Thus, individual trials were weighted equally within animal, whereas animals were weighted equally in the population average. Population traces show the mean across animal-level time courses, with shaded regions indicating the standard error of the mean (SEM) across animals.

Condition-related differences in fluorescence were evaluated using a paired sign-flip cluster-based permutation test applied to the animal-level peri-event time courses. At each time point, a paired-samples *t*-statistic was calculated across animals. Contiguous time points exceeding an uncorrected cluster-forming threshold of *p* < 0.05 were grouped into positive or negative clusters, with the cluster mass defined as the sum of the absolute signed *t*-statistics within each cluster. Null distributions of maximum cluster mass were generated using 5000 random within-animal sign reversals of the condition differences. Cluster-level *p*-values were calculated using plus-one correction according to the following formula:

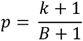

where *B* is the number of permutations (5000) and *k* is the number of permutations in which the maximum null cluster mass in the corresponding direction equaled or exceeded the observed cluster mass. Clusters with permutation-corrected *p* < 0.05 were considered significant. Only animals contributing complete time courses to both conditions were included.

### Neural encoding model

To reduce the computational burden and limit the number of covariates included in the hierarchical Gaussian process model, we first screened candidate predictors using a mass-univariate linear mixed-effects approach. This approach followed the general trial-level, time-resolved regression strategy described by Loewinger et al.^102^, but did not implement their functional linear mixed-modeling framework. Analyses were restricted to trials in the probabilistic reversal phase. Trial-level fluorescence signals were aligned separately to the Go cue (−1 to 1 s) and trial outcome (−1 to 2 s) and resampled into 100-ms bins. At each time bin, a separate linear mixed-effects model was fitted using MATLAB’s *fitlme*, with candidate variables entered as fixed effects and animal included as a random intercept.

The incremental predictive contribution of each variable was evaluated through prespecified nested-model comparisons. Beginning with an intercept-only model, we sequentially added current trial outcome and push–pull choice. Previous trial outcome, signed reward-prediction error (RPE), and absolute RPE were then evaluated as alternative predictors of trial history, and signed and absolute RPE were additionally evaluated after accounting for previous outcome. Movement-related predictors, including peak displacement, mean velocity, and the number of movement bouts, were also evaluated but provided little additional predictive information after push–pull choice was included. Current and previous outcomes were coded as 1 for rewarded trials and 0 for unrewarded trials, whereas choice was coded as +1 for push and −1 for pull. Signed and absolute RPE were scaled by their respective standard deviations without mean-centering. All nested comparisons within a cell type were performed using the same complete-case trials.

Predictive performance was assessed using fivefold cross-validation repeated five times, with animals rather than trials assigned to folds. For each partition, models were trained on four-fifths of the animals, and predictions for the held-out animals were generated using only the fixed-effects estimates, without animal-specific random effects. Fold assignments were held constant across nested-model comparisons and event alignments within each cell type. At each time bin and for each held-out fold, the incremental cross-validated fit of the full relative to the reduced model was defined as:

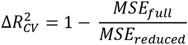

where MSE was calculated from predictions for the held-out trials. Thus, positive values indicated that adding the candidate predictor reduced out-of-sample prediction error. Values were averaged across time bins within predictor-specific, prespecified 1-s windows. Current outcome was evaluated from 0–1 and 1–2 s following outcome delivery; choice was evaluated from 0–1 s following the Go cue and from −1–0 s preceding outcome; previous outcome was evaluated from −1–0 and 0–1 s surrounding the Go cue and from −1–0 s preceding outcome; and RPE-related predictors were evaluated from 0– 1 and 1–2 s following outcome. A predictor was retained if its mean 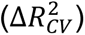 exceeded 0.015 and was positive in at least 17 of the 25 cross-validation evaluations within at least one prespecified window for at least one of the five recorded cell classes (SST, PV, TH, D1, or A2A). This procedure identified current outcome, previous outcome, and push–pull choice for inclusion in the hierarchical Gaussian process model.

Trial-level photometry signals were analyzed separately for each recorded cell type using a custom Bayesian hierarchical Gaussian process model. The complete mathematical derivation, validation, and software implementation of the model will be described separately (Rinaldi et al., in preparation). Briefly, the model expresses the fluorescence signal on each trial as the sum of smooth response functions associated with two temporally distinct task events: the Go cue and trial outcome. Because the interval between these events varies across trials, their respective contributions could be estimated simultaneously without time-warping the signal or analyzing separately averaged event-aligned traces.

For trial *j* from animal *s*, the expected fluorescence signal at time *t* was modeled as:

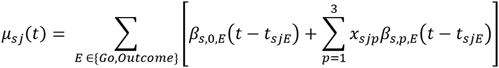

where *t*_sjE_ denotes the occurrence time of event *E*, β_s,0,E_ is an event-aligned intercept function, and β_s,p,E_ describes the time-varying modulation associated with predictor *p*. Based on the predictor-selection procedure described above, the model included variables for current reward, previous reward, and pull choice, with unrewarded trials and push choices serving as the respective reference conditions. Separate coefficient functions were estimated for each predictor and event over windows extending from −1 to 1 s relative to the Go cue and from −1 to 2 s relative to outcome.

Each animal-level coefficient function was modeled as the sum of a shared group-level function and an animal-specific deviation:

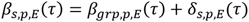

Gaussian process priors were placed on both the group-level functions and animal-specific deviations, allowing coefficients to vary smoothly over time while partially pooling information across animals. Gaussian processes were approximated using a Hilbert-space basis expansion with a Matérn-(5/2) covariance kernel, 40 basis functions, and a boundary-extension factor of 1.5. Group- and animal-level amplitude parameters were assigned exponential priors with rate parameter λ = 0.5 and λ = 3, respectively, and length-scale parameters were assigned gamma priors with shape parameter α = 4 and rate parameter of β = 3. To resolve nonidentifiability between the group functions and the average animal-specific deviation, the basis coefficients defining the animal deviations were assigned zero-sum normal priors, constraining their sum across animals to zero.

Residual fluorescence was modeled using a Student-*t* likelihood to reduce sensitivity to outlying observations. The degrees of freedom parameter ν of the Student-*t* was assigned a gamma prior with shape parameter α = 10 and rate parameter β = 0.5. The model was implemented in PyMC 5.25.1,^103^ and posterior inference was performed using the BlackJAX implementation of the no-U-Turn Sampler.^104^ Four independent chains were run with 1,000 tuning and 1,000 retained draws per chain and a target acceptance probability of 0.95. No divergent transitions were observed.

Under the zero-sum constraint, the fitted group-level HSGP coefficients correspond to the mean coefficient functions across the sampled animals. Their posterior therefore quantifies uncertainty in the latent mean response of those animals, but does not include the additional uncertainty associated with generalizing from the sampled animals to the broader animal population. To approximate the population-level posterior, each posterior draw was augmented in HSGP coefficient space. For each predictor, event, and basis function, a population coefficient was drawn from a normal distribution centered on the fitted group coefficient, with standard deviation equal to the corresponding basis-specific animal-level coefficient scale divided by √N, where N is the number of animals. The resulting coefficients were projected through the HSGP basis to generate draws of the complete population response function, thereby preserving the temporal covariance specified by the animal-level Gaussian process. Plotted functions show the mean and pointwise 95% credible interval of this reconstructed posterior distribution.

### Integrated fiber photometry and optogenetics

For in vivo experiments combining fiber photometry with optogenetic stimulation outside the behavioral task—including TH→SST, TH→D1-SPN, and TH→D2-SPN experiments—mice were allowed at least 2 weeks for viral expression following surgery (see Surgical procedures). Mice were then water-restricted and habituated to handling as described under Behavioral apparatus and training. On the first day of head fixation, mice were connected to a Neurophotometrics FP3002 system (MBF Bioscience) using a four-branch fiber-optic patch cord (NA, 0.37; 1.25-mm ferrules; Neurophotometrics; serial no. SNL2303068091). Fluorescence excitation and emission collection occurred through the same patch cord, which was also used to deliver optogenetic stimulation. Mice received 2.5 μL of 10% (w/v) sucrose solution through the lickometer spout at random intervals of 3–5 s. Independently, red-light stimulation (635 nm; 1-s trains; 20 Hz; 20-ms pulse width; 5 mW measured at the patch-cord tip) was delivered at random intervals of 5–15 s. Thus, sucrose delivery and optogenetic stimulation could overlap by chance. At least 20 stimulation trains were delivered per recording site.

For in-task experiments combining TH-interneuron stimulation with SST-interneuron photometry, mice were subsequently trained to criterion on the probabilistic reversal phase. After achieving ≥65% accuracy, mice advanced to an optogenetic stimulation phase in which red-light stimulation (635 nm; 1-s trains; 20 Hz; 20-ms pulse width; 2.5 mW measured at the patch-cord tip) began at outcome onset on a subset of trials. In this experiment only, stimulation was delivered on randomly selected subsets comprising 33% of rewarded trials and 50% of unrewarded trials. The higher stimulation probability on unrewarded trials ensured sufficient sampling of light-on unrewarded trials, which were comparatively infrequent. In subsequent optogenetic experiments (see Optogenetic manipulations), stimulation was instead delivered on a randomly selected 33% of all trials, independently of trial outcome.

The FP3002 system acquired alternating 415-nm isosbestic and 470-nm calcium-dependent fluorescence frames. Each frame had an exposure duration of 12.5 ms, with each wavelength acquired at 40 Hz, corresponding to a total camera acquisition rate of 80 frames/s. Emitted green and red light was separated by a dichroic mirror, passed through the corresponding bandpass filters, and projected onto spatially distinct regions of the camera sensor. A plastic blocker was inserted into the red emission path before the camera to reduce sensor contamination by the 635-nm optogenetic stimulation light.

Residual stimulation-light artifacts were corrected before filtering, debleaching, and calculation of Δ*F*/*F*. Optogenetic pulse times were reconstructed from the recorded onset of each stimulation train and the stimulation parameters of 20-Hz frequency and 20-ms pulse width. For each 12.5-ms fluorescence-acquisition frame, we calculated the duration of temporal overlap with an optogenetic pulse, denoted *t*_contam_. Overlap was calculated separately for 415- and 470-nm frames using their respective acquisition timestamps.

Successive same-wavelength frames within each stimulation period were grouped into nonoverlapping pairs. For each pair beginning with frame *jj*, we estimated the change in raw fluorescence per unit change in pulse-overlap duration:

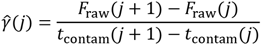

For frame pairs whose pulse-overlap durations differed by more than 4 ms, we used the slope above to estimate and subtract the fluorescence attributable to stimulation-light contamination from both frames:

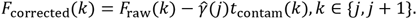

Frame pairs whose pulse-overlap durations differed by 4 ms or less were left unchanged.

Following artifact correction, photometry signals were filtered, processed, aligned, and analyzed as described under Photometry analysis.

### Electrophysiology

Acute slice recordings were performed as described previously.^105,106^ Mice were deeply anesthetized with isoflurane and transcardially perfused with ice-cold artificial cerebrospinal fluid (aCSF; pH 7.3-7.4) containing (in mM): 124 NaCl, 2.5 KCl, 1.2 NaH_2_PO_4_, 24 NaHCO_3_, 5 HEPES, 12.5 glucose, 1.3 MgSO_4_, and 2.5 CaCl_2_. The brain was rapidly removed, and 250-μm coronal sections were cut on a vibratome (VT1200s, Leica) in aCSF. Slices were incubated in a holding chamber for approximately 15 minutes at 32-34°C in an N-methyl D-glucamine (NMDG)-based recovery solution (pH 7.3-7.4, adjusted with HCl or NaOH as needed) containing (in mM): 92 NMDG, 2.5 KCl, 1.2 NaH_2_PO_4_, 30 NaHCO_3_, 20 HEPES, 25 glucose, 5 sodium ascorbate, 2 thiourea, 3 sodium pyruvate, 10 MgSO_4_, and 0.5 CaCl_2_. Osmolarity for the NMDG-based solution and aCSF was kept between 300-310 mOsm. Following incubation, slices were maintained in aCSF at room temperature (20-22°C) for at least 1 hour before recording.

For recording, slices were transferred to the recording chamber (Scientifica) fully submerged in carbogenated aCSF at a perfusion rate of 1.4-1.6 mL/min, bath temperature of 29-30°C, and secured using a slice anchor (Warner Instruments). Drugs were prepared in stock solutions in water or DMSO and diluted to their final concentration in aCSF. The final concentration of DMSO was < 0.1%. Electrophysiological data were acquired using custom-built Recording Artist software (Richard C. Gerkin; GitHub: https://github.com/rgerkin) and Igor Pro 6.37 (WaveMetrics). Recordings were sampled at 20 kHz and low-pass filtered at 2.8 kHz.

For interneuron-connectivity experiments, whole-cell voltage-clamp recordings were performed in aCSF containing no pharmacological blockers. Recording electrodes (3–5 MΩ) were pulled from borosilicate glass and filled with a chloride-loaded intracellular solution containing (in mM): 135 CsCl, 10 HEPES, 0.6 EGTA, 2.5 MgCl_2_, 10 Na-Phosphocreatine, 4 Na-ATP, 0.3 Na-GTP, 0.1 spermine (pH 7.3; 310 mOsm/L). ChrimsonR-expressing interneurons were stimulated through a 40× water-immersion objective (Olympus; 0.8 NA) using 2-ms pulses of 530-nm light (1-3 mW). Optogenetically evoked inhibitory postsynaptic currents (oIPSCs) were recorded at a holding potential of −70 mV and appeared as inward currents owing to the chloride-loaded intracellular solution. For interneuron-to-SPN experiments, red fluorescent dSPNs and green fluorescent iSPNs located within the same field were patched sequentially. For TH-to-SST/SPN experiments, recordings were obtained from green fluorescent SST interneurons and unlabeled putative SPNs sequentially within the same field. Putative SPNs were identified based on input resistance less than 300 ΜΩ. Voltages were not corrected for liquid junction potential. Recordings were excluded if access resistance increased by more than 20%.

For SPN AMPA/NMDA (A/N) ratio experiments, whole-cell voltage-clamp recordings were performed in aCSF containing 100 μM picrotoxin to block GABA_A_-receptor-mediated currents. Recording electrodes (3–5 MΩ) were pulled from borosilicate glass and filled with an intracellular solution containing (in mM): 115 CsMeSO₃, 20 CsCl, 10 HEPES, 0.6 EGTA, 2.5 MgCl₂, 10 Na-phosphocreatine, 4 Na-ATP, 0.4 Na-GTP, 0.1 spermine, and 1 QX-314 (pH 7.3–7.4; 290 mOsm/L). EPSCs were evoked using a bipolar stimulating electrode constructed from two tungsten wires housed in theta glass and positioned in the dorsomedial striatum (DMS). Electrical stimuli (1 ms duration) were delivered at 10s intervals, and stimulation intensity was held constant between recordings at −70 and +40 mV. AMPA/NMDA ratios were calculated by comparing the peak amplitude of AMPAR-mediated EPSCs recorded at −70 mV across 10 sweeps with the NMDAR-dominated current measured 50 ms after stimulus onset at +40 mV across 15 sweeps. Sweeps obtained at each holding potential were averaged before measurement. Peak AMPAR amplitude was quantified as the maximum inward current relative to the pre-stimulus baseline, and absolute current amplitudes were used to calculate the AMPA/NMDA ratio. Series resistance and whole-cell capacitance were monitored throughout the recording, and recordings were excluded if series resistance changed by more than 20% during the experiment.

### Optogenetic manipulations

To gain optogenetic control of SST or TH interneurons, we injected the inhibitory opsin eNpHR3.0 bilaterally into the dorsomedial striatum of SST-Cre or TH-Cre mice, respectively, as described under Surgical procedures. Control animals received the corresponding fluorophore-only construct. Mice were allowed to recover for at least 1 week before beginning behavioral training, which proceeded as described under Behavioral apparatus and training. Opsin-expressing and control animals underwent identical behavioral, tethering, and stimulation protocols. All session criteria and progression rules were specified before data collection and applied identically across experimental groups.

Valid completed trials (*V*) comprised rewarded and unrewarded trials in which a choice was registered during the choice window. Trials were classified as omissions (*O*) when no choice was registered during the choice window and as premature trials (*P*) when a choice was registered before choice-window onset. Choice accuracy was defined as the percentage of valid trials directed toward the higher-reward-probability side. A session met validity criteria when it contained ≥100 valid trials, *O*/*V* ≤ 0.40, and *P*/*V* ≤ 0.40. Behavioral collapse was defined by any of the following: fewer than 60 valid trials, *O*/*V* > 0.60, *P*/*V* > 0.60, or choice accuracy <50%. Sessions falling between the validity and collapse thresholds were considered invalid but noncollapsed.

After training on the deterministic reversal phase, animals were required to complete two consecutive sessions that met the validity criteria and had ≥77% choice accuracy. Animals meeting these criteria advanced to the probabilistic reversal phase without optogenetic stimulation. They remained in this phase until completing two consecutive sessions that met the validity criteria and had ≥65% choice accuracy. These two sessions constituted the pre-stimulation baseline (“Pre”). Beginning with the first Pre session, animals were tethered bilaterally during all subsequent experimental sessions, including sessions without light delivery.

Optogenetic stimulation was delivered using a TTL-triggered 625-nm LED (Optogenetics-LED-Dual-OR/OR, Prizmatix). Light passed through a 1.5-mm-core source fiber (NA, 0.63; OptoGenetics Fiber-1500, Prizmatix), a stationary FC–FC mating connector (Prizmatix), and a bifurcated patch cord with a 200-μm core per branch (NA, 0.57; SBP(2)_200/230/900-0.57_1m_FCM-2xMF1.25, Doric Lenses), through which animals were tethered bilaterally. Continuous light was delivered bilaterally for 1 s beginning at outcome onset on a randomly selected 33% of valid choice trials. Trial selection was independent of choice direction and reward outcome. Light power measured at the patch-cord tips ranged from 4 to 7 mW across sessions.

Each animal was required to complete six criterion optogenetic sessions. An optogenetic session counted toward this total when it met the validity criteria and choice accuracy was ≥65%; sessions not meeting both criteria did not count. After the first three criterion optogenetic sessions, animals completed one light-off probe session under otherwise identical probabilistic contingencies. The probe session was required to meet the validity criteria and have ≥65% choice accuracy; otherwise, light-off probe sessions were repeated until both criteria were satisfied. Animals then completed three additional criterion optogenetic sessions.

If an optogenetic session failed the validity criteria without meeting a collapse criterion, subsequent sessions were conducted under probabilistic contingencies without stimulation until one session met the validity criteria, after which optogenetic sessions resumed. If an animal met any collapse criterion, it was returned to deterministic light-off training until completing one session that met the validity criteria. The animal was then returned to probabilistic light-off training until completing one session that met the validity criteria, after which optogenetic sessions resumed from the point at which they had been interrupted. Recovery and retraining sessions were recorded but excluded from the primary phase-based behavioral analyses.

After completing six criterion optogenetic sessions, animals underwent probabilistic light-off sessions to assess the persistence of behavioral effects (“Post”). Four Post sessions meeting the validity criteria were collected from each animal. Additional light-off sessions were conducted as necessary until four criterion Post sessions were obtained, with the same validity and collapse criteria applied throughout.

For animals designated for ex vivo electrophysiology, two additional optogenetic “booster” sessions were administered after completion of the Post phase. Both booster sessions were required to meet the validity criteria. Animals were euthanized 24–48 h after the second booster session for slice electrophysiology.

### Latent state model for analysis of optogenetic data

#### Trial selection and predictor construction

Behavioral models were fitted jointly across animals to identify choice strategies and latent states shared across the population, rather than fitting separate latent states for individual animals. The final modeling dataset contained 123,714 trials from 942 sessions in 38 animals. The model was constructed from animals in the optogenetics experiment, including all their probabilistic reversal sessions, regardless of performance, optogenetic stimulation, and control vs. inhibition group. Animals associated with more than one behavioral box in the assembled dataset were excluded.

Choices were coded as push (+1) or pull (−1), with the Bernoulli response equal to 1 for push and 0 for pull. Reward was coded as 1 when trialOutcome = 1 and 0 otherwise. Trials were required to contain a push or pull response and a choice latency between 0 and 10 s. The assembled design table was then restricted to complete cases. Because the design matrix contained lagged and recent-history predictors, early trials within each session for which the required history was unavailable were consequently excluded.

Previous choice and outcome variables were calculated only within the same animal and behavioral session. Rewarded-choice and unrewarded-choice history terms were defined as the signed previous choice multiplied by indicators for whether the previous trial was rewarded or unrewarded, respectively.

Recent choice history was defined using the previous five available choices in the same session, excluding the current trial:

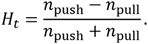

Thus, positive values indicated a recent bias toward pushing and negative values indicated a recent bias toward pulling. Partial windows were permitted near the beginning of a session. For the online RL-GLM and RL-GLM-HMM analyses, this predictor was divided by its within-animal standard deviation without subtracting the animal mean, thereby preserving its sign.

Recent switching was calculated over the previous 12 available choices within the same session as

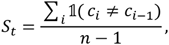

where *n*was the number of choices available in the window. At least two previous choices were required. Switching fraction was standardized within animal by subtracting the animal mean and dividing by its standard deviation, and entered the policy model as its interaction with the signed previous choice. The five-trial choice-history and 12-trial switching windows were selected by comparing candidate history depths using Akaike information criterion.

An animal-level choice-bias predictor was defined as the difference between that animal’s overall push and pull fractions. This value was calculated once using the full selected dataset and was constant across trials from the same animal.

#### Q-learning model

Implementation of our Q-learning model is discussed in detail in the Q-learning model section of the Methods. Briefly, action values were initialized to 4 and reset at the beginning of each session. Binary rewards were multiplied by 8 to express reward magnitude on the same scale as the initialized action values. Following choice of action *a_t_*, the value of the chosen action was updated according to

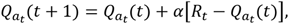

whereas the value of the unchosen action decayed according to

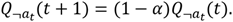

The relative action value was defined as

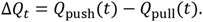

For the initial fixed-Δ*QQ* model comparison, a Q-learning model with a single learning rate and inverse-temperature parameter was fitted to the complete dataset. Parameters were estimated using 30 starting points and constrained optimization in MATLAB. The learning rate was bounded between 0.0001 and 0.9999 and inverse temperature between 0.0001 and 10. Trial-wise values generated using the retained parameters were subsequently held fixed during evaluation of the Bernoulli policy models.

#### Bernoulli policy models

A series of pooled Bernoulli generalized linear models with a logit link was evaluated. Predictors were added sequentially to a model containing ΔQ: animal bias; rewarded- and unrewarded-choice history; five-trial recent choice history; and previous choice interacted with 12-trial switching fraction. The resulting single-value-slope model was

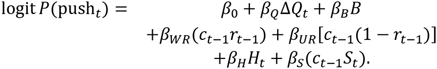

A second model allowed the influence of action value to differ depending on its sign by replacing ΔQ with

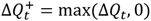

and

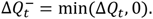

These models corresponded to the final single-ΔQ-slope and split-ΔQ-slope Bernoulli GLMs.

#### Session-level model evaluation

Five predefined cross-validation folds were constructed at the session level, such that all trials from a session were assigned to the same fold. Sessions were defined by animal and session identifiers. Animals were ordered by their total number of valid trials, sessions were reproducibly shuffled within animal, and sessions were assigned greedily to the eligible fold containing the fewest trials. Whenever possible, different sessions from a given animal were distributed across different folds before a fold was reused. Fold assignment was performed once and reused for all model classes.

The resulting folds contained 188-189 sessions and 23,903-25,448 trials each. On each iteration, models were fitted to four folds using data pooled across animals and evaluated on the remaining sessions. Predictive performance was quantified as the unpenalized held-out log likelihood per trial:

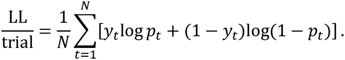

For HMMs, the corresponding session-level marginal likelihood was obtained using the forward algorithm, summed across held-out sessions, and divided by the number of held-out trials. Animal-level scores were obtained by summing each animal’s held-out log likelihood and trial count across folds before calculating log likelihood per trial.

The animal-bias predictor and the parameters used to generate fixed ΔQ were estimated once from the full selected dataset before fold-wise GLM fitting. Accordingly, these analyses should be interpreted as held-out session scoring conditional on globally estimated feature representations rather than as completely nested estimates of performance on an independent population.

#### Online RL-GLM

The selected policy structure was next fitted as an online RL–GLM in Python. In this model, the learning rate was estimated jointly with the policy coefficients and the ΔQ trajectory was recomputed inside each likelihood evaluation. A single learning rate and coefficient vector were shared across animals, while action values were reset at session boundaries.

Parameters were estimated using bounded L-BFGS-B optimization. The learning rate was restricted to

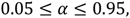

policy coefficients were bounded between −20 and 20, and L_2_ penalties of 10^−3^ were applied to the policy coefficients. Multiple initializations were fitted within each training fold, and the initialization with the lowest penalized training negative log likelihood was retained. Held-out performance was evaluated using the unpenalized Bernoulli log likelihood.

#### RL-GLM-HMM

The online RL–GLM was incorporated into a hidden Markov model adapted from the publicly available Iris Stone *glmhmm* implementation^56^ (https://github.com/irisstone/glmhmm). Two- and three-state models were evaluated. For latent state *k*, choice probability was

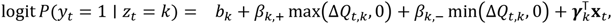

where *x_t_* contained rewarded- and unrewarded-choice history, animal bias, recent choice history, and the interaction between previous choice and recent switching. Each state had its own learning rate, intercept, positive and negative ΔQ coefficients, history coefficients, and corresponding Q-value trajectory. Each state’s values were updated in parallel using the animal’s observed choices and outcomes.

The latent states followed a first-order Markov process with a transition matrix shared across animals and sessions. Forward-backward inference was performed separately for each session, preventing transitions across session boundaries. The initial state distribution was fixed to a uniform distribution.

Parameters were estimated by expectation maximization. Transition probabilities were updated from expected transition counts, and state-specific emission parameters were optimized during the M-step using bounded L-BFGS-B optimization. Learning rates were bounded between 0.05 and 0.95, coefficients between −20 and 20, and L_2_ penalties of 10^−3^ were applied.

For cross validation, multiple initializations were fitted within each training fold, and the initialization with the greatest final training log likelihood was used to score the held-out fold. For the final three-state model fitted to the complete dataset, ten random initializations corresponding to seeds 0-9 were evaluated. The fit initialized with seed 3 produced the greatest log likelihood and was retained for subsequent state analyses.

After fitting, state labels were aligned by sorting states in descending order of their rewarded-choice-history coefficient. This relabeling affected only the state identities and did not alter model likelihood.

#### Posterior state assignment and state characterization

Trial-wise state probabilities were calculated using forward-backward inference. A hard state assignment was defined as the state with the greatest marginal posterior probability on that trial:

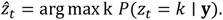

This assignment was a marginal maximum-a-posteriori assignment and was not a Viterbi path.

Hard assignments were used to calculate each animal’s push fraction, action-specific win–stay and lose–switch probabilities, choice accuracy, percentage of trials assigned to each state, mean state-run duration, and empirical state-transition counts. Win–stay and lose–switch measurements required the preceding observation to be a consecutive trial from the same session.

For the plots of push probability as a function of ΔQ, population-level state curves were posterior-weighted. Trials were divided into ΔQ bins centered from −8 to 8 in increments of 2, and state/bin combination was plotted when its effective posterior-weighted trial count was at least 20.

Empirical transition counts were calculated between successive retained observations belonging to the same animal and session. The fitted HMM transition matrix and the empirical row-normalized matrix represented the probability of the next state conditional on the current state. For visualizations restricted to changes between states, diagonal counts were removed before normalization. Row normalization then represented the destination conditional on leaving a given state:

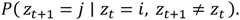

State characteristics were compared using paired Wilcoxon signed-rank tests at the animal level. Holm corrections were applied separately within each planned family of comparisons: state-versus-overall push fractions, action- and outcome-specific stay/switch measures, choice accuracy, state occupancy, and state duration.

#### Optogenetic effects on latent-state probabilities

Optogenetic analyses used posterior state probabilities rather than hard assignments. Sessions were grouped into PRE, early optogenetic manipulation, late optogenetic manipulation, early post-manipulation, and late post-manipulation bins. For each animal and experimental bin, posterior probabilities were averaged separately for each state.

SST and TH cohorts were analyzed separately. For each state, mean posterior probability was modeled using a linear mixed-effects model containing condition, experimental bin, and their interaction as fixed effects and an animal-specific random intercept:

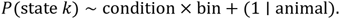

Models were fitted by maximum likelihood.

## Acknowledgments

This work was supported by NIMH F30 MH136699 to EAI, NINDS F31 NS130989 to SMF, NIMH F31 MH125542 to LV, NINDS R01 NS094450 to DJM, NIMH R01 MH118369 and NIMH RF1 MH138591 to MVF. SP is supported as a Penn ASPE postdoctoral fellow. KC was supported as a former Penn ASPE postdoctoral fellow.

This work was supported by the European Union – NextGenerationEU – PNRR M4C2-I.1.1 through PRIN Project no. 2022XE8X9E, CUP G53D23004590001, granted by the Italian government (E.P.).

We would like to thank Alessandro Jean-Louis, Myra Granato, and Wenxin Tu for outstanding technical assistance, and Maxime Assous and Fulva Shah for TH-Cre transgenic mice. We also thank Yuma Kajihara for his contributions to our behavioral models. We would like to thank University Laboratory Animal Services staff in Penn Clinical Research Building and John Morgan animal facilities for outstanding mouse husbandry support, with special thanks to Lenny Coleman. We would like to thank Alessandro Jean-Louis and Nathan Henderson for their support in cloning of viral constructs for experiments that were ultimately not included in this manuscript.

We would like to thank Olivia Hon and Sage Aronson from Neurophotometrics/MBF Bioscience, for their outstanding technical assistance and support with troubleshooting our fiber photometry and simultaneous fiber photometry and optogenetics experiments. We thank Laura Haetzel, Laura McGarry, Sinéad Moyles, and Kayla Peelman for valuable conceptual input and feedback on paper figures.

3D printed / fabricated objects courtesy of the University of Pennsylvania Libraries’ Holman Biotech Commons with design consultation (RRID:SCR_024735), with special thanks to Varvara Kountouzi. Mechanical assistance courtesy of Penn Research Instrumentation Shop (RRID:SCR_022424), with special thanks to Dieter Hunt. Electronic designs courtesy of Penn Electronic Design Shop (RRID:SCR_021107), with special thanks to Vincent Lau.

## Author contributions

EAI – conceptualization, data curation, formal analysis, funding acquisition, investigation, methodology, project administration, resources, software, supervision, validation, visualization, writing – original draft, writing – review and editing.

JMM – conceptualization, data curation, formal analysis, investigation, methodology, resources, validation, visualization, writing – review and editing.

ANR – conceptualization, data curation, formal analysis, investigation, methodology, resources, validation, writing – review and editing.

FR – conceptualization, data curation, formal analysis, investigation, methodology, software, validation, visualization, writing – review and editing.

JG – conceptualization, data curation, formal analysis, investigation, methodology, resources, validation, visualization, writing – review and editing.

SP – data curation, formal analysis, investigation, resources, visualization, writing – review and editing.

ILG – conceptualization, investigation, methodology, resources, software, writing – review and editing.

ART – data curation, investigation, resources, validation, visualization, writing – review and editing.

EZS – investigation, resources, validation, writing – review and editing.

LV – conceptualization, methodology, software, writing – review and editing.

KC – data curation, formal analysis, investigation, methodology, resources, validation, visualization, writing – review and editing.

JTW – conceptualization, data curation, formal analysis, investigation, methodology, software, validation, visualization, writing – review and editing.

SMF – conceptualization, investigation, methodology, resources, software, writing – review and editing.

EDH – conceptualization, investigation, methodology, resources, software, writing – review and editing.

ENH – conceptualization, funding acquisition, validation, writing – review and editing.

DJM – conceptualization, funding acquisition, project administration, supervision, writing – review and editing.

EP – conceptualization, funding acquisition, investigation, methodology, project administration, resources, software, supervision, validation, visualization, writing – review and editing.

MVF – conceptualization, data curation, funding acquisition, methodology, project administration, resources, supervision, validation, visualization, writing – original draft, writing – review and editing.

## Code availability

Analysis code for this project is available on the Fuccillo Lab GitHub (https://github.com/Fuccillo-Lab/Iliakis-Striatal-Interneuron-bioRxiv-2026).

## Declaration of generative AI and AI-assisted technologies

During the preparation of this work, E.A.I. used OpenAI ChatGPT (GPT-4, GPT-4o, and GPT-5-family models) to improve the readability and language of the manuscript, and for assistance with drafting of analysis code. After using this service, the authors reviewed and edited the content as needed and take full responsibility for the content of the published article and analysis code.

