## Supplemental Figures for "Striatal interneuron microcircuits gate reinforcement to stabilize adaptive choice"

**Iliakis et al. (2026)**

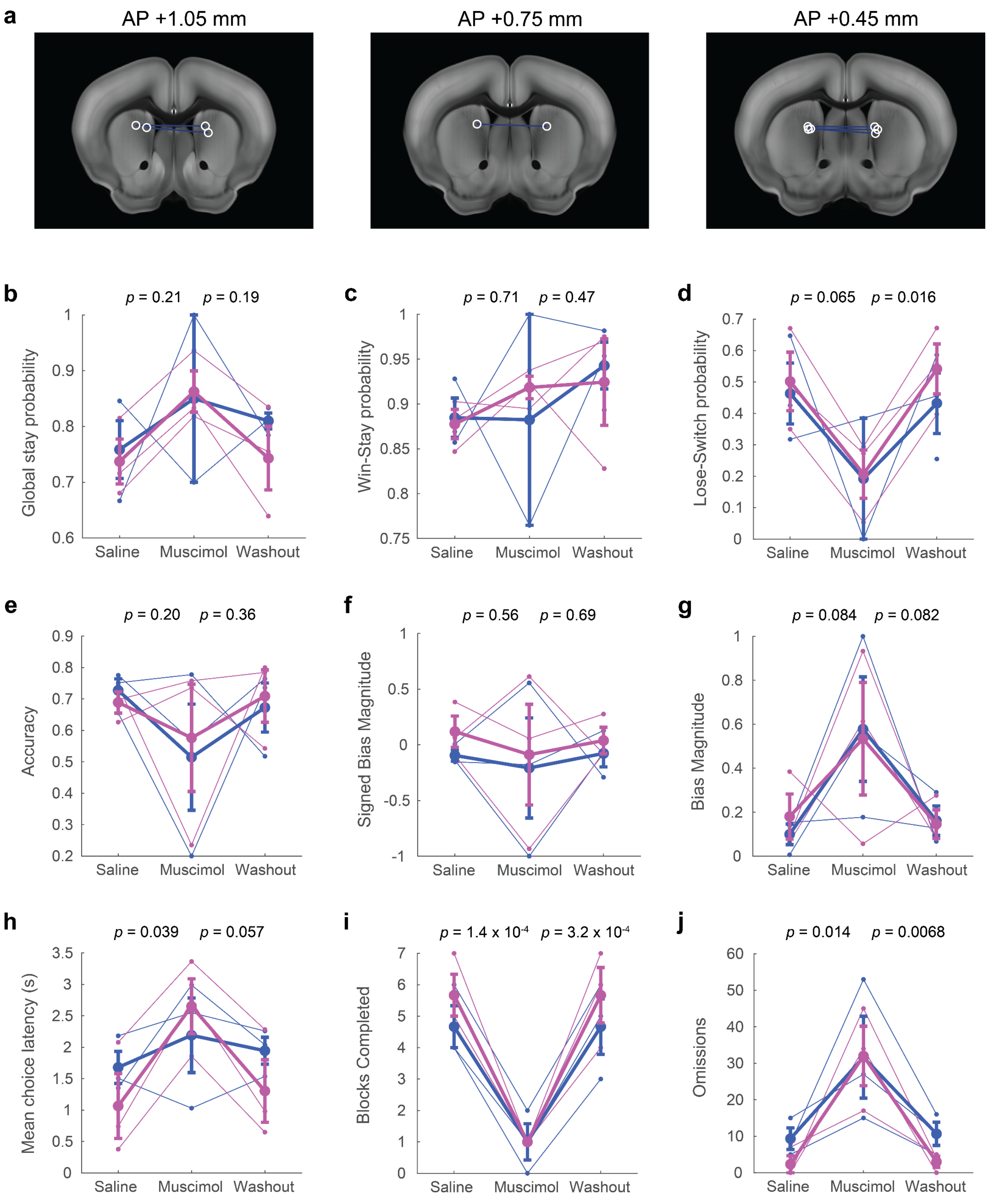


**Figure S1. Inhibition of the dorsomedial striatum with muscimol impairs execution of the probabilistic reversal task.**

(a) Anatomic targeting of included animals (n = 6; 3 females, 3 males) registered to Allen Mouse Brain Common Coordinate Framework version 3[1] using SHARCQ[2] and shown on coronal sections at 300-μm intervals. Approximate infusion sites were determined by post hoc bilateral infusion of fluorescent muscimol (40 nL per hemisphere, 0.5 mM Muscimol, BODIPY™ TMR-X Conjugate) using the stereotaxic coordinates and infusion parameters employed in the behavioral experiments. White circles indicate approximate infusion sites, blue traces connect infusion sites from the same animal.

(b-j) The infusion protocol consisted of an initial saline-control session ("Saline"), a muscimol session ("Muscimol"), and a post-muscimol saline-control session ("Washout"). The following measures were compared across sessions: (b) global stay probability, (c) win-stay probability, (d) lose-switch probability, (e) accuracy, (f) signed bias magnitude, (g) bias magnitude, (h) mean choice latency, (i) blocks completed, and (j) omissions. Thin lines represent individual animals; thick lines and error bars represent mean ± SEM for males (blue) and females (magenta). The displayed *p*-values are from paired, two-tailed *t*-tests comparing Saline with Muscimol, and Muscimol with Washout.

Global stay probability was the probability of repeating the preceding choice, irrespective of its outcome. Win-stay probability was the probability of repeating a rewarded choice, whereas lose-switch probability was the probability of selecting the alternative action after an unrewarded choice. Only pairs of consecutive valid choice trials contributed to these measures; omission trials and trials containing a premature choice before the Go cue were excluded. Accuracy was the proportion of valid choices directed toward the high-reward-probability alternative.

Signed bias magnitude was defined as:

$$bias_{signed} = \frac{N_{push} - N_{pull}}{N_{push} + N_{pull}},$$

such that negative and positive values indicated push and pull biases, respectively. Unsigned bias magnitude was the absolute value of signed bias, with larger values indicating a stronger bias towards either action.

Choice latency was the elapsed time from Go-cue onset to choice-threshold crossing. Blocks completed was the number of blocks in which transition criterion was met; a block transition required at least 17 rewarded choices on the high-reward-probability side. Omissions were trials in which no choice was registered during the 10-s choice window.

Effects of Condition and Sex were assessed using a linear mixed-effects model of the form: Metric ~ Condition * Sex + (1 | animalID), with Condition comprising Saline,

Muscimol, and Washout. No significant main effects of Sex were observed (all $p \geq 0.05$). A significant Condition × Sex interaction was observed for choice latency ($F(2,12) = 5.2$, $p = 0.024$); no significant interactions were observed for the remaining measures (all $p \geq 0.05$). Significant main effects of Condition were observed for blocks completed ($F(2,12) = 61.9$, $p = 4.8 \times 10^{-7}$), omissions ($F(2,12) = 13.1$, $p = 0.00096$), lose-switch probability ($F(2,10.8) = 6.9$, $p = 0.012$), and choice latency ($F(2,12) = 19.2$, $p = 0.00018$). No significant main effects of Condition were observed for global stay probability, win-stay probability, accuracy, signed bias magnitude, or unsigned bias magnitude (all $p > 0.05$).

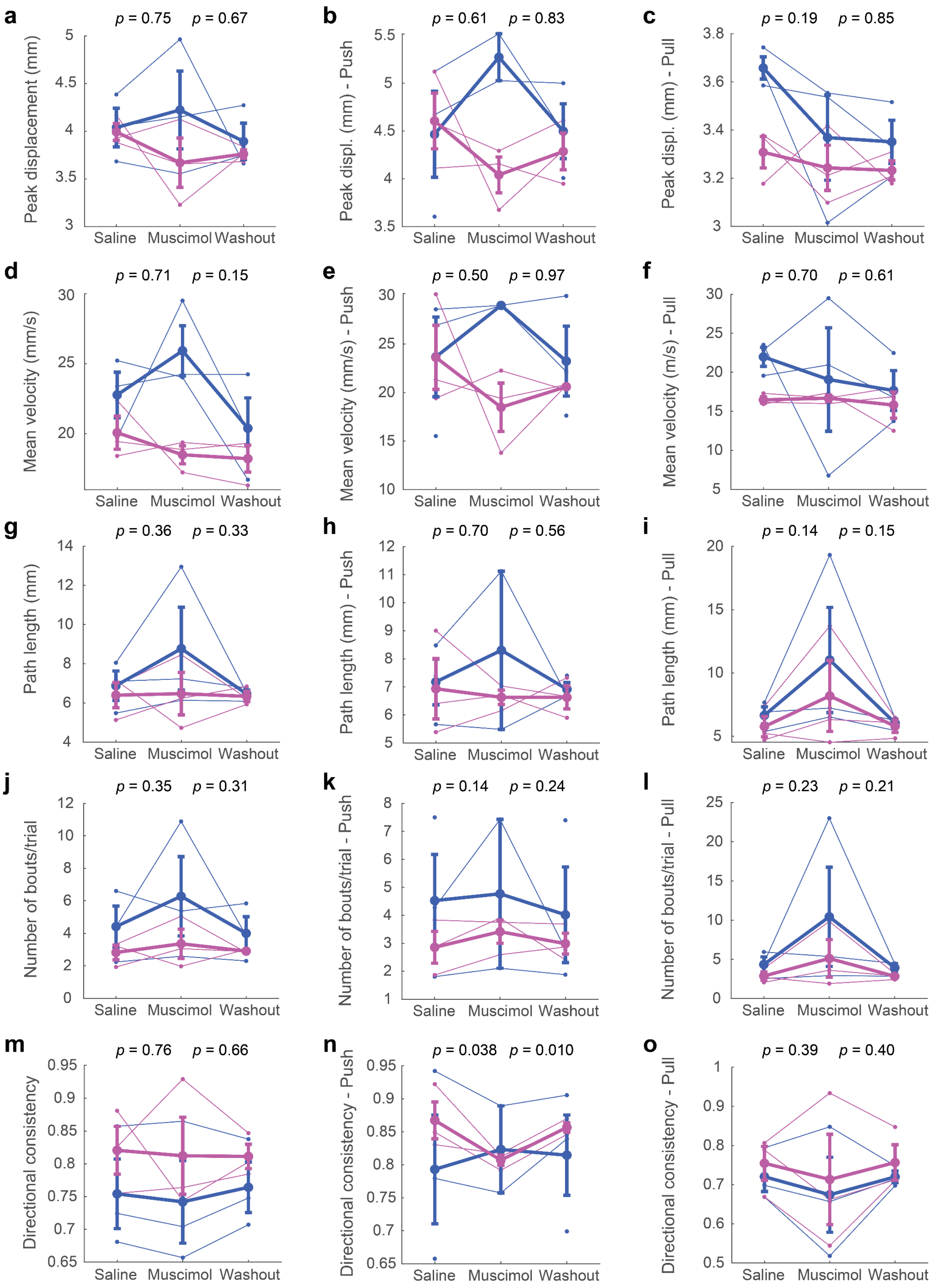


**Figure S2. Muscimol inhibition largely preserves choice kinematics.**

(a-o) The infusion protocol consisted of an initial saline-control session ("Saline"), a muscimol session ("Muscimol"), and a post-muscimol saline-control session ("Washout"). The following measures were compared across sessions: (a-c) peak displacement (mm), (d-f) mean velocity (mm/s), (g-i) path length (mm), (j-l) number of bouts per trial, (m-o) directional consistency. For each measure, results are shown for all choices combined, push choices, and pull choices respectively. Thin lines represent individual animals; thick lines and error bars represent mean ± SEM for males (blue) and females (magenta). The displayed *p*-values are from paired, two-tailed *t*-tests comparing Saline with Muscimol, and Muscimol with Washout.

Peak displacement was the largest joystick displacement from baseline during the decisive bout – that is, the movement bout that crossed choice threshold. Mean velocity was the mean velocity during the decisive bout. Path length was the total distance travelled by the joystick during a single trial. Movement bouts were identified from the magnitude and direction of the smoothed velocity signal, as outlined in the Methods. Number of bouts per trial was the total number of bouts detected within the extracted trial trajectory. Directional consistency was the proportion of bouts whose direction matched that of the decisive bout:

$$\text{Directional consistency} = \frac{n_{\text{bouts in decisive direction}}}{n_{\text{total bouts}}}.$$

Effects of Condition and Sex were assessed using a linear mixed-effects model of the form: Metric ~ Condition * Sex + (1 | animalID), with Condition comprising Saline, Muscimol, and Washout. Significant main effects of Sex were observed for peak displacement on pull choices ($F(1,16.5) = 9.8$, $p = 0.0062$). No significant main effects of Sex were observed for all other metrics (all $p \geq 0.05$). A significant Condition × Sex interaction was observed for peak displacement on push trials ($F(2,9.8) = 4.4$, $p = 0.043$); no significant interactions were observed for the remaining measures (all $p \geq 0.05$). Significant main effects of Condition were observed for directional consistency on push trials ($F(2,10.9) = 5.4$, $p = 0.023$); no significant main effects of condition were observed for the remaining measures (all $p \geq 0.05$).

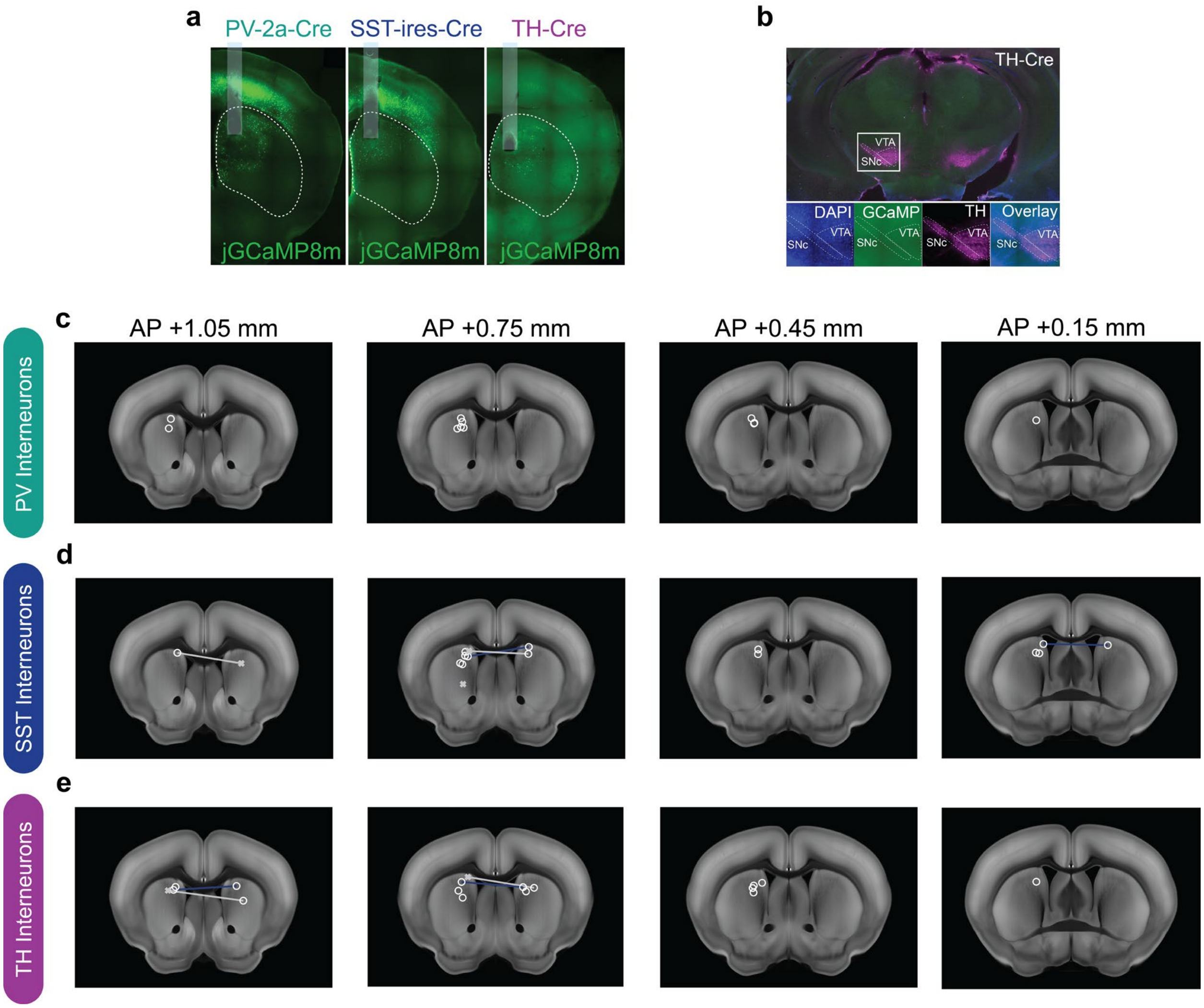


**Figure S3. Dorsomedial striatal interneuron photometry: anatomic targeting.**

(a) Representative histological sections showing viral expression and placement of the fiber optic cannula in the dorsomedial striatum in PV-Cre (N = 11 animals; 5 females, 6 males), SST-Cre (N = 12 animals; 6 females, 6 males), and TH-Cre (N = 13 animals; 5 females, 8 males) animals.

(b) Representative coronal section from a TH-Cre mouse injected in the dorsomedial striatum with a Cre-dependent, jGCaMP8m-encoding adeno-associated virus (AAV9) showing no detectable jGCaMP8m expression in dopamine neurons of the ventral tegmental area (VTA) or substantia nigra pars compacta (SNc). Insets show DAPI (blue), jGCaMP8m (green), tyrosine hydroxylase (TH; magenta), and the merged image.

(c-e) Fiber-tip locations for (c) PV, (d) SST, and (e) TH interneuron photometry cohorts, registered to Allen Mouse Brain Common Coordinate Framework version 3[1] using SHARCQ[2] and shown on coronal sections spaced at 300-μm intervals. White circles indicate included fiber placements, whereas grey crosses indicate placements excluded because of poor signal. Lines connect bilateral fiber-tip locations from the same animal; blue lines indicate animals with two included placements, whereas grey lines indicate animals with one excluded placement.

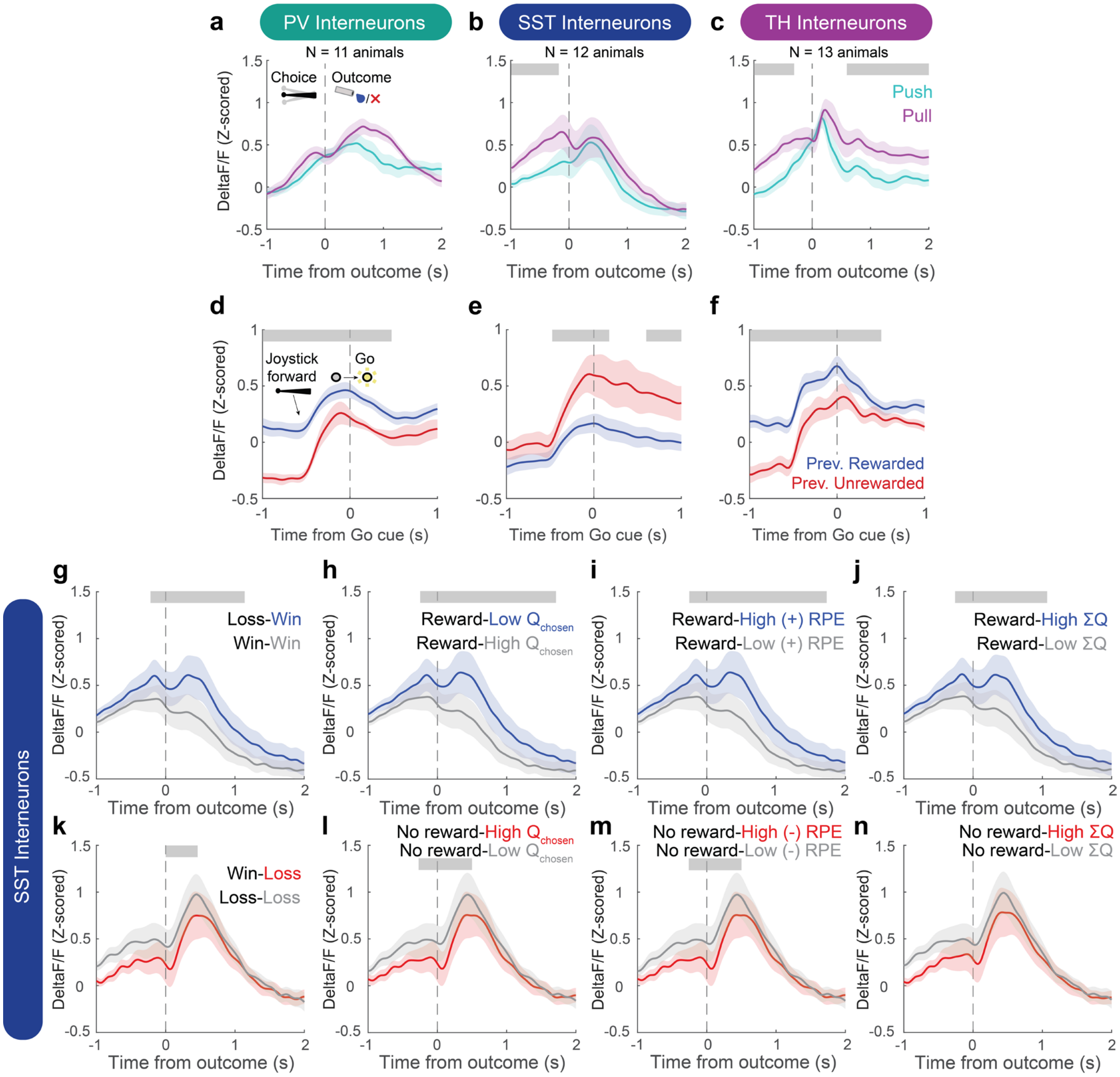


**Figure S4. Dorsomedial striatal interneuron photometry: extended data.**

(a-c) Population photometry responses of (a) PV (N = 11 animals; 5 females, 6 males), (b) SST (N = 12 animals; 6 females, 6 males), and (c) TH (N = 13 animals; 5 females, 8 males) interneurons aligned to outcome and separated according to current trial choice. Cyan and magenta traces indicate push and pull choices, respectively; shaded regions indicate SEM. The vertical dashed line marks outcome onset. Grey bars indicate time periods with significant differences identified using cluster-based permutation testing.

(d-f) Population responses of (d) PV, (e) SST, and (f) TH interneurons aligned to trial start, grouped by previous outcome.

(g-n) Population responses of SST interneurons aligned to outcome and stratified according to reinforcement history and model-derived variables. The top row shows rewarded trials, and the bottom row shows unrewarded trials. Responses are grouped according to (g, k) previous trial outcome, (h, l) the model-estimated value of the currently chosen action ($Q_{chosen}$), (i, m) current-trial reward prediction error (RPE), and (j, n) the summed value of both actions (ΣQ). In (g,k), the first and second terms in each label indicate the previous- and current-trial outcomes, respectively. Values of $Q_{chosen}$ and ΣQ greater than 4 μL were classified as high, and values ≤ 4 μL as low. Within rewarded trials, RPEs > 4 μL were classified as high positive RPEs, and RPEs ≤ 4 μL as low-magnitude positive RPEs. Within unrewarded trials, RPEs < -4 μL were classified as high-magnitude negative RPEs, and RPEs ≥ -4 μL as low-magnitude negative RPEs. Group identities are indicated within each panel; shaded regions indicate SEM. The vertical dashed line marks outcome onset. Grey bars indicate time periods with significant differences identified using cluster-based permutation testing.

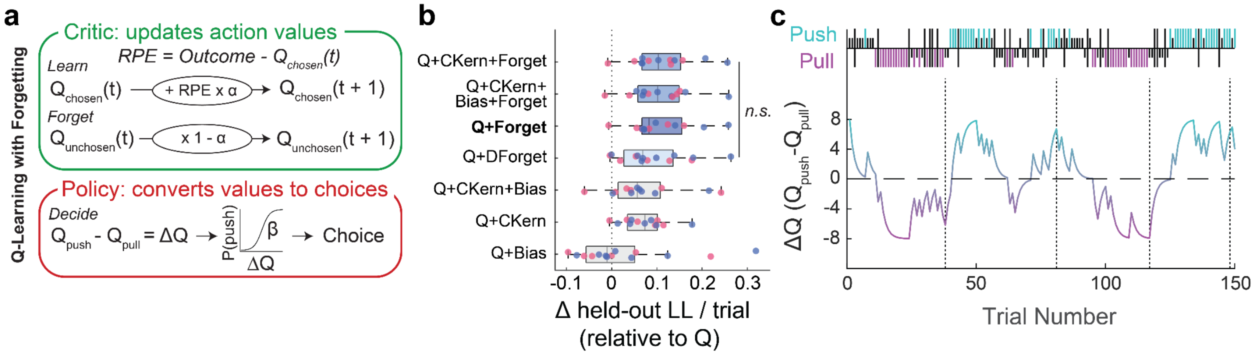


**Figure S5. A Q-learning model with forgetting captures choice behavior in the probabilistic reversal task.**

(a) In the Q-learning model with forgetting, the value of the chosen action was updated after each trial according to its reward prediction error (RPE), scaled by the learning rate (α). The value of the unchosen action decayed toward zero through multiplication by the retention factor (1-α). Choice probability was determined from the difference in action values, $\Delta Q = Q_{push} - Q_{pull}$, using an inverse-temperature parameter β.

(b) Leave-one-session-out cross-validation of candidate Q-learning models incorporating different combinations of an action-bias term (Bias), choice kernels (CKern), and forgetting of the unchosen action. In the Forget model, the forgetting rate was constrained to equal the learning rate $\alpha$; in the differential-forgetting model (DForget), the forgetting rate $\phi$ was estimated independently. Models were fit separately for each animal. For each candidate model and cross-validation iteration, parameters were estimated from all but one session using 25 randomly initialized optimization runs, and the parameter set producing the lowest training negative log-likelihood was retained. These parameters were then held fixed while scoring choices in the omitted session. This procedure was repeated until every session had served once as the test session. Held-out log-likelihoods were summed across sessions for each animal, divided by the total number of held-out trials, and expressed relative to the corresponding score under the standard Q-learning model. The Q-learning model with forgetting outperformed simpler variants, whereas the addition of more complex components did not significantly improve held-out prediction. Displayed $p$-values are from two-sided Wilcoxon signed-rank tests across animals.

(c) Example trial-by-trial evolution of ΔQ, shown beneath the corresponding choice and outcome history. Long colored bars indicate rewarded choices; short black bars indicate unrewarded choices; and long black bars indicate omissions or premature responses before the Go cue. Push is shown in green, and pull in purple.

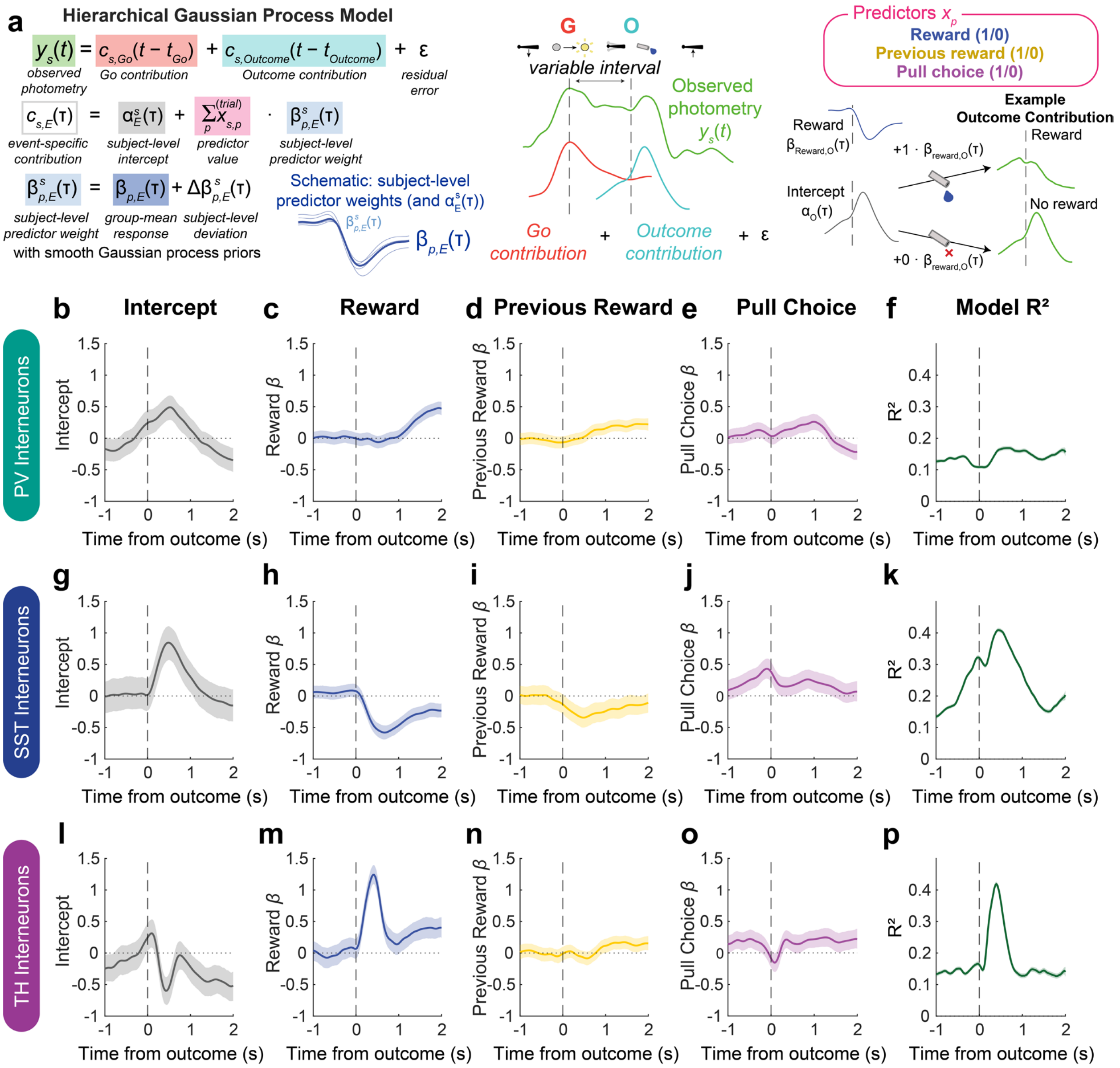


### Figure S6. Hierarchical Gaussian process model reveals differential representation of choice and outcome across striatal interneuron subtypes.

(a) Schematic of the hierarchical Gaussian process model (HGPM) used to analyze photometry signals, following Rinaldi et al. (manuscript in preparation). The photometry signal on each trial was modeled as the sum of Go cue- and Outcome-aligned contributions plus residual error. Each event-aligned contribution comprised a subject-specific intercept function and subject-specific predictor-weight functions multiplied by the corresponding trial-level predictor values. Predictors included current-trial reward (1, rewarded; 0, unrewarded), previous-trial reward (1, previous rewarded; 0, previous unrewarded), and choice (1, pull; 0, push). Smooth Gaussian process priors were placed on the group-level functions and subject-specific deviations, allowing the model to capture both group-level dynamics and variation across mice. The example illustrates how the Outcome-aligned intercept and Reward coefficient combine to predict the Outcome contribution on rewarded and unrewarded trials when the remaining predictors equal zero. At each time point, the magnitude and sign of β describe the association between a predictor and photometry activity, conditional on the other predictors: positive coefficients indicate greater predicted activity when the predictor equals 1, whereas negative coefficients indicate lower predicted activity. Thus, the negative Reward coefficient in SST interneurons indicates lower activity on rewarded than unrewarded trials.

(b-f) Outcome-aligned population-level posterior functions for PV interneurons: (b) intercept, (c) current-trial Reward, (d) Previous Reward, and (e) Pull Choice; together with (f) $R^2$.

(g-k) Corresponding functions for SST interneurons.

(l-p) Corresponding functions for TH interneurons. Intercept functions (b,g,l) represent the estimated Outcome-aligned contribution for the reference condition: an unrewarded current trial following an unrewarded previous trial with a push choice. Solid lines show posterior estimates, and shaded regions indicate 95% credible intervals. $R^2$ values (f,k,p) indicate the proportion of the signal variance explained by the full hierarchical model, including both group-mean and subject-level contributions.

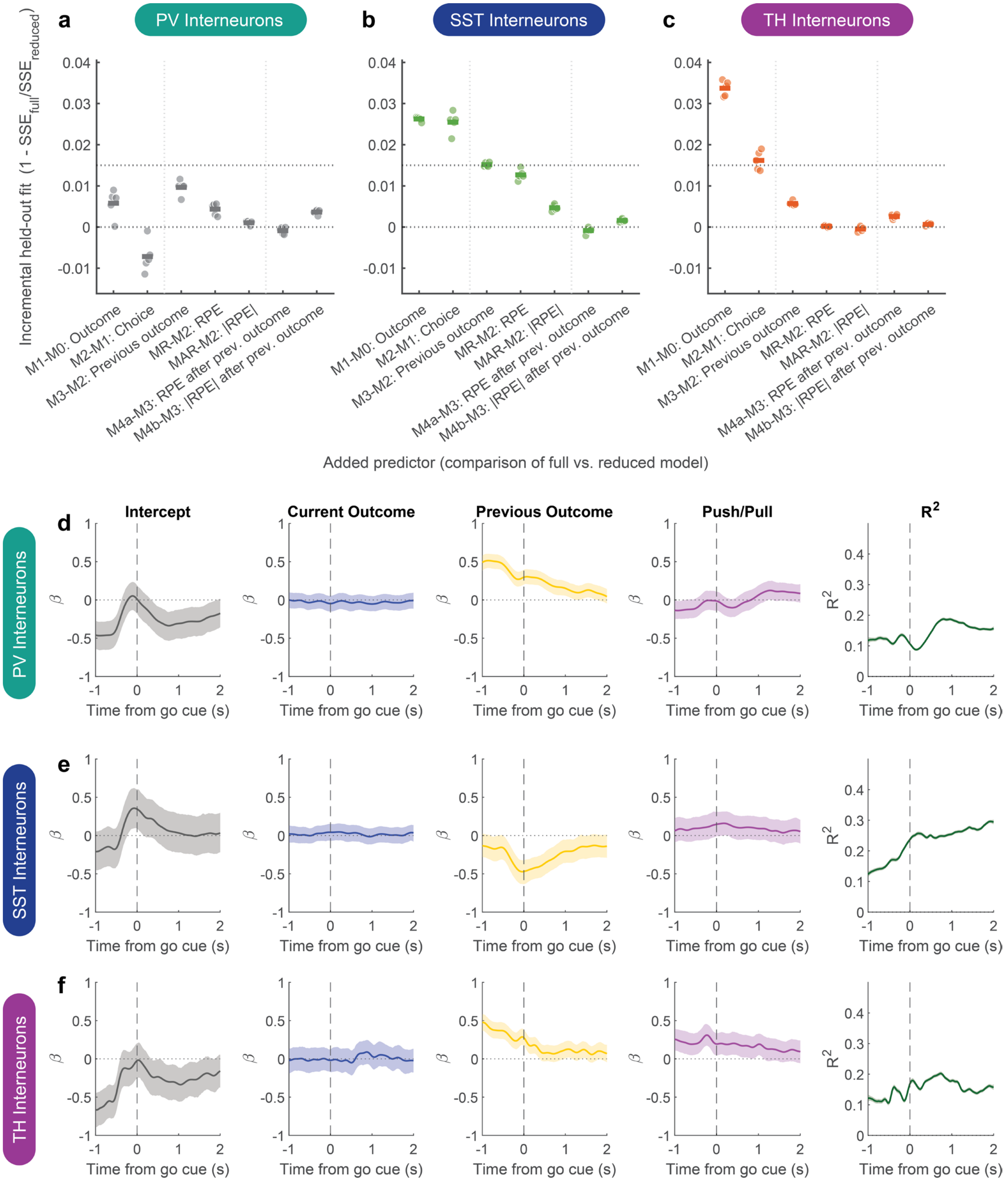


**Figure S7. Predictor selection and Go cue-aligned contributions from the hierarchical Gaussian process model.**

(a–c) Cross-validated improvement in photometry-signal prediction obtained by adding individual task variables, shown for (a) PV, (b) SST, and (c) TH interneurons. Positive values indicate that the added predictor reduced prediction error in held-out animals; values near or below zero indicate little or no improvement.

Predictors were evaluated in three stages. First, we constructed a core task model sequentially by adding current-trial outcome to an intercept-only model (M1–M0), push/pull choice after current outcome (M2–M1), and previous-trial outcome after current outcome and choice (M3–M2). Second, we asked whether signed reward-prediction error (RPE) or absolute RPE (|RPE|) could replace previous-trial outcome as a trial-history predictor, adding each to the model containing current outcome and choice (MR–M2 and M4R–M2). Third, we asked whether either RPE predictor explained additional variance after previous-trial outcome was already included (M4a–M3 and M4b–M3). Vertical dotted lines separate these three sets of comparisons.

Incremental held-out fit was calculated as $1 - SSE_{\mathrm{full}}/SSE_{\mathrm{reduced}}$, where the full model contained the added predictor and the reduced model did not. Models were evaluated using five repetitions of fivefold cross-validation, with animals assigned to folds and predictions for held-out animals generated from population-level fixed effects. Each point represents one cross-validation repetition, and horizontal bars show the mean across repetitions. Within each repetition, prediction errors were pooled across folds and incremental-fit scores were averaged across the prespecified time bins and evaluation windows for that comparison.

Horizontal dotted lines indicate zero improvement and the prespecified mean incremental-fit threshold of 0.015. Because predictors were evaluated over different prespecified time windows, the magnitudes of their composite scores should be compared only descriptively. Final predictor retention was based on the window-specific cross-validation criteria described in the Methods, rather than on these composite scores alone.

(d–f) Go cue-aligned population-level HGPM functions for (d) PV, (e) SST, and (f) TH interneurons. Columns show the intercept and the estimated contributions of current-trial outcome, previous-trial outcome, and push/pull choice, followed by the variance explained by the full model ($R^2$). The intercept represents the estimated signal for the reference condition: a push choice on an unrewarded trial following an unrewarded trial. Solid lines show posterior estimates, and shaded regions indicate 95% credible intervals. $R^2$ includes variance explained by both population- and animal-level model components.

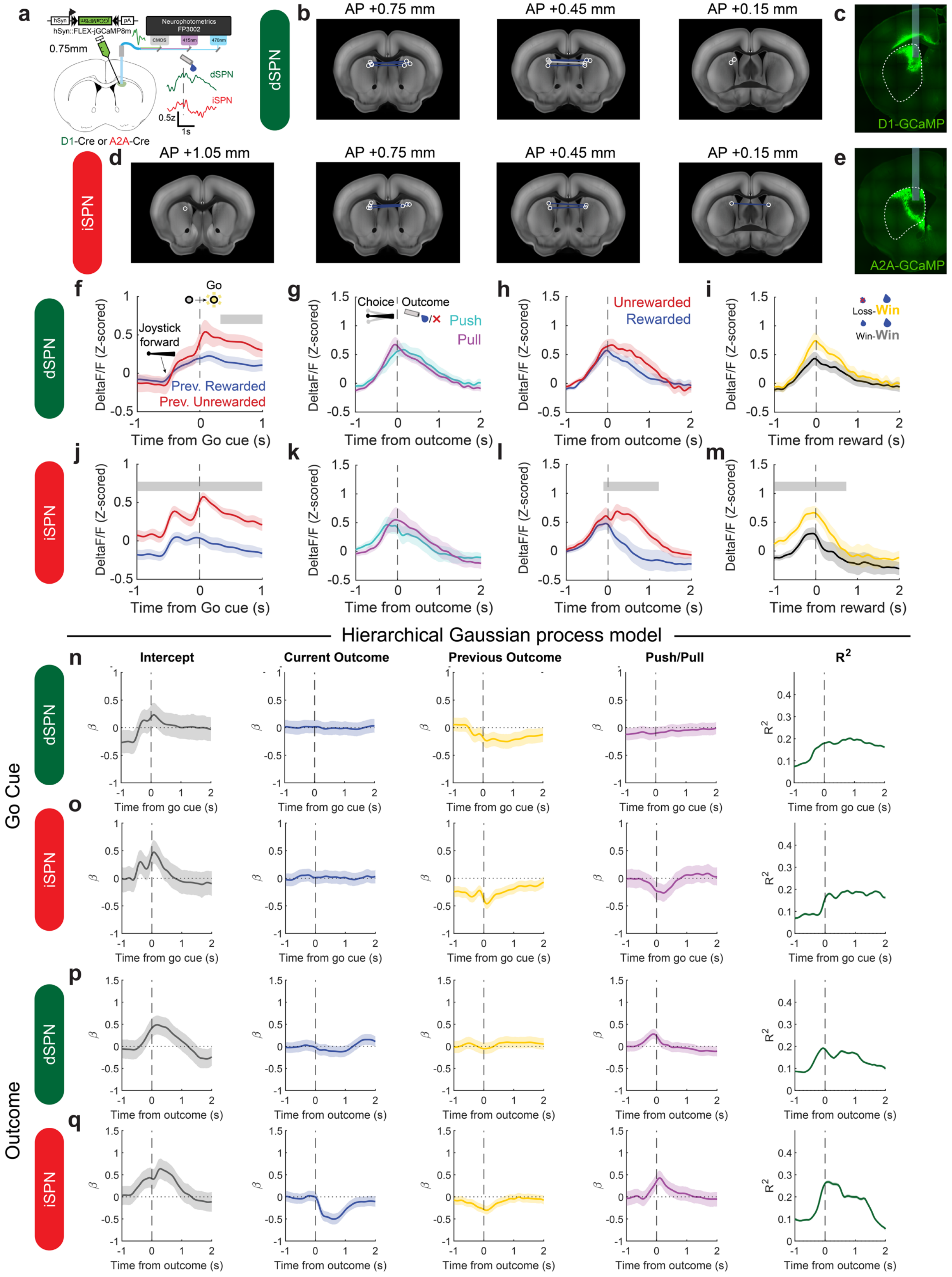
a
hSyn::FLEX-jGCaMP8m
Neurophotometrics FP3002
0.75mm
dSPN
iSPN
0.5z
1s
D1-Cre or A2A-Cre
b
AP +0.75 mm
AP +0.45 mm
AP +0.15 mm
c
D1-GCaMP
d
AP +1.05 mm
AP +0.75 mm
AP +0.45 mm
AP +0.15 mm
e
A2A-GCaMP
f
Go
Joystick forward
Prev. Rewarded
Prev. Unrewarded
DeltaF/F (Z-scored)
Time from Go cue (s)
g
Choice
Outcome
Push
Pull
Time from outcome (s)
h
Unrewarded
Rewarded
Time from outcome (s)
i
Loss-Win
Win-Win
Time from reward (s)
j
k
l
m
Hierarchical Gaussian process model
n
Intercept
Current Outcome
Previous Outcome
Push/Pull
$R^2$
Time from go cue (s)
o
p
q
Time from outcome (s)
Go Cue
Outcome
dSPN
iSPN

**Figure S8. Calcium activity in dorsomedial striatal spiny projection neurons reflects outcomes and reward history.**

(a) jGCaMP8m was expressed in dorsomedial striatal direct- or indirect-pathway spiny projection neurons (dSPNs and iSPNs, respectively), and population activity was recorded during task performance using fiber photometry.

(b,d) Fiber-tip locations for (b) dSPN, and (d) iSPN photometry cohorts, registered to the Allen Mouse Brain Common Coordinate Framework version 3[1] using SHARCQ[2] and shown on coronal sections spaced at 300-μm intervals. White circles indicate included fiber placements, whereas grey crosses indicate placements excluded because of poor signal. Lines connect bilateral fiber-tip locations from the same animal; blue lines indicate animals with two included placements, whereas grey lines indicate animals with one excluded placement.

(c,e) Representative histological sections showing jGCaMP8m expression and fiber-optic cannula placement in the dorsomedial striatum for recordings from (c) dSPNs and (e) iSPNs. Dashed anatomical outline indicates approximate position of caudoputamen.

(f-m) Population photometry responses of dSPNs (N = 10 animals; 5 females, 5 males) and iSPNs (N = 8 animals; 2 females, 6 males). (f,j) Go cue-aligned responses grouped according to the preceding trial outcome. (g,k) Outcome-aligned responses grouped according to current-trial push-pull choice. (h,l) Outcome-aligned responses grouped according to current outcome. (i,m) Reward-aligned responses grouped according to the previous trial outcome. In the Loss-Win and Win-Win labels, the first and second terms indicate the previous- and current-trial outcomes respectively. Solid traces indicate population means, and shaded regions indicate SEM. Vertical dashed lines mark Go-cue or outcome onset. Grey bars indicate time periods with significant differences identified using cluster-based permutation testing.

(n-q) Group-level hierarchical Gaussian process model (HGPM) functions aligned to (n,o) the Go cue- and (p,q) the Outcome for (n,p) dSPNs and (o,q) iSPNs. Functions show the intercept and contributions of current-trial outcome, previous-trial outcome, and push-pull choice, together with time-resolved $R^2$ of the full model. Intercept functions represent the estimated group-level response when the current- and previous-trial outcome and push-pull predictors equal 0. Solid lines show posterior means, and shaded regions indicate 95% credible intervals. $R^2$ indicates the proportion of signal variance explained by the full hierarchical model, including group- and subject-level contributions.

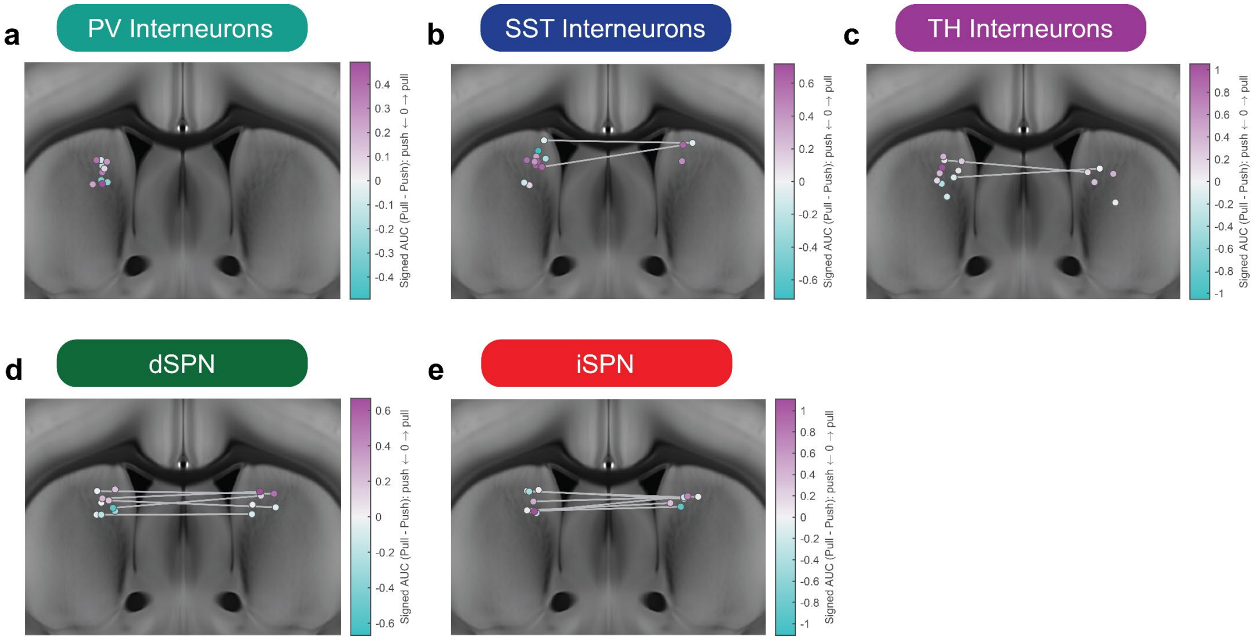


**Figure S9. Anatomical distribution of push-pull choice tuning across dorsomedial striatal cell types.**

(a-e) Anatomical distribution of push-pull choice tuning in (a) PV interneurons, (b) SST interneurons, (c) TH interneurons, (d) direct-pathway spiny projection neurons (dSPNs), and (e) indirect-pathway spiny projection neurons (iSPNs). For each recording site, choice tuning was quantified by trapezoidal integration of the site-specific, outcome-aligned pull-minus-push photometry difference trace from -0.5 to +0.5 s relative to outcome:

$$signed\ AUC = \int_{-0.5\ s}^{0.5\ s} \left[\overline{\Delta FF}_{pull}(t) - \overline{\Delta FF}_{push}(t)\right] dt.$$

Positive values indicate greater activity on pull trials, whereas negative values indicate greater activity on push trials. Each marker represents one recording site and is colored according to the sign and magnitude of the signed AUC. Fiber-tip coordinates were registered to the Allen Mouse Brain Common Coordinate Framework version 3[1] using SHARCQ[2]. Each site's mediolateral and dorsoventral coordinates were retained, whereas anteroposterior coordinates were collapsed onto a representative coronal section at AP +0.45 mm for visualization. Gray lines connect bilateral recording sites from the same animal. Color scales were determined separately for each cell type and therefore should not be compared directly across panels.

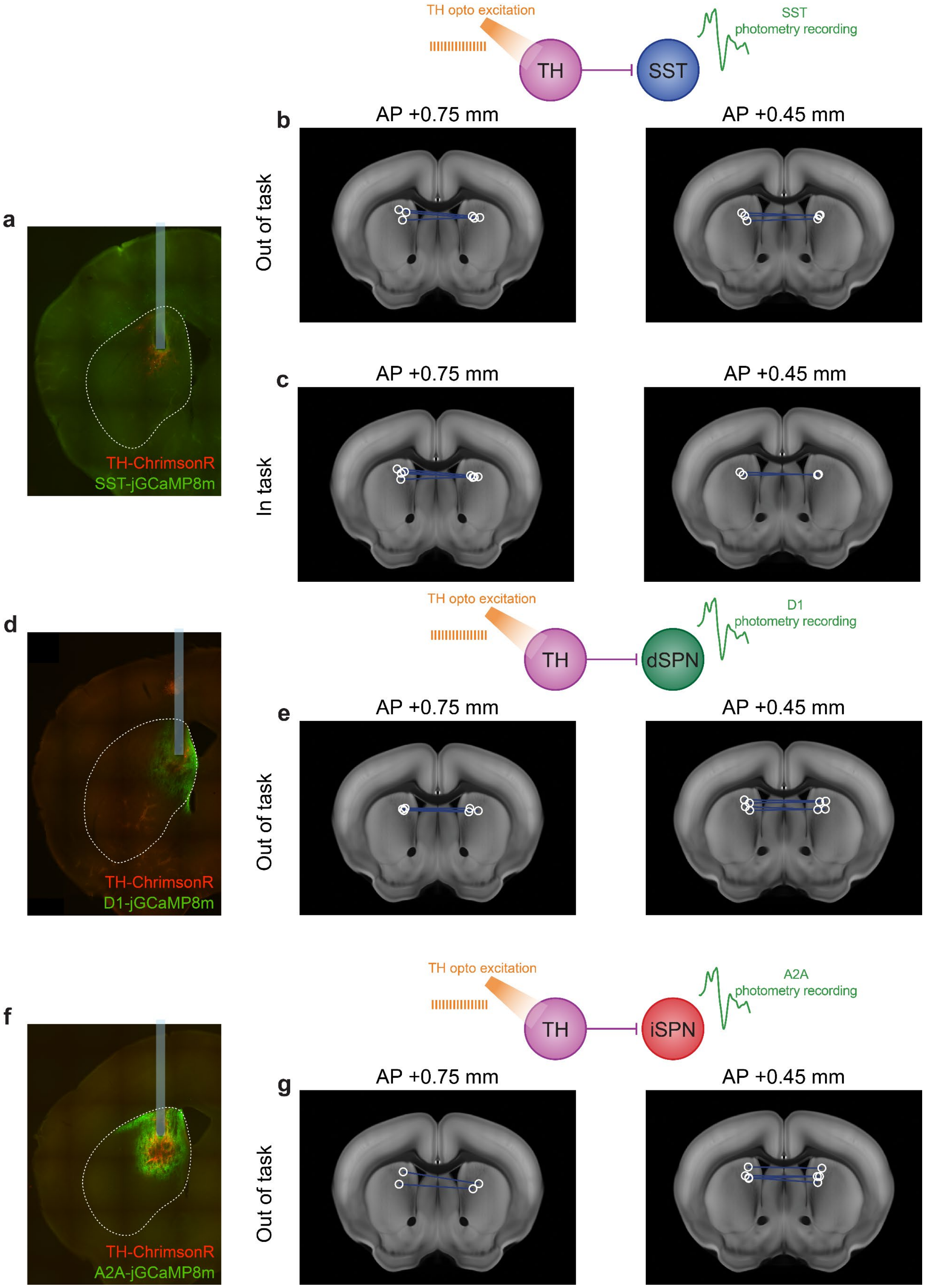


**Figure S10. Histological validation and anatomical targeting for TH-interneuron excitation with simultaneous SST interneuron or SPN photometry.**

(a,d,f) Representative histological section showing TH interneuron ChrimsonR expression (red), cell-type-specific jGCaMP8m expression (green), and fiber-optic cannula placement in the dorsomedial striatum for experiments combining TH-interneuron excitation with photometry recordings from (a) SST interneurons, (d) direct-pathway spiny projection neurons (dSPNs), or (f) indirect-pathway spiny projection neurons (iSPNs). Unlettered schematics illustrate the corresponding optogenetic stimulation and photometry-recording configurations.

(b,c,e,g) Fiber-tip locations for TH interneuron excitation combined with SST interneuron recordings (b) outside the behavioral task or (c) during task performance, and for TH-interneuron excitation combined with (e) dSPN or (g) iSPN recordings outside the task. Locations were registered to the Allen Mouse Brain Common Coordinate Framework version 3[1] using SHARCQ[2] and are shown on coronal sections spaced at 300-μm intervals. White circles indicate included fiber placements, and blue lines connect bilateral fiber-tip locations from the same animal.

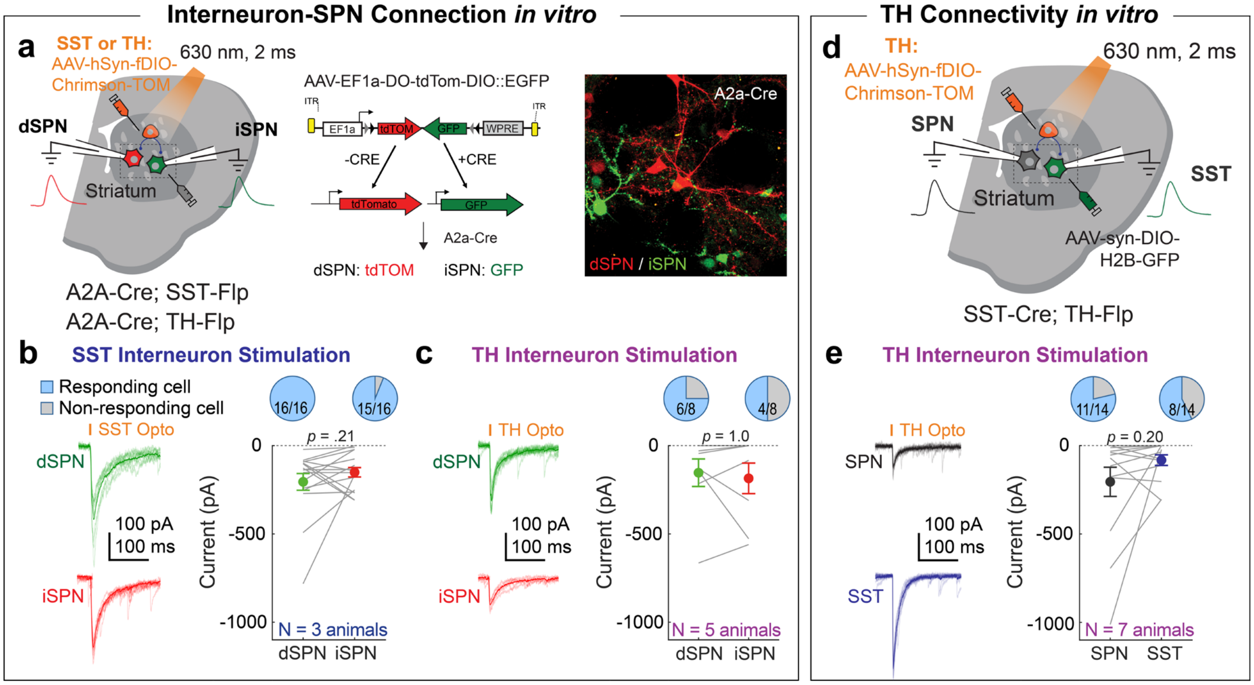


**Figure S11. Local microcircuit connectivity of SST and TH interneurons in vitro.**

(a) Viral strategy used to express ChrimsonR in SST or TH interneurons, tdTomato in direct-pathway spiny projection neurons (dSPNs), and GFP in indirect-pathway SPNs (iSPNs), enabling interneuron stimulation and cell-type specific voltage-clamp recordings from SPNs in vitro.

(b) Representative optogenetically evoked inhibitory postsynaptic currents (oIPSCs) recorded from dSPNs and iSPNs during SST interneuron excitation in 40x field, showing individual sweeps (light) and mean traces (dark). oIPSC amplitudes (cell-level mean ± SEM) did not differ significantly between dSPNs and iSPNs (paired Wilcoxon signed-rank test). Pie charts show the proportions of responsive cells.

(c) As in b, but during TH interneuron stimulation.

(d) Viral strategy used to express ChrimsonR in TH interneurons and GFP in SST interneurons, enabling comparison of TH-evoked responses in SST interneurons and neighboring unlabeled putative SPNs in vitro.

(e) Comparison of TH-evoked oIPSCs in SST interneurons and neighboring putative SPNs.

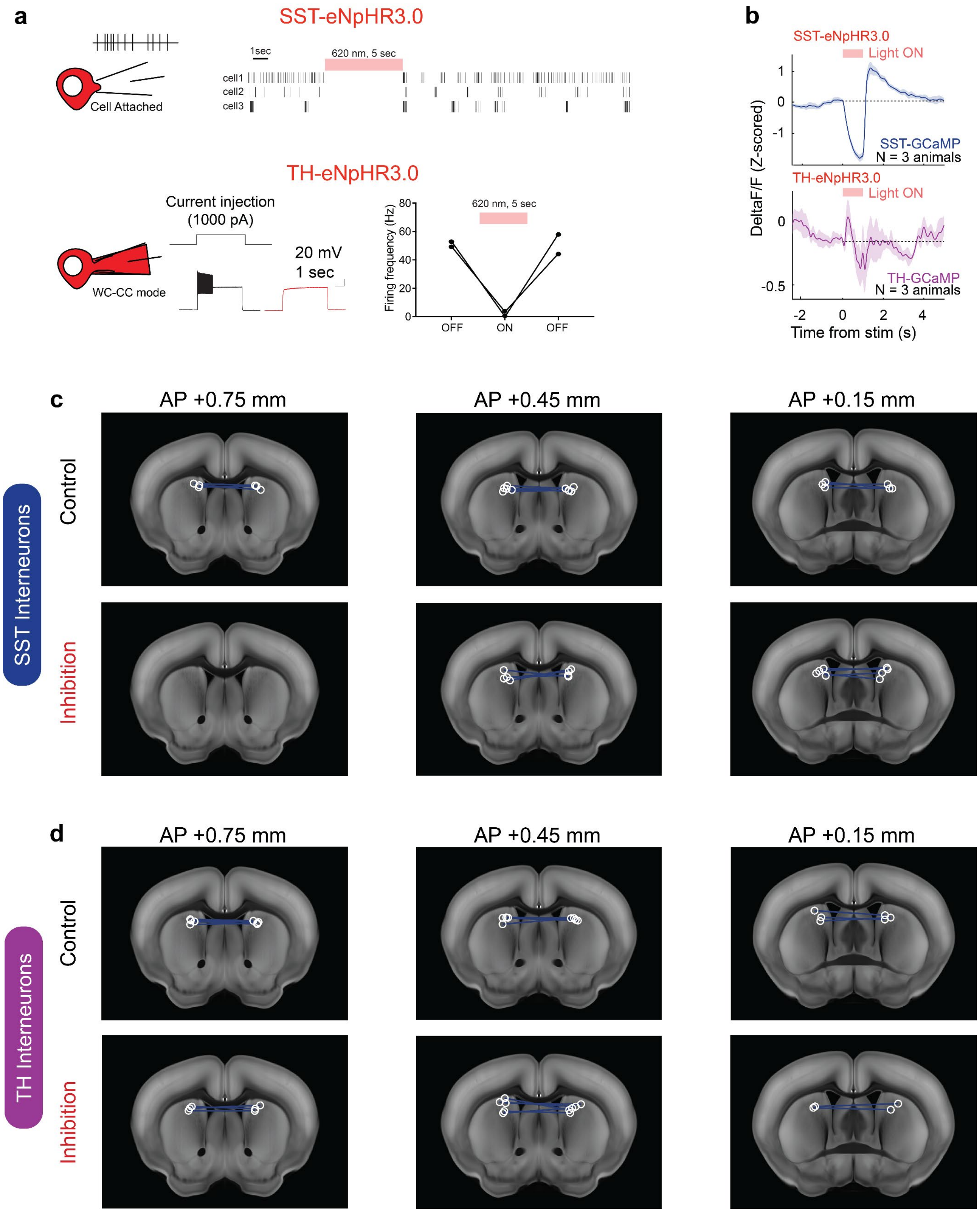


**Figure S12. Validation of eNpHR3.0-mediated inhibition and anatomical targeting for SST and TH interneuron inhibition experiments.**

(a) Electrophysiological validation of the ability of the inhibitory opsin eNpHR3.0 to suppress SST and TH interneuron activity. For SST interneurons (top), three cells from one animal were recorded in the cell-attached configuration. Continuous 620-nm illumination for 5 s suppressed spontaneous spiking, which resumed after illumination ended. For TH interneurons (bottom), two cells from one animal were recorded in whole-cell current-clamp configuration. Spiking evoked by depolarizing current injection (1000 pA) was measured before, during, and after continuous 620-nm illumination. Illumination reduced the evoked firing frequency, which recovered after illumination ended.

(b) In vivo, out of task validation of eNpHR3.0-mediated inhibition using simultaneous fiber photometry and optogenetic stimulation. Continuous 635-nm illumination was delivered for 1 s while calcium-dependent fluorescence was recorded from SST (top: $n$ = 3 animals) or TH (bottom: $n$ = 3 animals) interneurons expressing GCaMP and eNpHR3.0. Traces show mean z-scored $\Delta F/F$, and shaded regions indicate SEM. The red bar indicates the illumination period, and the horizontal dashed line indicates baseline.

(c,d) Fiber-tip locations for (c) SST and (d) TH interneuron cohorts, registered to the Allen Mouse Brain Common Coordinate Framework version 3[1] using SHARCQ[2] and shown on coronal sections spaced at 300-μm intervals. Fluorophore-control cohorts are shown in the top rows, and eNpHR3.0 inhibition cohorts are shown in the bottom rows. White circles indicate included fiber placements, and blue lines connect bilateral placements from the same animal.

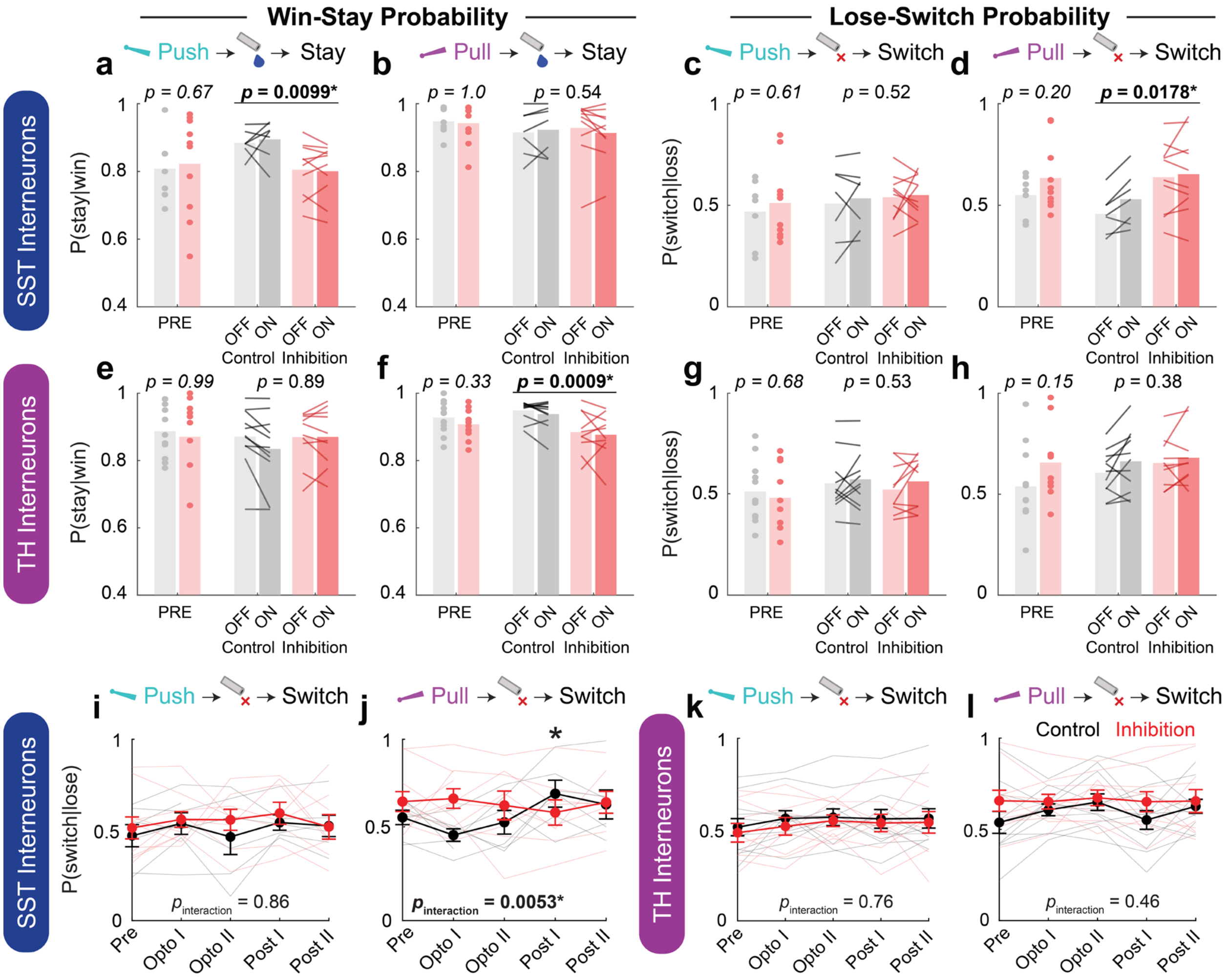


**Figure S13. Repeated outcome-period optogenetic inhibition of dorsomedial striatal SST and TH interneurons produces action-specific behavioral changes that are not time-locked to individual stimulation trials.**

(a-d) Push win-stay (a), pull win-stay (b), push lose-switch (c), and pull lose-switch (d) probabilities in SST-interneuron cohorts.

(e-h) Corresponding measures in TH-interneuron cohorts: Pre bars compare control and inhibition groups before optogenetic sessions. For data pooled across Opto I and Opto II, pale and saturated bars show behavior following light-off and light-on outcome trials, respectively; lines connect light-off and light-on values from the same animal. Bars show group means, and points show individual-animal means. Control and inhibition groups did not differ at baseline on any measure in either the SST cohort (control, *N* = 7 animals; inhibition, *N* = 10 animals) or the TH cohort (control, *N* = 11 animals; inhibition: *N* = 10 animals). Baseline differences were tested using binomial generalized linear mixed-effects models (winStay or loseSwitch ~ condition + (1 | animalID) + (1 | session)). Across Opto I and Opto II, no condition × prevLight interactions were detected for any measure (all $p > 0.05$). Within the inhibition groups, direct contrasts likewise detected no light-on versus light-off differences (all $p > 0.16$), providing no evidence that inhibition during an individual outcome acutely altered the subsequent choice. Optogenetic session data were analyzed using binomial generalized linear mixed-effects models (winStay or loseSwitch ~ condition*prevLight + (1 + prevLight | animalID) + (1 | session)). Significant main effects of condition were observed for push win-stay in the SST cohort ($p = 0.0099$), pull lose-switch in the SST cohort ($p = 0.0178$), and pull win-stay in the TH cohort ($p = 0.0009$).

(i-j) Lose-switch probabilities across experimental phases in SST-interneuron cohorts.

(k-l) Corresponding measures in TH-interneuron cohorts. Thin lines show individual-animal trajectories; thick lines and points show group means ± SEM. For each interneuron subype, trajectories in the control and inhibition groups were compared using generalized linear mixed-effects models (GLMEs; loseSwitch ~ phase*condition + (1 | animalID) + (1 | sessionID)). Marginal tests identified significant phase × condition interactions for SST pull lose-switch ($p = 0.0053$). Relative to Pre, a change in SST pull lose-switch was detected during Post I ($p = 0.018$).

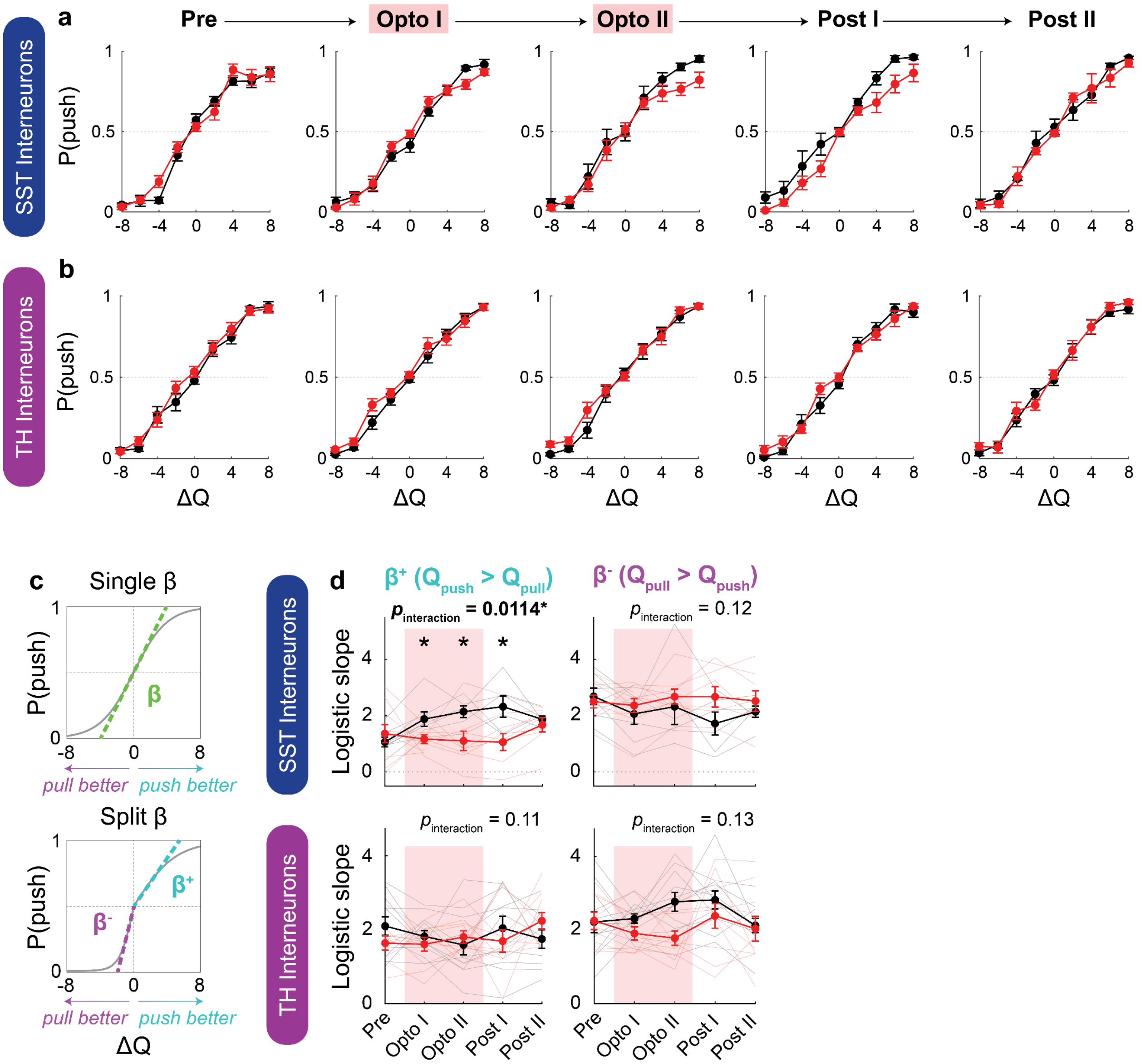


**Figure S14. Optogenetic inhibition of SST and TH interneurons produces no acute trial effects, but dissociable, sustained changes in action policy.**

(a,b) Empirical probability of choosing push as a function of the current-trial push-pull action-value difference ($\Delta Q = Q_{push} - Q_{pull}$), estimated using the Q-learning model with forgetting (Fig S5), during the Pre, Opto I, Opto II, Post I, and Post II phases for (a) SST (control, $N$ = 7 animals; inhibition, $N$ = 10 animals) and (b) TH (control, $N$ = 11 animals; inhibition: $N$ = 10 animals) interneuron cohorts. Trials were grouped into 2-μL ΔQ bins centered from -8 to 8 μL. For each animal, the proportion of push choices was calculated separately within each ΔQ bin and experimental phase; animal-level estimates were included only when at least 10 trials contributed to the corresponding bin. Points and error bars show the mean ± SEM across contributing animals, and lines connect adjacent ΔQ bins. Black indicates fluorophore controls, and red indicates eNpHR3.0-mediated inhibition.

(c) Schematic illustrating how the inverse temperature parameter β controls the steepness of the logistic choice function (top). In the split-β model (bottom), separate inverse-temperature parameters were estimated for trials on which push had the higher estimated value ($\Delta Q > 0$; $\beta^+$) and trials on which pull had the higher estimated value ($\Delta Q < 0$; $\beta^-$). This formulation permits choice-policy stochasticity to differ according to which action has the higher estimated value.

(d) Effects of SST (top) and TH (bottom) interneuron inhibition on $\beta^+$ and $\beta^-$ across experimental phases. Thin lines indicate individual animals; thick lines and error bars indicate mean ± SEM. Black indicates fluorophore controls, red indicates eNpHR3.0 inhibition, and red shading indicates the two optogenetic-inhibition phases. Trajectories in the inhibition and control cohorts were compared using linear mixed-effects models of the form β ~ Phase*Condition + (1 | animalID)). An omnibus F-test from the linear mixed-effects model identified a significant Phase × Condition interaction for $\beta^+$ in the SST cohort ($p = 0.0114$). Relative to the Pre phase, the corresponding Phase × Condition interaction coefficients were significant during Opto I ($p = 0.029$), Opto II ($p = 0.0060$), and Post I ($p = 0.0020$). No significant interactions were observed for SST $\beta^-$ or for either TH parameter (all $p > 0.05$).

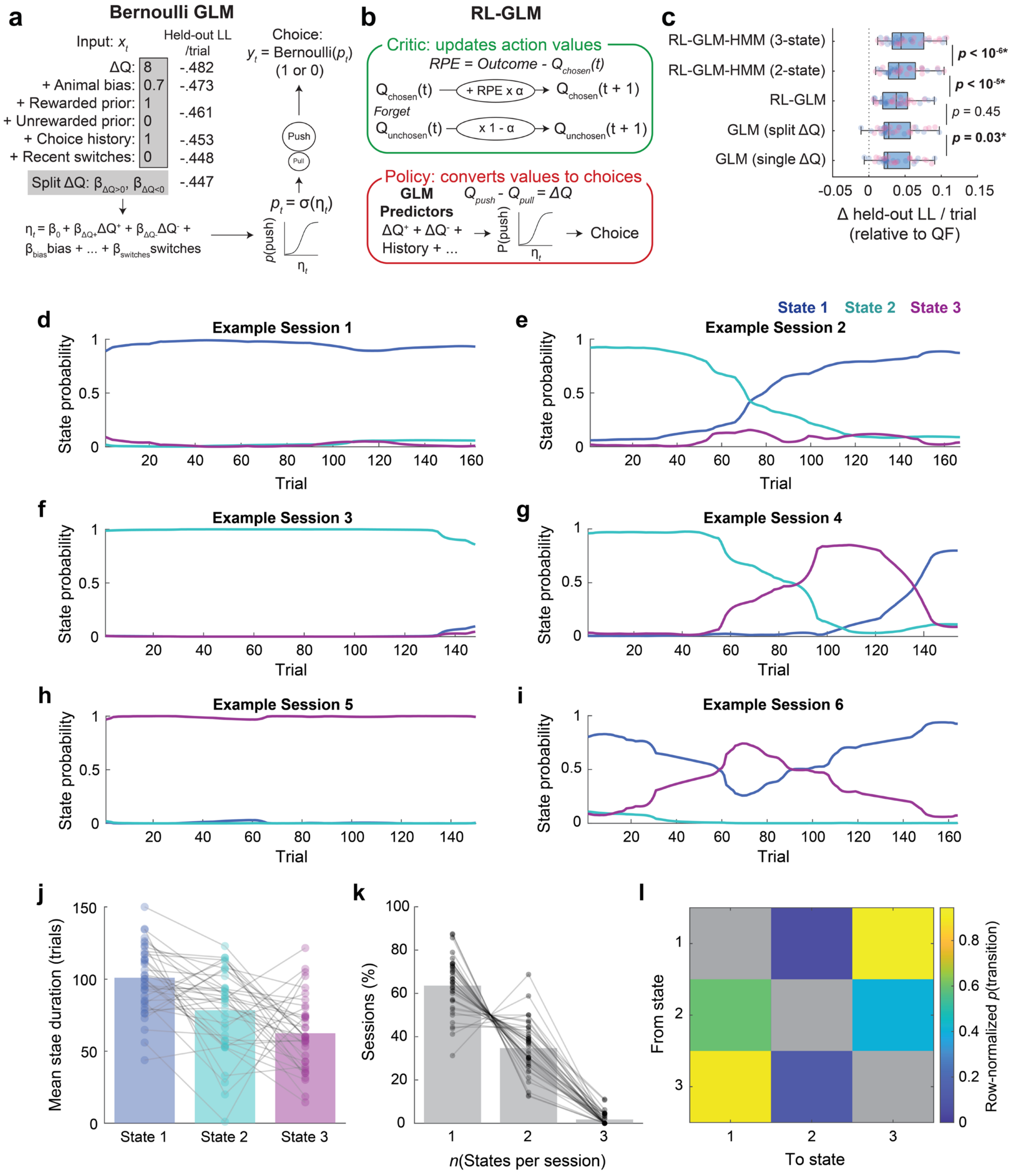


**Figure S15. Three-state reinforcement learning–generalized linear model–hidden Markov model (RL-GLM-HMM): extended analysis of state occupancy and transitions.**

(a) Schematic of the Bernoulli GLM, used to predict trial-by-trial choices. Held-out log-likelihood per trial is shown for models constructed by successively adding predictors.

(b) In the RL-GLM, action values (Q) are updated after each trial according to the reward prediction error (RPE) for the chosen action, scaled by the learning rate (α), while the unchosen action value is multiplied by 1 – α, as in the Q-learning model with forgetting described in Fig S5. Choice probability is determined from a weighted linear combination of GLM predictors, including ΔQ, with a separately estimated coefficient (β) for each predictor.

(c) Five-fold, session-level cross-validation of model variants, including GLMs with a single or split ΔQ coefficient, the RL-GLM, and two- and three-state RL-GLM-HMMs. For each iteration, models were fit at the population level using sessions from four folds. The resulting fixed parameters were then used to score sessions in the held-out fold, with log-likelihood contributions recorded separately for each animal. Held-out log-likelihoods were summed across each animal's sessions and folds, normalized by the animal's total number of held-out trials, and expressed relative to the Q-learning model with forgetting (QF). Pairwise differences between successive models were evaluated using Wilcoxon signed-rank tests; *p* values are shown. Predictive performance increased with addition of latent states, with the three-state RL-GLM-HMM outperforming the two-state model. The RL-GLM did not significantly differ from the GLM using offline-computed Q-values but was retained because it permits estimation of state-specific learning rates.

(d-i) Posterior probabilities of occupying the optimal State 1, push-preferring State 2, and pull-preferring State 3 on each trial of six example sessions, estimated using the RL-GLM-HMM described in the Methods and in Fig 5. Blue, cyan, and magenta indicate States 1, 2, and 3, respectively.

(j) Mean duration, in trials, of contiguous occurrences of each state. Each trial was assigned to the state with the highest posterior probability, and state duration was defined as the number of consecutive trials assigned to the same state. Bars indicate the mean across animals; points represent individual animals, and lines connect values from the same animal. Animals were pooled across the SST control (*n* = 7 animals), SST inhibition (*n* = 10 animals), TH control (*n* = 11 animals) and TH inhibition (*n* = 10 animals) cohorts (*n* = 38 animals total).

(k) Percentage of sessions containing one, two, or three states. Bars indicate the mean across animals; points represent individual animals, and lines connect values from the same animal.

(l) Row-normalized probabilities of transitions between different decoded states. Self-transitions were excluded and are shown in grey. Rows indicate the state being departed, and columns indicate the next state entered. Thus, element ($m$, $n$) represents the conditional probability that State *n* was entered given departure from State *m*:

$$P(S_{t+1} = n \mid S_t = m, S_{t+1} \neq m).$$

Yellow indicates higher and blue indicates lower transition probability.
